# How gradual evolution, fitness landscapes, and trait associations shape mutation rates in large populations

**DOI:** 10.64898/2026.11.19.767823

**Authors:** Arthur Weyna, Ludovic Maisonneuve, Laurent Lehmann

## Abstract

Mutation rates are evolvable traits that respond to selection partly through the associations of modifier alleles with other variants affecting individual fitness. Yet, these variants can be subject to various selective regimes, whose respective role in shaping the evolution of mutation rates remains incompletely understood. To clarify the links between the fitness landscape and selection on mutation rates, we analyse a multi-locus continuum-of-alleles model where traits can affect both individual fitness and the mutation rates expressed across loci. We obtain a closed hierarchical system of moments describing the change in the distribution of traits, and thus in mutation rates, under small trait variation. We find that variation in mutation rates across individuals builds skew in the trait distribution, which plays a key role in the evolution of mutation rates; to track such asymmetries, we go beyond classical normal closure and adapt a cumulant-neglect method. We show that positive mutation rates are favoured by indirect selection out of evolutionary equilibrium (under directional, fluctuating or diversifying selection) and selected against at equilibrium (under homogenising selection). We illustrate these cases with a two-trait model, which we also study using individual-based simulations to evaluate the robustness of our deterministic results to genetic drift. Beyond these examples, our results classify the selective mechanisms acting on mutation rates and rank their importance according to the local shape of the fitness landscape and the variational properties of the population. This provides an account of the different ways selection can act on transmission fidelity in large populations.

**Article summary:** Selection can act on mutation rates indirectly, through its effect on new genetic variants. Here the authors built a mathematical model that summarises the selective mechanisms acting on mutation rates, given the shape of the fitness landscape. They show that higher mutation rates are favoured far from evolutionary equilibrium, under directional, fluctuating or diversifying selection, while lower rates are favoured near equilibrium, under stabilising selection. Individual-based simulations support these conclusions, and explore their limits in the presence of genetic drift. Overall the framework helps explain how selection regimes can shape the diversity of mutation rates across environments and genomic regions.

## 1 Introduction

The rate at which *de novo* genetic mutations appear and are transmitted by an individual to its offspring is influenced by a wide range of evolvable traits (Baer et al. 2007). First, at the molecular level, the mechanisms of DNA replication and maintenance modulate the rate at which genetic modifications appear and persist, both during a cell’s lifetime and during each event of cell duplication (Liu et al. 2016; Chatterjee & Walker 2017; Bębenek & Ziuzia-Graczyk 2018). Because these mechanisms can act at a genome-wide scale (e.g., DNA polymerase) but also interact with local heterogeneity (e.g., DNA conformation, nucleotide composition and epigenetic marks; Monroe et al. 2022), mutation rates may differ and evolve independently in separate regions of the same genome (Hodgkinson & Eyre-Walker 2011; Lang & Murray 2011; Martincorena et al. 2012; Foster et al. 2013; Liu & Samee 2023; Farr et al. 2025; Fyon & Úbeda 2025). Second, at the cellular level, structural characteristics (e.g., cell size, peroxisomes) and cytoplasm content further affect the exposure of genetic material to internal (e.g., free radicals) and external (e.g., radiation) mutagens (Goldman & Travisano 2011; Castenholz & Garcia-Pichel 2012; Terlecky et al. 2012). In multicellular organisms, mutation rates also depend on developmental dynamics, which effectively sort the cell lineages that will be used for propagule production. In particular, the processes of germline segregation and cell death (Otto & Orive 1995; Baer et al. 2007; Aanen & Debets 2019; Lesaffre 2021; Majic et al. 2026) affect the identity and genealogy, and thus the content, of the transmitted genomic copies. Finally, mutation rates are also modulated by the behaviour of organisms, since the many actions taken by individuals directly affect their exposure to mutagens (Castenholz & Garcia-Pichel 2012; Yin et al. 2018), and the timing of their reproduction (Gao et al. 2016). Because mutation rates are shaped by such heritable features, they are most likely subject to the action of natural selection.

The question of the evolutionary forces shaping the diversity of mutation rates has long been addressed, both empirically and theoretically. The first line of reasoning relevant for this question concerns the average fitness effect of mutations. It was understood early and proven empirically that, since mutation is a random process, the majority of mutations occurring in functional genomic regions have a negative impact on fitness (Sturtevant 1937; Eyre-Walker & Keightley 2007). Any genetic variant responsible for increasing mutation rates (i.e., a mutator allele; via any of the mechanisms described above) should thus produce, and become associated with, a higher genetic load (Kimura 1960). As a result, purifying selection alone is expected to favour anti-mutator alleles. From a theoretical perspective, the above argument is usefully framed by studies analysing the evolutionary dynamics of a modifier biallelic genetic locus that controls the mutation probability at one or several linked polymorphic loci affecting fitness (i.e., viability loci; Leigh 1973; Karlin & McGregor 1974; Gillespie 1981a; Holsinger & Feldman 1983; Liberman & Feldman 1986; Kondrashov 1995; Dawson 1998; Dawson 1999; Liberman et al. 2016; Altenberg et al. 2017). These studies provided mathematical proofs that in large populations, when viability loci have reached a selection-mutation equilibrium (such that mutation produces less fit alleles on average), indirect selection (i.e., selection via associations with selected loci) always disfavours mutator alleles and brings mutation rates ever closer to zero.

Since all observed mutation rates are strictly positive, additional evolutionary mechanisms have been proposed that would offset the effect of mutation load. Some authors have proposed functional explanations, stating that mutation rates should be bounded by the physical constraints of biochemistry, and that decreasing these rates any further would be too costly for the organism (Kondrashov 1995; Dawson 1998; Sniegowski et al. 2000; Tannenbaum et al. 2003) due to trade-offs between transmission fidelity and other life-history aspects (Avila & Lehmann 2023). Other authors have pointed out that random changes in mutation rates are expected to increase rather than decrease them (i.e., the distribution of mutational effects on mutation rates is biased towards higher mutation rates; Drake 1993). Both the above mechanisms promote the maintenance of positive mutation rates. In addition, since they are direct, these mechanisms can overpower indirect selection for reduced mutation rates when mutation rates are low. This leads to the general expectation that mutation rates should converge towards stable internal equilibrium values. A key finding is then that these equilibrium values should be higher in smaller populations, due to the reduced relative strength of indirect selection under genetic drift (Lynch 2010; Martincorena & Luscombe 2013; Gervais & Roze 2017). This is consistent with large-scale empirical evidence showing a strong negative correlation between proxies of mutation rates and effective population size across the tree of life (Lynch et al. 2016; Bergeron et al. 2023).

The idea that the evolution of mutation rates is governed primarily by a balance between direct selection due to a cost of fidelity and indirect selection due to genetic load induced by deleterious mutations does not fit well, however, with a number of empirical observations. It does not, for instance, fully explain why mutator genotypes can be selected for in stressful or fluctuating environments, in both prokaryotes (Bjedov et al. 2003; Loh et al. 2010; Elgrail et al. 2024; Barnett et al. 2025) and eukaryotes (Lamb et al. 2008; Raynes et al. 2011). It also fails to explain strong variation in mutation rates across regions of the same genome, which (despite some local variation in effective population size) mostly share a common population-level context (Hodgkinson & Eyre-Walker 2011; Martina et al. 2012; Foster et al. 2013; Monroe et al. 2022; Farr et al. 2025; Fyon & Úbeda 2025). To understand these observations, it is useful to turn to theoretical studies that have modelled evolutionary dynamics at a mutation-modifier locus in contexts where the affected viability loci have not reached an equilibrium between purifying selection and mutation (e.g., due to environmental change or frequency-dependent selection; Eshel 1973; Painter 1975; Gillespie 1981b; Ishii et al. 1989; Taddei et al. 1997a; Taddei et al. 1997b; Kessler & Levine 1998; Johnson 1999a; Johnson 1999b; Tenaillon et al. 1999; Travis & Travis 2002; Tanaka et al. 2003; André & Godelle 2006; Allen & Rosenbloom 2012; Carja et al. 2014; Rosenbloom & Allen 2014; Good & Desai 2016; Shen et al. 2020; Mintz & Fu 2022; Ferrare & Good 2024; see also Levins 1967). In these models, the most common viability alleles at any given time are not necessarily the best ones. When the best alleles are rare, mutator alleles are disproportionately responsible for introducing favourable variants, and can thus become associated with the latter. As a result, they can increase in frequency via genetic hitch-hiking, even if new mutations still have a deleterious effect over one generation on average. Via this mechanism of indirect positive selection, and depending on the specific ecological scenario, higher mutation rates can be promoted in the short term (during adaptation to a new environment) or in the long term (under some forms of environmental fluctuations).

Theoretical results show that several selective mechanisms act on mutation rates, and that their relative strength depends on the selection regime (the type and strength of the active selective mechanisms). Nonetheless, the idea that differences in selection regimes could explain part of the global diversity of mutation rates remains under-explored (Lynch et al. 2016). Current models also rarely disentangle the different effects of direct selection (e.g., costs) and indirect selection (i.e., via associations with viability loci) on the evolutionary dynamics of mutation modifiers, and on the dynamics of their association with viability loci (see Johnson 1999b; Gervais & Roze 2017; Fyon & Úbeda 2025 for meaningful exceptions). It would thus be valuable to study the interplay of direct and indirect selection under arbitrary selective pressures, such that the respective effects of different selection regimes can be compared. Understanding the evolutionary dynamics of mutation rates during transient environmental change (i.e., change that cannot be described as a long-term distribution of environmental states), when viability loci experience bursts of positive selection, would also be relevant.

In this article, we address the above theoretical challenges by studying the evolution of mutation rates conceived as functions of quantitative traits. The essence of our approach is that we do not model the evolutionary dynamics of a single modifier locus with global effects. Instead, we describe the evolutionary dynamics of a vector of quantitative traits, each encoded by a separate locus with its own local mutation rate and an infinite number of possible alleles, and each affecting fitness and any mutation rate in arbitrary ways. That is, each trait can be a viability trait and/or a mutation-modifying trait (or modifier trait for short). This allows for a quantitative description of the evolution of mutation rates by natural selection, through the change in the trait distribution, summarised in its moments (i.e., trait averages and their associations; Bürger 2000; Walsh & Lynch 2018). This approach has three relevant features. First, it leads to a partitioning of the respective roles of direct and indirect selection in the evolution of mutation rates. Second, it is valid for many different fitness functions, such that the selection pressures acting on the population at any given point can be described and classified using the local shape of that function (Phillips & Arnold 1989), including social and ecological feedbacks. Third, it allows for intragenomic heterogeneity in mutation rates. With this representation, we obtain an understanding of how and when natural selection can act on global and local mutation rates. To illustrate our results, we apply our model to specific evolutionary scenarios involving the coevolution of two traits (a viability and a modifier trait). We check our results against individual-based simulations of the same scenarios, which lets us assess the role of genetic drift.

## 2 The model

### 2.1 Set-up

#### 2.1.1 Biological assumptions

We consider an infinitely large well-mixed population of haploid organisms reproducing with non-overlapping generations. Every individual in this population carries a single chromosome with *n*_*x*_ linked loci, with each locus encoding a quantitatively varying trait value following the continuum-of-alleles model (Kimura 1965; Fleming 1979; Iwasa et al. 1991; Bürger 1998; Waxman 2003; Otto & Day 2011). We denote by ***x*** = (*x*_1_, …, *x*_*n*_ ) the trait vector, where *x*_*i*_ *∈* ℝ is the *i*^th^ trait value encoded by the *i*^th^ locus (see fig. 1), and by ∫*p*_*t*_(***x***) = *n*_*t*_(***x***)*/N*_*t*_ the frequency density of that vector in the population at time *t*. This depends on the density *n*_*t*_(***x***) of individuals with trait ***x*** at *t* and on the total density *N*_*t*_ = ∫*n*_*t*_(***x***)d***x*** of individuals in the population (where *p*_*t*_(***x***)d***x*** = 1). Throughout, integrals are taken over all reals, and integration over a vector is used as a shorthand for integration over all of its elements (i.e., 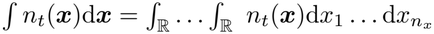).

**Fig. 1:**
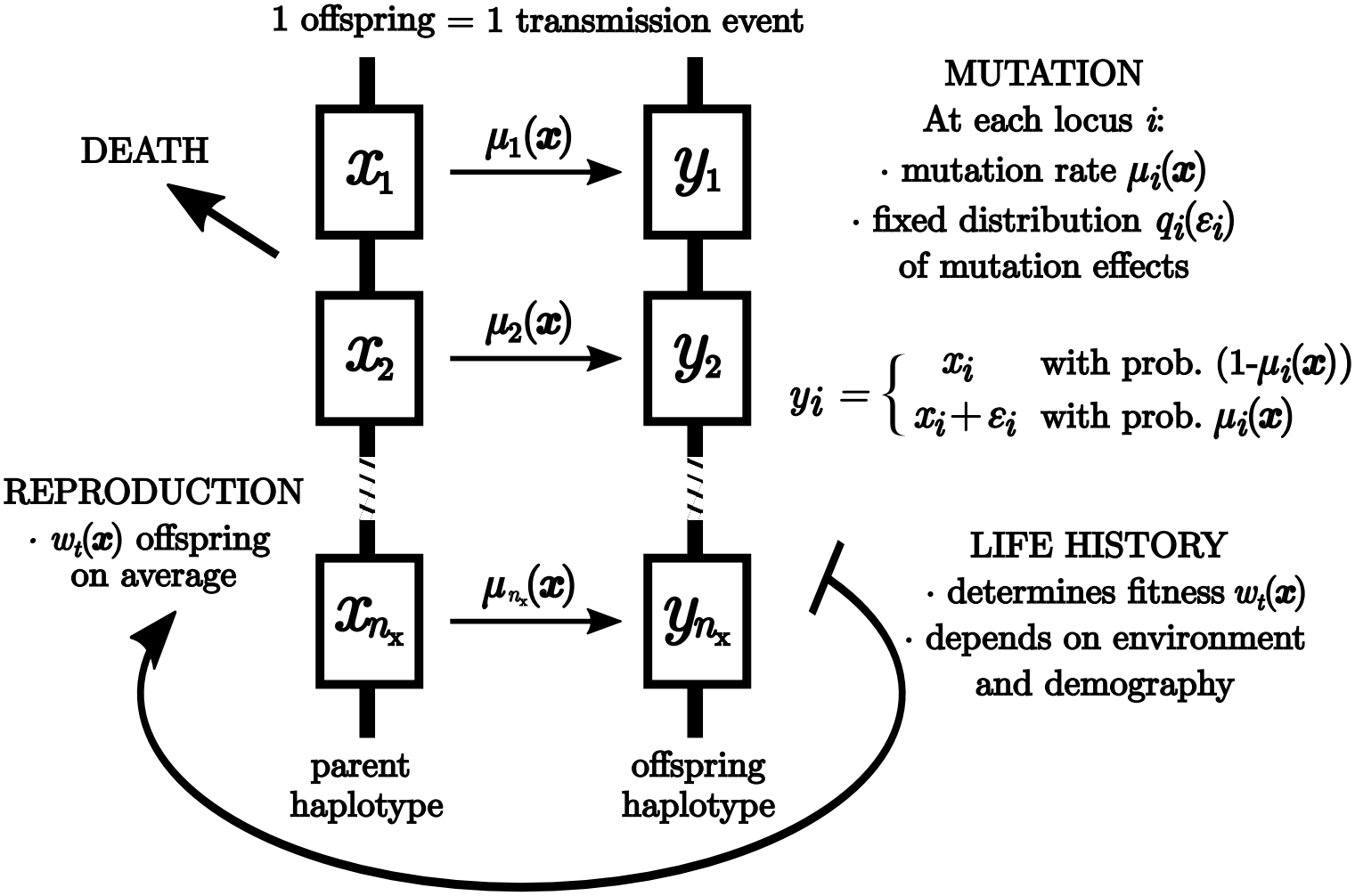
Life cycle assumptions. The diagram summarises the model of section 2. The vectors 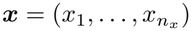 and 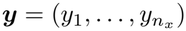 collect the trait values of a focal individual in generation *t* and of one of its offspring in generation *t* + 1, respectively. Each trait is encoded by a separate locus. Trait values affect life history (survival, reproduction…) and the mutation rate at each locus. Mutation effects are independent across loci and change parental trait values by a small amount. **Alt text:** Diagram of one generation of the model. A parent carrying a vector of trait values produces offspring that inherit its trait values, except that each offspring can carry a small new mutation at each locus, with a probability that depends on the parent’s trait values.

Each generation, individuals produce offspring clonally and die. The expected number of viable descendants that a focal individual with traits ***x*** leaves in generation *t* + 1, i.e., its individual fitness, is denoted *w*_*t*_(***x, ϕ***_*t*_(***x***)) and assumed differentiable with respect to all its arguments (as are all functions throughout). The trait values ***x*** of an individual can affect its fitness both directly and by changing its interactions with the rest of the population. We capture these interactions through the vector 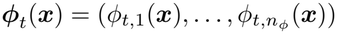, collecting *n*_*ϕ*_ regulating factors of the form

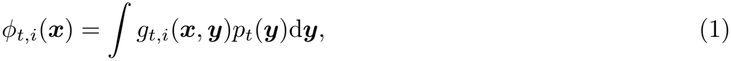

where *g*_*t,i*_(***x, y***) is a regulating kernel giving the contribution of individuals with traits ***y*** to the *i*^th^ regulating factor. Possible regulating factors include population density (obtained by setting *g*_*t,i*_(***x, y***) = *n*_*t*_(***x***)*/p*_*t*_(***x***)), the average of some trait *k* in the population (obtained by setting *g*_*t,i*_(***x, y***) = *y*_*k*_), or more complicated statistics of the population. Note that we use a subscript *t* in *w*_*t*_ and *g*_*t,i*_ to indicate that these functions can vary from one time step to the next due to exogenously changing environmental conditions (e.g., precipitation, temperature).

The traits of a focal individual also determine the probability that each of its offspring inherits *de novo* mutations. Specifically, we assume that mutations occur independently at each locus, with a maximum of one mutation per locus per reproduction event, and denote by *µ*_*i*_(***x***) the mutation probability at locus *i*. The effect size of new mutations on traits is independent of ***x***, and we write *q*_*i*_(*ζ*_*i*_) for the probability density that a mutant offspring’s *i*^th^ trait value differs from that of its parent by an amount *ζ*_*i*_. Throughout, we assume that mutation effects are symmetric on each trait, i.e., that *q*_*i*_(*ζ*_*i*_) = *q*_*i*_(*−ζ*_*i*_) for all *i*, such that 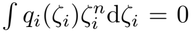 for any odd positive integer *n*. With this, the probability density that an offspring inherits trait value *x*_*i*_ + *ζ*_*i*_ given that its parent has traits ***x*** is

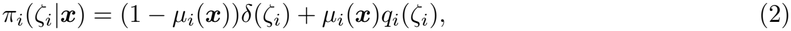

where *δ*(*ζ*_*i*_) denotes the Dirac delta function (which is zero everywhere except at *ζ*_*i*_ = 0 and satisfies *∫δ*(*ζ*_*i*_)d*ζ*_*i*_ = 1). The first term in eq. (2) accounts for the probability that the offspring does not carry a *de novo* mutation at locus *i* (and thus inherits trait value *x*_*i*_), while the second term accounts for the probability that the offspring carries a mutation with effect *ζ*_*i*_, and therefore we have ∫*π*_*i*_(*ζ*_*i*_|***x***)d*ζ*_*i*_ = 1. Finally, because the occurrence and effect of mutations across loci are independent, the probability density that an individual inherits traits ***x*** + ***ζ*** (where 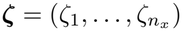) given that its parent had traits ***x*** is

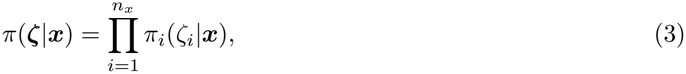

and satisfies ∫*π*(***ζ***|***x***)d***ζ*** = 1.

### 2.2 Evolutionary analysis

#### 2.2.1 Dynamics of the trait distribution

To track the changes in the distribution of traits due to the combined effects of selection and mutation over one generation, we start by describing the change in the density *n*_*t*_(***x***) of individuals with traits ***x***. Since the descendants of these individuals at generation *t* + 1 have density *n*_*t*_(***x***)*w*_*t*_(***x, ϕ***_*t*_(***x***)), and since the probability density that these descendants have traits ***y*** = ***x*** + ***ζ*** is *π*(***ζ***|***x***) = *π*(***y*** *−* ***x***|***x***), the density 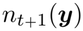 of individuals with traits ***y*** at generation *t* + 1 is

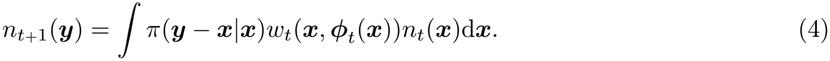

Eq. (4) describes how trait density changes over one generation as the result of the joint effects of reproduction and mutation in the parental generation. Integrating eq. (4) over all traits ***y*** gives

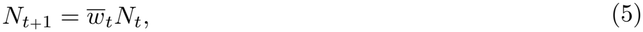

where 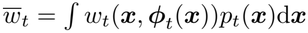 is the average individual fitness at generation *t*, showing that changes in the total population density do not depend directly on mutation. Dividing eq. (4) by 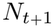 yields

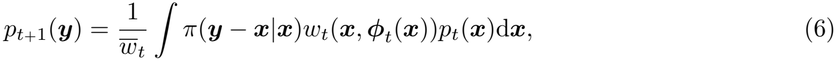

which is the main recursion we use to study the change in the distribution of traits.

#### 2.2.2 Dynamics of moments and closure assumptions

Directly tracking change in the full distribution of traits (determined by eq. 6) is not feasible analytically. It is also costly numerically and rarely allows for direct biological interpretations of the evolutionary mechanisms at work. To gain more insight, a common strategy is to approximate the evolutionary process by following the trait distribution through a finite number of its moments, which provide summary statistics of the distribution’s position and shape (e.g., Bulmer 1971; Barton & Turelli 1987; Frank & Slatkin 1990; Bürger 1993; Bürger 2000; Walsh & Lynch 2018; Lion et al. 2023). In doing so, the choice of which moments to track depends on which aspects of the trait distribution are expected to evolve and to matter for a particular question.

Here, to follow the overall change in mutation probabilities, we first want to track the position of the population in trait space, given by the vector of average traits 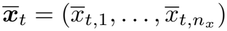, where

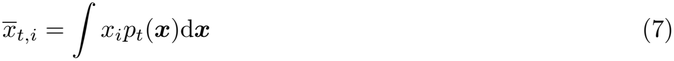

is the population average of trait *i* at generation *t*. To do so, we aim to express the change 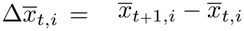 in that average over one generation.

Because we are interested in the evolution of mutation probabilities, which are expected to affect trait variation, we also want to follow trait (co)variances (i.e., second-order central moments), defined using 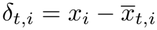 as

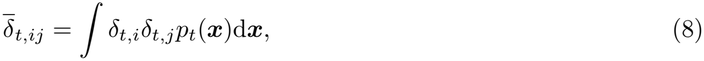

for all traits *i* and *j*. To achieve this, we seek an expression for the change 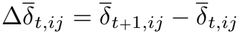 in the (co)variance 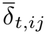 over one generation.

When there is variation in mutation probabilities within the population (i.e., due to variation in traits), parts of it can generate trait variation at a faster pace, which can lead to asymmetries in the trait distribution (i.e., the distribution becomes “fatter” on one side). Since they may affect relevant selective effects, we also need to track these asymmetries via third-order central moments, namely trait (co)skewnesses, defined as

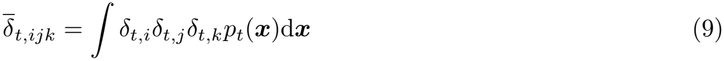

for all traits *i, j* and *k*. With *i* = *j* = *k*, the skewness 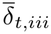 carries information about asymmetry in the distribution of trait *i*; when it is positive (negative), this distribution has a larger upper (lower) tail. With *i≠ j* = *k*, the two-way coskewness 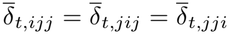 tells us whether trait *i* is associated with variation in trait *j*; when it is positive (negative), there is more variation in trait *j* among individuals with larger (smaller) values of trait *i*, but also, equivalently, more positive (negative) covariation between traits *i* and *j* among individuals with larger values of trait *j*. Finally, with *i≠ j≠ k* all distinct, the three-way coskewness 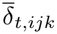 measures association between traits *i, j* and *k*; when it is positive (negative), higher values for two of these traits are found among individuals with larger (smaller) values of the third. To track these associations, we aim to express the change 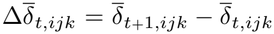 in the (co)skewness 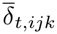.

Since describing the evolutionary dynamics including third-order moments is already involved and captures a number of phenomena (see below), neglecting higher-order moments offers a useful balance between tractability and accuracy. Therefore, our aim is to understand the coevolution of modifier and viability traits, by focusing on the evolutionary dynamics of first-, second- and third-order moments described above. To achieve this, we start by deriving the moment-generating function of the offspring trait distribution in terms of that of the parent trait distribution (see appendix A.1), and use it to express dynamical equations for 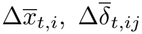 and 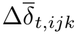 for any *i, j* and *k* (appendix A.2). Because these equations depend on the full trait distribution however, we make two additional assumptions to close the system of moments, i.e., to express the change in moments of order up to three in terms of these moments themselves.

First, we assume that the trait distribution is narrow, that is *δ*_*t,i*_ = *O*(*ϵ*) where *ϵ* is a small scalar. With this, individual fitness and mutation probabilities can be Taylor expanded about the vector of average traits (see appendix B.1) and change in the trait moments expressed as a sum of terms with a hierarchy of scales, meaning that terms involving higher-order moments are generally orders of magnitude smaller. This allows us to approximate the change in moments by its lower-order (i.e., leading) terms. Since the leading-order selection terms for the change in moments (generally, second, third and fourth order for 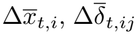, and 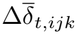, respectively) can vanish at evolutionary attracting fitness extrema, however, we also include terms of the next order (third, fourth and fifth order, respectively) which describe selection at such extrema. We assume that the latter never vanish at the same time as leading order terms, so that terms of even higher order (fourth, fifth and sixth order onward, respectively) never significantly affect the dynamics of trait moments. Note that when sorting terms by order, we also assume that mutation effects are of the same size as trait deviations (i.e., *ζ*_*i*_ = *O*(*ϵ*)).

While the leading-order truncation described above greatly simplifies moment dynamics, it does not by itself close the system of moments at third order. The system still depends on moments of order four and five (on which the dynamics of (co)variances and (co)skewnesses depend), themselves changing according to higher moments. To achieve full moment closure, we adopt the cumulant-neglect moment-closure method (see Wojtkiewicz et al. 1996 and Gillespie 2009 for general descriptions), which uses the relationship between moments and cumulants to approximate moments of order four and five as functions of moments of order three and less. This cumulant-neglect assumption states that cumulants of order four and five are small enough to be neglected. We describe this method and the resulting approximations in detail in appendix B.2.

#### 2.2.3 Road map of the results

Our main goal is to understand how modifier traits, and thus mutation probabilities, evolve over time under different selection regimes. As such traits can evolve both through direct selection (when they affect fitness) and through indirect selection (via genetic associations with other traits under selection), this requires that we understand how the trait moments of interest in the population change over time. This change involves several selective effects that can interact in different ways with varied consequences for mutation probabilities, and we thus proceed in two steps. We thus first describe the many components affecting the change in trait moments in the general case, where each trait may affect both mutation probabilities and individual fitness (sections 3.1–3.3). This account is descriptive, and does not immediately make obvious what mechanisms act on the mutation probabilities themselves. Once all evolutionary components are laid out, we examine their respective strengths in different contexts (section 3.4) and how their effects combine to shape the evolution of mutation probabilities under different selection regimes and biological settings (section 3.5).

## 3 Results

### 3.1 Change in averages

We show in appendix B.3.1 that, to leading orders, the change 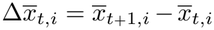 in the average of trait *i* over one generation can be partitioned as

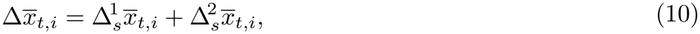

into a change 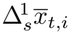 (of order 2 in trait deviations) due to first-order (directional) selection and a change 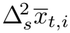 (of order 3) due to second-order (quadratic and correlational) selection.

The change due to directional selection is

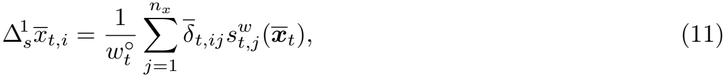

where 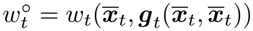, with 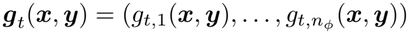 the vector of regulating kernels, and

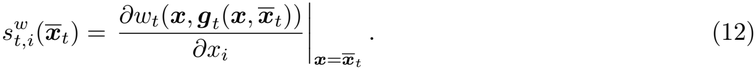

This is the derivative of an individual’s fitness with respect to its *i*^th^ trait value *x*_*i*_ evaluated in a population monomorphic for the average traits, and informs us about the slope in the local fitness landscape. Positive (negative) values of 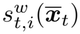 indicate that individuals with higher (lower) values of trait *i* have higher fitness on average, such that directional selection favours an increase (decrease) in that trait’s average. A point in trait space where 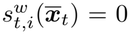 for all *i* is referred to as a singular trait. The fact that the derivative 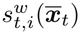 is enough to predict the effect of directional selection, despite fitness depending on the whole trait distribution via the regulating factors, is a consequence of assuming small trait variation (see appendix B.1.1). Due to that same assumption, only the leading-order term 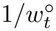 of the normalisation factor 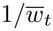 (originating from eq. 6) remains in eq. (11), and higher-order terms can be neglected (see appendix B.1.1). This holds for all forthcoming equations.

Eq. (11) has the same form as results derived from polygenic quantitative genetics models (Lande 1979, eq. 7; Lande & Arnold 1983, eq. 6; Phillips & Arnold 1989, eq. 1) and under the continuum-of-alleles model (Iwasa et al. 1991, eq. A6; Débarre et al. 2014, eq. 4; Mullon & Lehmann 2019, eq. 3). It shows that average traits can change due to both direct directional selection (i.e., the component of eq. 11 with *i* = *j*) and indirect directional selection (i.e., the components of eq. 11 with *i≠ j*; see tab. S1). Direct directional selection acts on the average of trait *i* when this trait varies (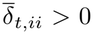; see fig. 2A). Indirect directional selection, on the other hand, acts on the average of trait *i* when this trait covaries with another trait 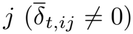, because individuals with more favourable values of trait *j* then also possess higher or lower values of trait *i* (see fig. 2B).

**Fig. 2:**
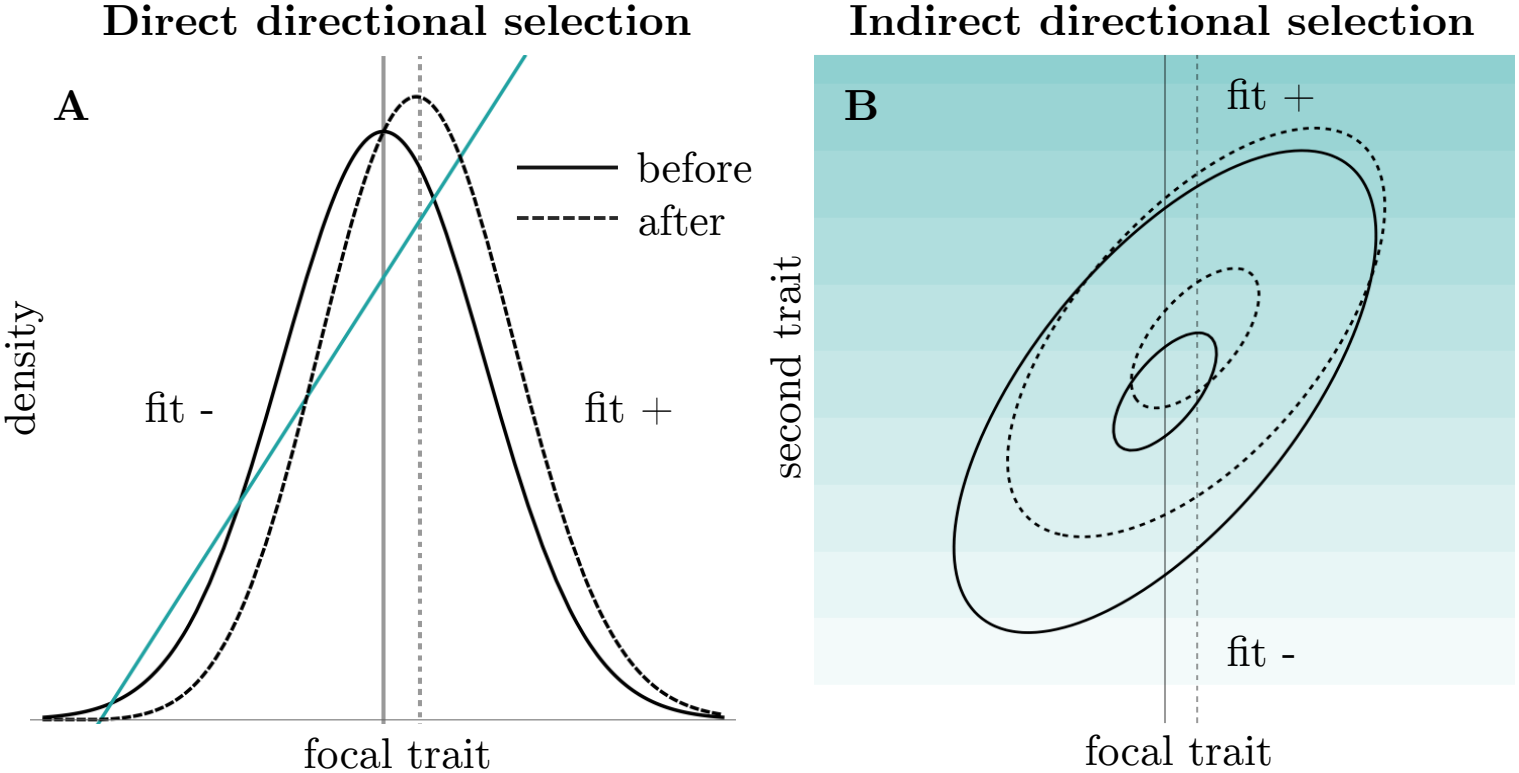
Change in the mean due to directional selection alone. Each panel shows the change in the trait distribution over one generation, obtained numerically from eq. (6) without mutation; vertical lines give the means before and after selection. **A**. Viability trait under direct directional selection (blue line: local fitness landscape). **B**. Neutral trait under indirect directional selection, when a second trait (vertical axis) is under directional selection and the two traits are associated. In **B**, the background colour shows the local fitness landscape (darker is fitter); larger and smaller ellipses circumscribe 80 and 20 percent of the distributions, respectively. The code and numerical values for this figure are in Supplementary Mathematica Notebook 1. **Alt text:** Panel A shows a trait distribution on a fitness curve, with its mean after selection shifted relative to its mean before selection. Panel B shows ellipses for a two-trait distribution on a shaded fitness landscape, with the mean of the neutral trait shifting because it is associated with the selected trait.

Change in the average of trait *i* can also be due to second-order selection, through the term

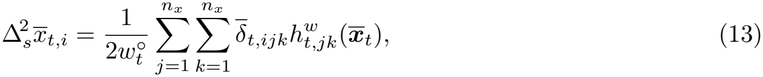

where

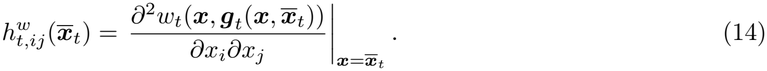

This second-order selection coefficient informs us about curvature in the local fitness landscape. With *i* = *j*, positive or negative values of 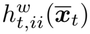 indicate that individuals with extreme values of trait *i* have higher or lower fitness, respectively, than average. In that case we say that quadratic selection is diversifying or homogenising, respectively. In turn, with *i≠ j*, positive (negative) values of 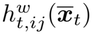 indicate that individuals with positive (negative) combinations of traits *i* and *j* have higher fitness on average, such that correlational selection favours these combinations. Since eq. (13) contains third-order moments as factors, it vanishes when the trait distribution is symmetric (e.g., under normality). Analogous expressions can thus only be found in general descriptions of change in trait moments (Barton & Turelli 1987; Bürger 1991; Rice 2002; Rice 2020). When asymmetries in the trait distribution are taken into account, average traits can change due to both direct second-order selection (i.e., the components of eq. 13 with *i* = *j* or *i* = *k*) and indirect second-order selection (i.e., the components of eq. 13 with *i≠ j* and *i≠ k*; see tab. S1).

Direct second-order selection on the average of trait *i* can be categorised into direct quadratic selection (the element of eq. 13 with *i* = *j* = *k*) and direct correlational selection (the elements with *i* = *j≠ k* or *i* = *k≠ j*; see tab. S1). Direct quadratic selection acts when the distribution of trait *i* is skewed (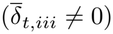), such that it possesses a longer tail that can be amplified (or trimmed) by diversifying (or homogenising) selection (fig. 3A). In turn, direct correlational selection acts when there is more covariation between two co-selected traits *i* and *j* among individuals with larger (or smaller) values of trait *i* ( 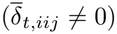 ), so that the trait combinations favoured by correlational selection contain an excess of large (or small) values of trait *i* (see fig. 3B).

**Fig. 3:**
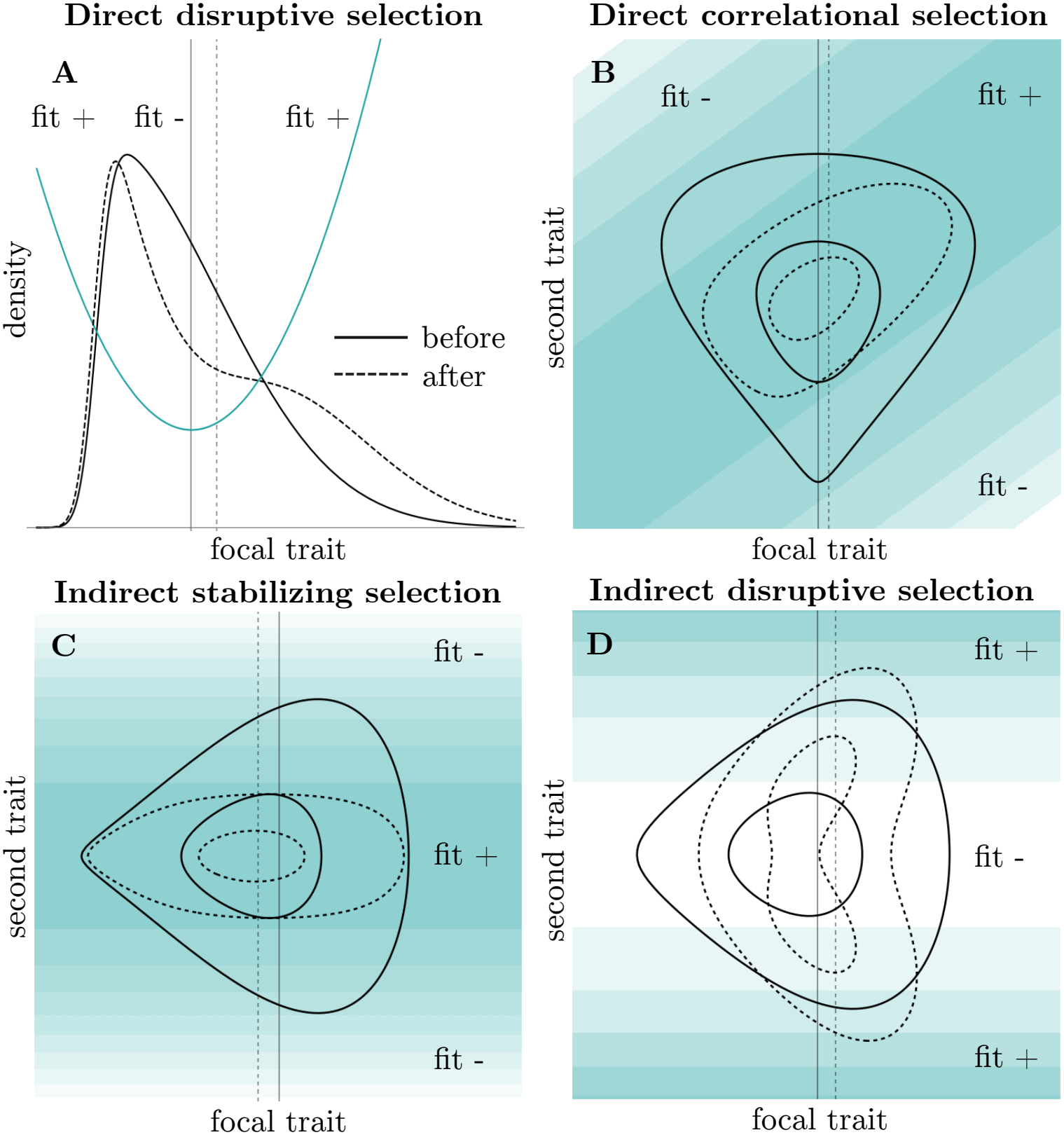
Change in the mean due to quadratic and correlational selection alone. Each panel shows the change in the trait distribution over one generation, obtained numerically from eq. (6) without mutation or directional selection; vertical lines give the means before and after selection. **A**. Viability trait under direct diversifying selection (blue line: local fitness land-scape). With the same positively skewed starting distribution, homogenising selection would trim the upper tail and decrease the mean (see main text). **B**. Direct positive correlational selection: the favoured trait combinations contain an excess of large focal values, increasing the focal mean. **C**. Indirect homogenising selection (average values at a second trait are favoured), which corresponds to the reduction principle (see main text). **D**. Indirect diversifying selection (extreme values at a second trait are favoured). In **B–D**, the background colour shows the local fitness landscape (darker is fitter); larger and smaller ellipses circumscribe 80 and 20 percent of the distributions, respectively. The code and numerical values for this figure are in Supplementary Mathematica Notebook 1. **Alt text:** Panel A shows a skewed trait distribution on a fitness curve, whose long upper tail is amplified so that its mean increases. Panels B to D show ellipses for two-trait distributions on shaded fitness landscapes, with the mean of the focal trait shifting under correlational selection (B), homogenising selection on a second trait (C) and diversifying selection on a second trait (D).

Likewise, indirect second-order selection on the average of trait *i* can be categorised into indirect quadratic selection (the element of eq. 13 with *i≠ j* = *k*) and indirect correlational selection (the elements with *i≠ j≠ k* all distinct; see tab. S1). Indirect quadratic selection acts when trait *i* is associated with variation in trait 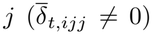, while trait *j* experiences quadratic selection 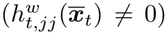. In that case, selection favours individuals with average (or extreme) values of trait *j*, which contain an excess of small (or large) values of trait *i*, leading to a decrease (or increase) in the latter trait’s average (see fig. 3C and 3D). In turn, indirect correlational selection acts when trait *i* is associated with different levels of covariation between two traits *j* and *k* (i.e., if 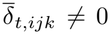; see tab. S1) under correlational selection 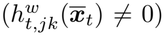. Then, selection favours associations between traits *j* and *k* that are found more often within individuals with larger (or smaller) values of trait *i*, leading to an increase (or decrease) in that trait’s average.

### 3.2 Change in variances and covariances

We now turn to the change in trait variance and covariances. We show in appendix B.3.2 that, to leading orders, the change 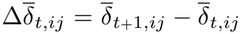 in the (co)variance between traits *i* and *j* over one generation can be partitioned as

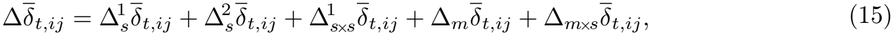

into the change 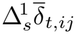 (of order 3) due to directional selection, the change 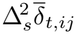 (order 4) due to second-order selection, the change 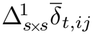 (order 4) due to change in average traits, the change 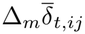 (order 2) due to mutation alone, and the change 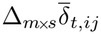 (order 4) due to the interaction of mutation and selection.

The change in the (co)variance between traits *i* and *j* due to directional selection is

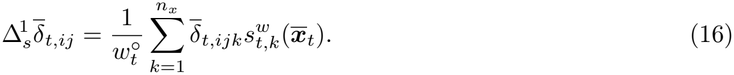

Since it contains third-order moments as factors, eq. (16) vanishes when the distribution of traits is symmetric, and analogous terms can only be found in studies that do not assume normality (Barton & Turelli 1987; Bürger 1991; Rice 2002; Gilpin & Feldman 2019; Rice 2020). This equation shows that, just like average traits, trait (co)variances can change due to both direct directional selection (i.e., the component of eq. 16 with *i* = *k* or *j* = *k*) and indirect directional selection (i.e., the components of eq. 16 with *i≠ k* and *j≠ k*; see tab. S1).

The variance in trait *i*, 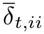, can change due to direct directional selection when the distribution of that trait is skewed 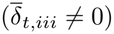, such that it comprises a larger tail that can be trimmed or amplified, leading respectively to a decrease or an increase in variance (see fig. 4A). In addition, 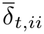 can change due to indirect directional selection if variation at that trait is associated with another trait 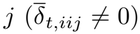 itself under directional selection 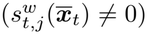. In this case, the individuals favoured by selection have higher (or smaller) levels of variation in trait *i*, resulting in a change in 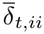 (see fig. 4B).

**Fig. 4:**
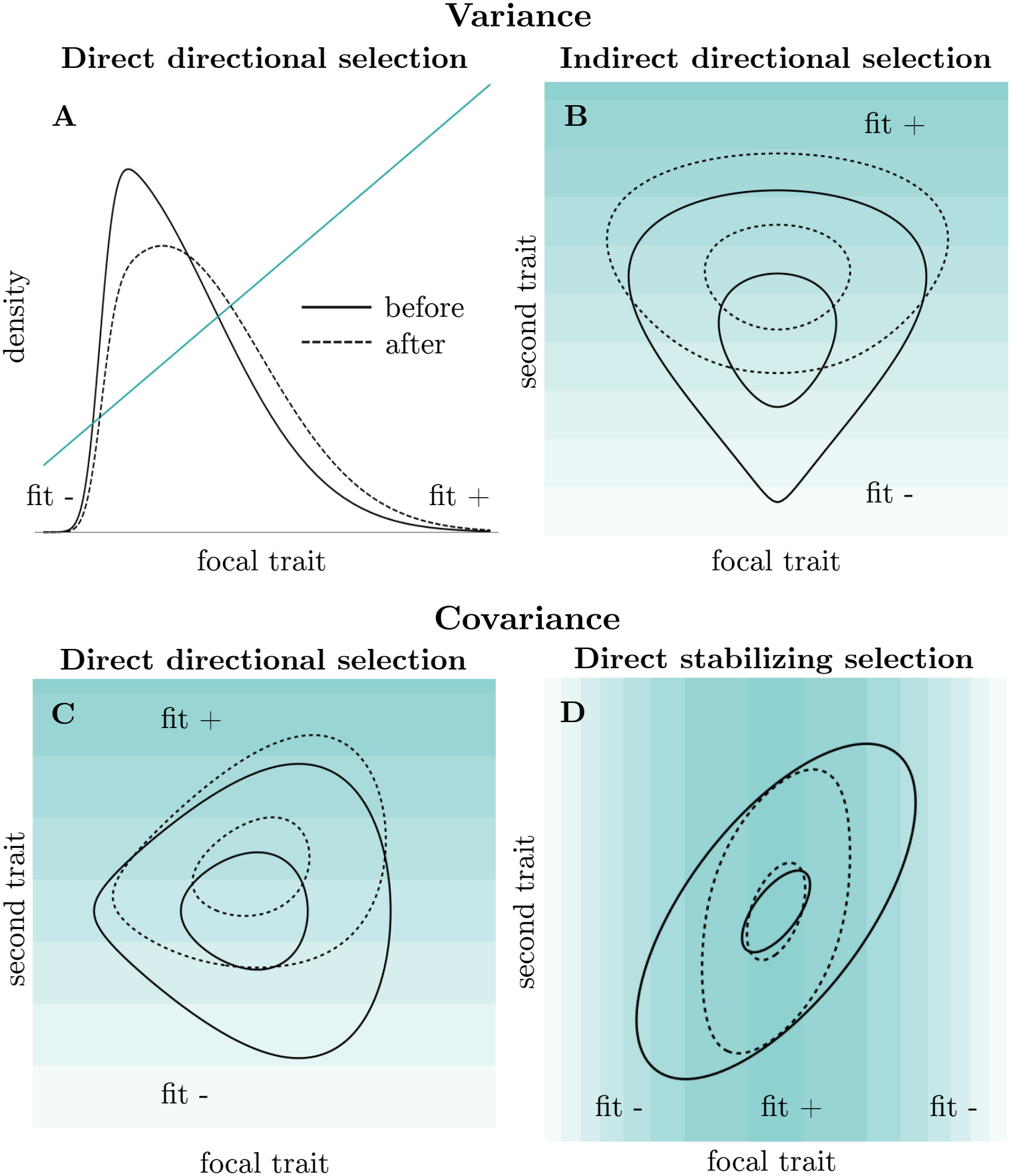
Change in variance and covariance due to selection alone. Each panel shows the change in the trait distribution over one generation, obtained numerically from eq. (6) without mutation. **A**. Variance under direct directional selection (blue line: local fitness land-scape). With the same positively skewed starting distribution, selection for smaller values would trim the upper tail and decrease variance (see main text). **B**. Variance under indirect directional selection: favoured individuals with high values at the second trait vary more in the focal trait. **C**. Covariance under direct directional selection on the second trait: favoured individuals also tend to carry high focal values. This is the main mechanism by which selection builds covariance between a modifier trait (here the focal trait) and a viability trait (see main text). **D**. Covariance erosion under direct homogenising selection. In **B–D**, the background colour shows the local fitness landscape (darker is fitter); larger and smaller ellipses circum-scribe 80 and 20 percent of the distributions, respectively. The code and numerical values for this figure are in Supplementary Mathematica Notebook 1. **Alt text:** Panel A shows directional selection trimming the long upper tail of a skewed distribution, which reduces its variance. Panel B shows the focal trait being more variable among favoured individuals, which increases its variance. Panel C shows directional selection on a second trait building positive covariance. Panel D shows homogenising selection eroding covariance.

In turn, the covariance 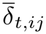 between two traits *i* and *j* can change under direct directional selection if one of these traits (say *j*) experiences directional selection (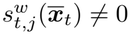) while its variation is associated with values of the other ( 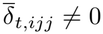; see tab. S1). This is because individuals with the most extreme values of trait *j* then also tend to carry high (or low) values of trait *i*, such that directional selection builds associations (see fig. 4C). Finally, indirect directional selection can also affect 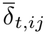 if the covariation between traits *i* and *j* is associated with a third trait *k* (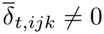), itself under directional selection (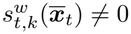; see tab. S1). In this case, covariation between traits *i* and *j* is higher (or lower) among individuals favoured by directional selection, leading to an increase (or decrease) in 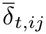 over one generation.

Beyond directional selection, the (co)variance 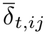 can also change due to second-order selection,

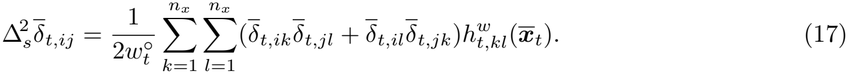

Similarly to eq. (11), eq. (17) has analogues in polygenic quantitative genetics models (Lande & Arnold 1983, eq. 15b to fourth order and without directional selection; Phillips & Arnold 1989, eq. 2 without directional selection) and under the continuum-of-alleles model (Débarre et al. 2014, eq. 6; Mullon & Lehmann 2019, eq. 3b without mutation and directional selection). It shows that second-order selection on any pair of traits *k* and *l* can affect the (co)variance between any pair of traits *i* and *j*, as long as there is more (or less) (co)variation between the latter traits among individuals with larger values of the product δ_*t,k*_ δ _*t,l*_ than in the population as a whole (an association approximated by the sum in parentheses in eq. 17 under our cumulant-neglect assumption). Here, such selection is referred to as indirect for elements of eq. (17) with *i≠ k, i≠ l, j≠ k* and *j≠ l*, and as direct otherwise.

We can see from eq. (17) that direct homogenising selection on a trait *i* (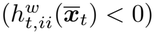) by itself always tends to erode its covariance with other traits. To see this, notice that terms corresponding to this component of selection then have factors whose sign is always opposite to that of 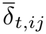 (e.g., 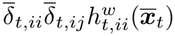; see tab. S1), and thus always drive covariance to zero (see fig. 4D). Indirect homogenising selection on a third trait *k* may have an opposing effect on 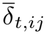, but that can never last since it requires that both 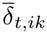 and 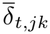 are non-zero, while both moments are eroded by direct homogenising selection on trait *k*. As a result, the only way that trait covariance can be maintained in the long term by second-order selection alone is via correlational 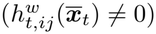 or diversifying selection 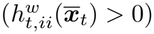.

Let us now turn to the component of change due to change in average traits, which is

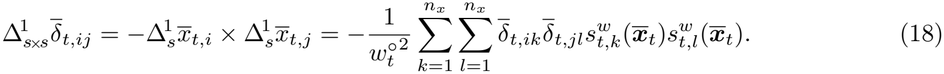

Analogues of eq. (18) can be found in previous work (Lande & Arnold 1983, eq. 15b; Phillips & Arnold 1989, eq. 2; Mullon & Lehmann 2019, eq. 3b). Mathematically, this component of change accounts for the change in the reference point used to calculate second-order central moments at generation *t* (i.e., 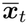) and *t* + 1 (i.e., 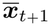). Accordingly, it plays a role only under directional selection (i.e., it vanishes when 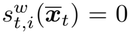 for all *i*), and is absent from analyses where (co)variance is calculated using the same reference point across generations (e.g., Barton & Turelli 1987; Rice 2002). It is also specific to discrete-time dynamics and vanishes under continuous-time dynamics (Lion et al. 2023). Note that although eq. (18) technically depends on directional selection, the mechanism it describes is fundamentally different from the mechanism described by eq. (16). When a distinction is needed we refer to the former as secondary directional selection components and to the latter as primary (this distinction is carried over to higher-order analogues of eq. 18 below).

When *i* = *j*, eq. (18) shows that secondary directional selection on a trait *i* can only decrease its variance, since 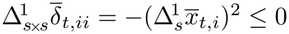. Concordant components of secondary directional selection (i.e., terms in eq. 18 where 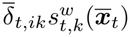 and 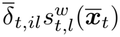 have the same sign) strengthen this decrease, while antagonistic components (i.e., terms where they have opposite signs) weaken it. With *i≠ j*, eq. (18) also shows that the covariance 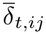 tends to decrease when traits *i* and *j* are selected to change in the same direction, that is, when the net changes in the averages of traits *i* and *j* due to directional selection have the same sign. Conversely, 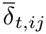 tends to increase when traits *i* and *j* are selected to change in opposite directions.

The component of change due to mutation alone is

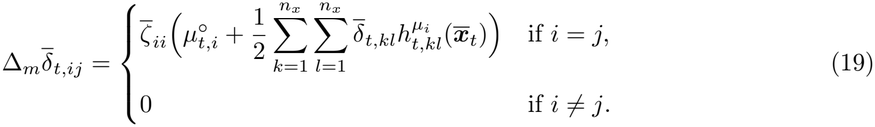

where 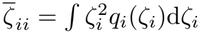 is the variance in mutation effects at locus *i*, 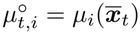, and

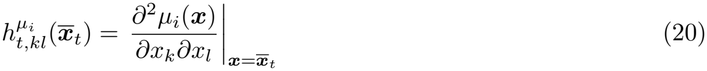

is a second-order mutation coefficient measuring the interaction between trait *k* and *l* in their effect on the mutation probability at locus *i*. When *k* = *l*, a positive (negative) value means that the effect of trait *k* on the mutation probability at locus *i* becomes more positive (more negative) as trait *k* increases, i.e., the mutation probability is convex (concave) in trait *k*. When 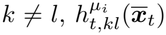 measures the interaction of traits *k* and *l* in determining the mutation probability at locus *i*, that is how the effect of trait *k* on that probability changes with trait *l*.

Eq. (19) shows that mutation cannot directly contribute to trait covariance (the case *i≠ j*). This follows directly from our assumption that mutations and their effects are independent across loci. The same equation shows, however, that mutation contributes to trait variance 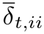, proportionally to the average mutation probability at locus *i* (the quantity between parentheses in eq. 19 is in fact a second-order approximation of the average mutation probability; see eq. B-68) and to the variance in their effects. An analogue of this term for the case of a constant mutation rate can be found in Mullon & Lehmann (2019, eq. 3b). In this previous model however, all traits are encoded by a single locus such that mutations are pleiotropic and can contribute to covariance.

Finally, the component of change due to the interaction of mutation and selection is

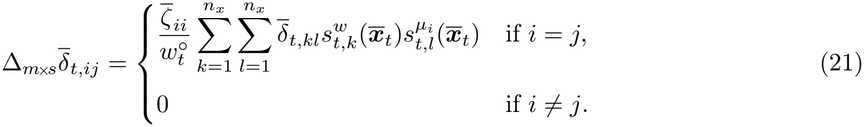

where

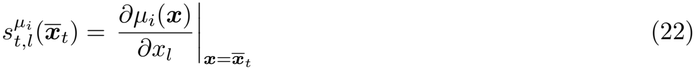

is a first-order mutation coefficient measuring the effect of trait *l* on the mutation probability at locus *i*. Positive (negative) values of that coefficient indicate that an increase in trait *l* increases (decreases) this probability.

Eq. (21) stems from the assumption that traits can affect mutation probabilities, such that traits and mutation probabilities can be associated in the population. In fact, the sum in this equation is the first-order approximation of the covariance between *w*_*t*_(***x, ϕ***_*t*_(***x***)) and *µ*_*i*_(***x***) (see eq. B-73). When fitness and the mutation probability at locus *i* covary, more successful individuals also produce mutations more (or less) frequently, contributing an extra change in trait variance, proportional to mutational variance.

### 3.3 Change in skewnesses and coskewnesses

We now turn to the dynamics of third-order trait moments. We show in appendix B.3.3 that primary directional selection does not change these moments. This is because, under our cumulant-neglect assumption and accounting for the change in average traits, all the corresponding terms cancel out (see eq. B-94). As a result, to leading orders the change 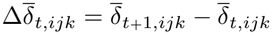 in the coskewness between traits *i, j* and *k* over one generation can be partitioned as

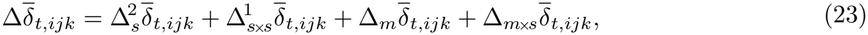

into the change 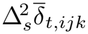 (order 5) due to second-order selection, the change 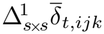 (order 5) due to change in average traits and (co)variance (via selection alone), the change 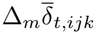 (order 4) due to mutation alone and the change 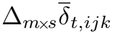 (order 5) due to the interaction of mutation and selection.

The change in 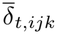 due to second-order selection is

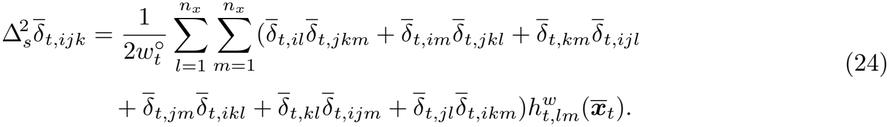

Terms analogous to eq. (24) are obtained in general descriptions of change in trait moments (Barton & Turelli 1987; Bürger 1991; Rice 2002; Rice 2020), but in a different form due to different genetic and closure assumptions. Eq. (24) shows that second-order selection on any pair of traits *l* and *m* can affect the (co)skewness between any three traits *i, j* and *k*, as long as the (co)skewness between the latter traits is larger (or smaller) among individuals with larger values of the product *δ*_*t,l*_*δ*_*t,m*_ than in the population as a whole (an association which, under the cumulant-neglect closure and once the change in average traits is accounted for, is approximated by the term in parentheses in eq. 24).

The main takeaway from eq. (24) is that when covariances are small (as they will be in a prolonged homogenising selection regime, see previous section), direct homogenising selection on a trait 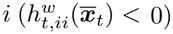 alone always erodes (co)skewness (see fig. 5A). To see this, observe that the components of change in 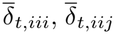 or 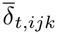 then only have terms whose sign is opposite to that of the corresponding (co)skewness (e.g., 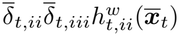 or 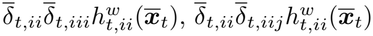; see tab. S2), such that the latter is pushed towards zero. As a result, the only way that asymmetries in the trait distribution can be maintained in the long term by selection alone is via correlational 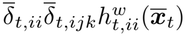 or diversifying selection 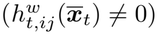.

**Fig. 5:**
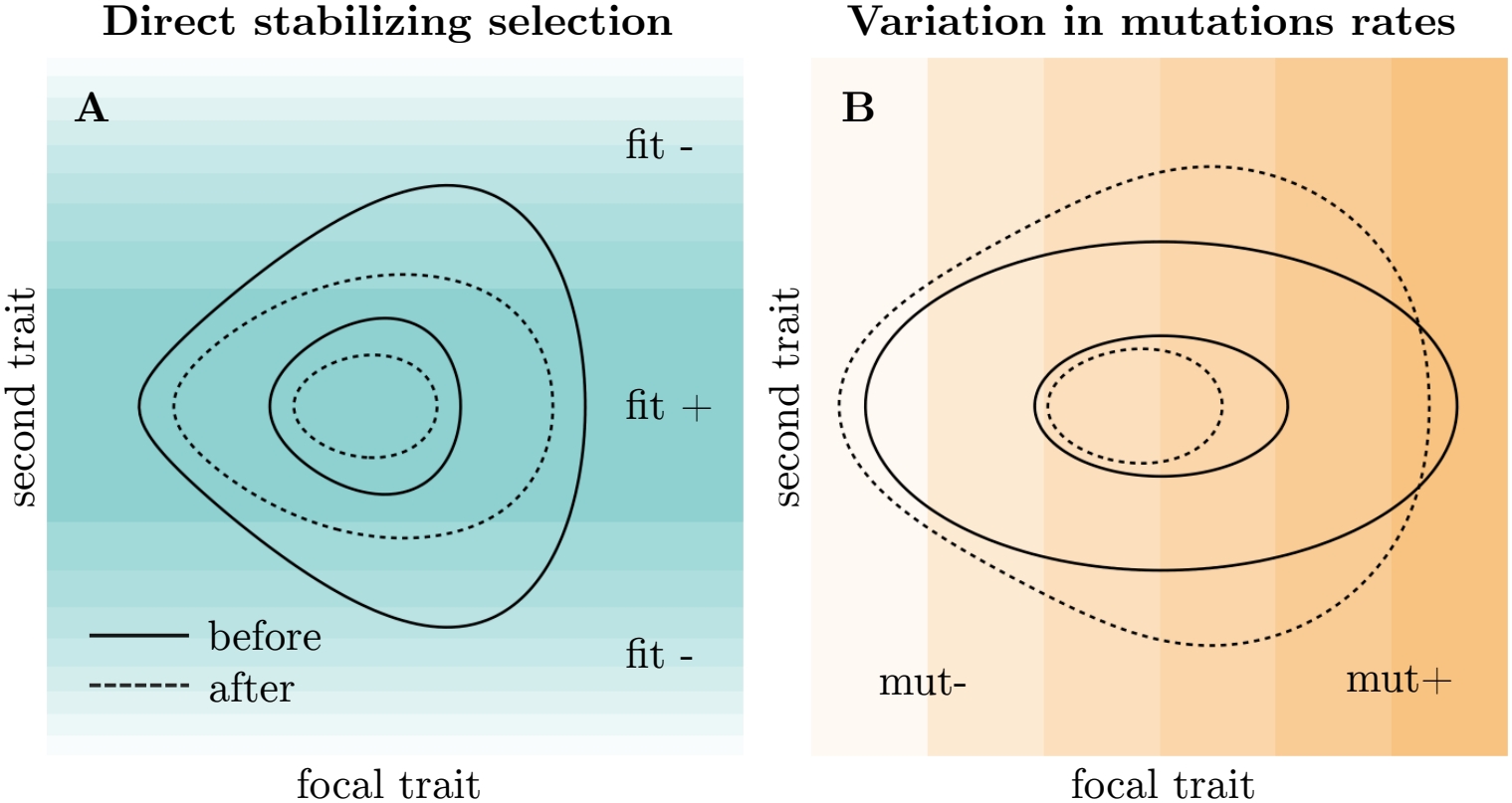
Change in two-way coskewness due to homogenising selection and mutation. Each panel shows the change in the trait distribution over one generation, obtained numerically from eq. (6). **A**. Coskewness erosion under homogenising selection. **B**. In the absence of selection, variation in mutation probabilities builds an association between a focal modifier trait and variation at another trait. Here the focal trait affects the mutation probability at the locus encoding the second trait linearly, and mutation is very strong for illustration. The background colour shows the local fitness landscape in **A** (darker is fitter) and the local mutation rate in **B** (darker is higher). Larger and smaller ellipses circumscribe 80 and 20 percent of the distributions, respectively. The code and numerical values for this figure are in Supplementary Mathematica Notebook 1. **Alt text:** Panel A shows homogenising selection eroding the coskewness of a two-trait distribution on a shaded fitness landscape. Panel B shows that, in the absence of selection, mutation whose rate depends on the focal trait makes variation in the second trait differ across values of the focal trait, building coskewness.

Besides the role of second-order selection above, eq. (23) contains a component of change due to change in average traits and (co)variance (themselves due to selection, and given by eqs. 11 and 16),

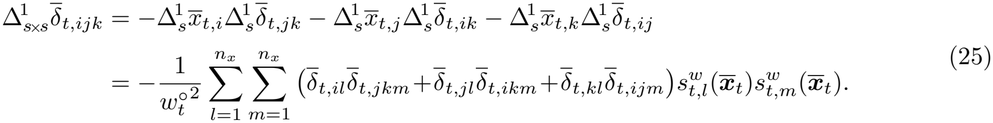

Similarly to eq. (18) for (co)variances, eq. (25) accounts for the first order of change in the reference point used to calculate third-order central moments in generations *t* (i.e., 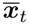) and *t* + 1 (i.e., 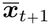). Accordingly, these secondary components act only under directional selection (i.e., vanish when 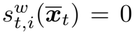 for all *i*), and are absent from analyses where (co)variances and (co)skewnesses at generations *t* and *t* + 1 are calculated using the same reference point (Barton & Turelli 1987; Rice 2002).

Eq. (25) shows that the change in the average of any trait *i* due to directional selection affects the change in the (co)skewness 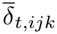 proportionally to the simultaneous change in the (co)variance 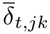 due to directional selection. For instance, the element of the right-hand side of eq. (25) with *i* = *j* = *k* = *l* = *m* says that the skewness 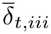 will have a component of change proportional to 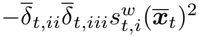, that drives it to zero. We can thus see that secondary directional selection on a trait contributes to eroding skew in its distribution. Likewise, eq. (25) shows that the two-way coskewness 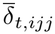 will have a component of change proportional to 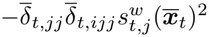, that drives it to zero. This means that secondary directional selection on a viability trait *j* tends to erode any association between its variation and another trait *i*.

Beyond the effect of selection alone, the component of change in trait (co)skewness due to mutation alone is

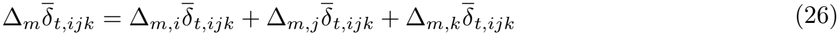

with

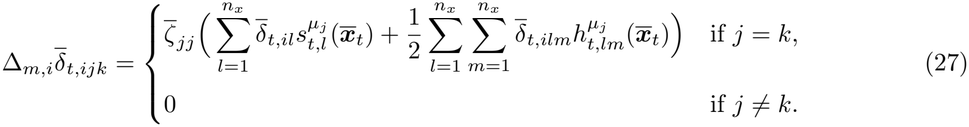

Eqs. (26) and (27) show that mutation alone cannot directly contribute to trait three-way coskewness (i.e., 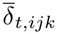 with *i≠ j≠ k* all distinct). This follows directly from our assumption that mutationsand their effects are independent across loci. Mutation can, however, contribute to building skewness and two-way coskewness, when there are associations between trait values and mutation probabilities. In fact, the quantity between parentheses in eq. (27) is the third-order approximation of the covariance between trait *i* and the mutation probability *µ*_*j*_(***x***) (see eq. B-75). When trait *i* and *µ*_*j*_(***x***) covary, individuals with larger (or smaller) values of trait *i* mutate more often at locus *j*, and thus accumulate new trait variation in trait *j* at a faster pace, leading to the build-up of asymmetries (see fig. 5B). Note that with strictly positive trait and mutational variances, this mechanism will always be active when trait *i* affects *µ*_*j*_(***x***), in which case it represents a steady source of asymmetries in the trait distribution, active irrespective of the action of selection.

Finally, the component of change in trait (co)skewness due to the interaction of mutation and selection is

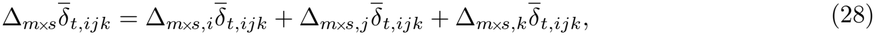

where

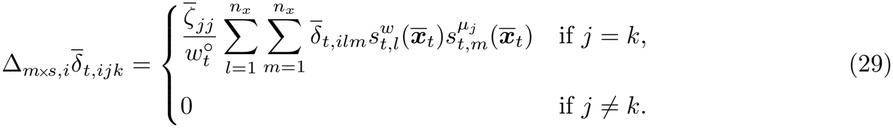

When there already are asymmetries in the trait distribution and the population is under directional selection, eqs. (28) and (29) provide an extra source of trait skewness and two-way coskewness that stems from three-way associations between trait values, fitness and mutation probabilities. In fact, the sum in eq. (29) is the third-order approximation of the third-order moment measuring these associations (see eq. B-74). When this moment is positive (negative), it means that the association between fitness and the mutation probability at locus *j* is more positive (more negative) among individuals with larger values of trait *i* than among those with smaller values, such that selection amplifies (dampens) the input of (co)skewness from mutation.

### 3.4 Orders of change and selection on viability traits

#### 3.4.1 Closed system of moments

Eqs. (10)–(29) provide a closed dynamical system of equations describing the effect of mutation and selection on trait averages, (co)variances and (co)skewnesses. The components of this system display a hierarchy of orders in the parameter *ϵ* (see tab. 1), with lower-order components tending to be larger than higher-order ones. Thus, we can expect the former components to dominate (i.e., largely determine) the moment dynamics, which is useful to identify the leading causes of change in properties of interest (here the mutation probabilities; see section 3.5).

**Tab. 1:** Orders of change in trait moments. The table gives the order (in trait deviations *ϵ*) of each component of change in trait moments. Orders are given assuming that mutation probabilities can be large relative to trait deviations. When they are small, terms in brackets become of higher orders. Greyed-out terms are active only when traits experience directional selection.

|  | **Orders** |  |  |  |
|---|---|---|---|---|
|  | 2 | 3 | 4 | 5 |
| $\Delta\overline{x}_{t,i}$ | $\Delta_s^1\overline{x}_{t,i}$ | $\Delta_s^2\overline{x}_{t,i}$ |  |  |
| $\Delta\overline{\delta}_{t,ij}$ | $[\Delta_m\overline{\delta}_{t,ij}]$ | $\Delta_s^1\overline{\delta}_{t,ij}$ | $\Delta_s^2\overline{\delta}_{t,ij}$, $\Delta_{s\times s}^1\overline{\delta}_{t,ij}$, $[\Delta_{m\times s}\overline{\delta}_{t,ij}]$ |  |
| $\Delta\overline{\delta}_{t,ijk}$ |  |  | $[\Delta_m\overline{\delta}_{t,ijk}]$ | $\Delta_s^2\overline{\delta}_{t,ijk}$, $\Delta_{s\times s}^1\overline{\delta}_{t,ijk}$, $[\Delta_{m\times s}\overline{\delta}_{t,ijk}]$ |

Which components of the system actually dominate the dynamics in a specific case also depends on the relative size of mutation probabilities and selection coefficients, relative to trait deviations. Change in the average of a trait *i* is dominated by directional selection 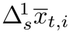 as long as the population is far from any singular trait, and the quadratic selection component 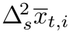 can thus be neglected until directional selection vanishes. Change in the variance is dominated by the mutation component 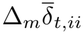 when the mutation probability is large relative to trait deviations, and other components of change can then be neglected. When mutation probabilities are small, however, selection components must be taken into account. In that case, change in 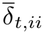 far from singular traits is determined by primary directional selection (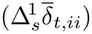) if the trait distribution is skewed, or by the combination of secondary directional selection 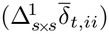 second-order selection 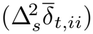 and mutation-selection interactions 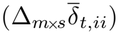 otherwise. Close to a singular trait, second-order selection is the only active selective mechanism; when it decreases trait variance (i.e., homogenising selection), it may balance out mutational input (variance is reduced until mutational input is just as strong as selection), leading to a mutation-selection balance. Since mutation does not directly affect covariance, its dynamics obey the same hierarchy as the variance, but with the mutation-related components 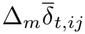 and 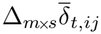 removed, such that no mutation-selection balance can exist for trait covariance.

The hierarchy of components for (co)skewnesses is simpler than for (co)variances, as primary directional selection does not play a role. When mutation probabilities are large relative to trait deviations and vary with traits, change in skewness 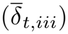 and two-way coskewness 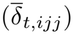 is dominated by the mutation component 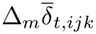, and other components of change can be neglected (i.e., skew simply builds up due to mutational input). When mutation probabilities are small or constant however, selection components must be included. In that case, change in (co)skewness far from singular traits is determined by secondary directional selection 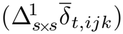, second-order selection 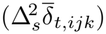 and mutation-selection interactions 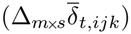 together, or by second-order selection alone near singular traits. When the latter erodes skew (i.e., homogenising selection), it may balance out mutational input (asymmetries are reduced until mutational input is just as strong as selection). Since mutation does not directly affect three-way coskewnesses ( 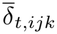 with *i, j* and *k* distinct), their dynamics obey the same hierarchy, but with mutation-related components (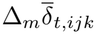and 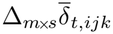) removed, such that no mutation-selection balance can exist for three-way coskewnesses.

#### 3.4.2 Viability traits only

Our next aims is to qualitatively understand the selective mechanisms acting on modifier traits. When modifier traits have no direct fitness effects however, their evolution will depend entirely on indirect selection pressures, operating via two-way associations with co-evolving viability traits. Therefore, it is useful to first recall the dynamics of viability traits.

For viability traits, all mutation coefficients (eqs. 20 and 22) are zero and eqs. (21), (26) and (29) vanish. In the absence of fluctuating selection, trait averages first evolve by directional selection towards the closest evolutionary attractor ***x***^*\**^ where 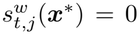 for all *j* (i.e., a convergence stable singular trait value). At such a point, and since there are no sources of asymmetries in the trait distribution (eq. 26 is not active), indirect second-order selection (eq. 13) cannot act and (co)variances are shaped by direct second-order selection (via eq. 17) and neutral mutational input (via eq. 19). Two qualitatively different evolutionary outcomes are then possible, depending on the values of second-order selection coefficients. If selection is homogenising on all traits (i.e., 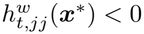 for all *j*) and correlational selection on pairs of traits is not too strong, second-order selection is globally stabilising, and eventually balances mutational input such that variance in trait *j* settles at an equilibrium (e.g., Débarre et al. 2014; Mullon & Lehmann 2019).

If, on the other hand, quadratic selection is diversifying on some traits at the attractor (i.e., 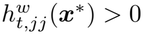 for some *j*) or correlational selection on some pairs of traits is strong enough, second-order selection is disruptive along at least one trait dimension (i.e., Débarre et al. 2014; Mullon & Lehmann 2019). These terms then imply that trait variance along the corresponding dimensions will increase rapidly under the coupled effect of mutational input and selection. This can lead to evolutionary branching and to the coexistence of several distinct morphs (i.e., several modes in the trait distribution; see Geritz et al. 1998; Kisdi & Geritz 1999 for examples of evolutionary branching). Such diversification quickly leads to the violation of our working assumption that trait variation is small, such that the evolutionary dynamics in polymorphic populations cannot be predicted from our model.

### 3.5 Selection on modifier traits

#### 3.5.1 Modifier traits without direct fitness effects

Let us now introduce a modifier trait (say trait *i*) affecting the mutation probability at a viability locus (say *j*), such that the corresponding mutation coefficients (eqs. 20 and 22) are non-zero and eqs. (21), (26) and (29) affect the dynamics. Then, eq. (26) predicts a change in the two-way coskewness 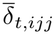 proportional to 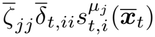. Since variances are positive, this term generates positive (negative) values of 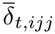 whenever trait *i* increases (decreases) *µ*_*j*_(*x*_*i*_). Once generated, this asymmetry allows for selection pressures acting on viability trait *j* to translate into a change 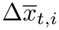 in trait *i*. Such indirect selection, however, will act via different selective mechanisms (mediated by either 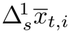 or 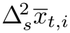) and lead to qualitatively different outcomes depending on whether the viability trait experiences homogenising, diversifying, or directional selection (see tab. 2).

**Tab. 2:** Selective mechanisms acting on mutation probabilities.

| **Mechanism** | **How it acts** | **Mutation probabilities** |
|---|---|---|
| **Directional selection on a modifier trait** | A trait affecting mutation probabilities also affects fitness, so selection acts directly. Examples include costs to fidelity and other functional trade-offs. | Increase or decrease depending on the fitness effect |
| **Directional selection on a viability trait** | Mutator alleles produce and associate with more new variants. Variants that shift the trait in the favoured direction spread, and mutators spread along with them (genetic hitch-hiking). | Increase |
| **Homogenising selection on a viability trait** | Mutator alleles produce and associate with more extreme viability trait values, which are disfavoured (the reduction principle). | Decrease |
| **Diversifying selection on a viability trait** | Mutator alleles produce and associate with more extreme viability trait values, which are favoured. | Increase |

##### Indirect directional selection favours higher mutation probabilities

When the viability trait *j* is under directional selection, such that 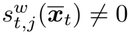, a non-zero two-way coskewness 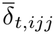 means that the covariance 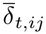 has a component of change proportional to 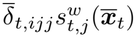, favouring positive values of 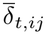 when 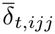 and 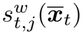 have the same sign, and negative values otherwise (eq. 16). Direct selection on trait *j* thus associates values of trait *i* producing higher mutation probabilities (large or small values when 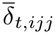 is positive or negative, respectively) with values of trait *j* producing higher fitness (large or small values when 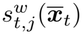 is positive or negative, respectively). As a consequence of non-zero values of 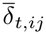, directional selection on trait *j* can then act on the average of trait *i*: the term 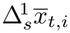 has a component of change 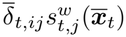 (see eq. 11 and fig. 2B). This shows that, by favouring values of trait *j* with higher fitness, directional selection also favours values of trait *i* producing higher mutation probabilities. This mechanism is analogous to the genetic hitch-hiking of mutator alleles under positive selection (e.g., Eshel 1973; Gillespie 1981b; Ishii et al. 1989; Taddei et al. 1997b; Johnson 1999b; Carja et al. 2014; Fyon & Úbeda 2025).

##### Indirect homogenising selection favours lower mutation probabilities

If viability trait *j* is under homogenising selection, such that 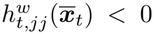, the term 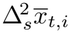 has a component of change 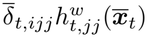 whose sign is always opposite to that of 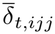 (see eq. 13 and tab. S1). This component thus favours either an increase in trait *i* when this trait decreases *µ*_*j*_(*x*_*i*_) (such that 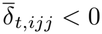, as detailed above), or a decrease in trait *i* when this trait increases *µ*_*j*_(*x*_*i*_) (such that 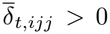; see fig. 3C), in both cases favouring a decrease in *µ*_*j*_(*x*_*i*_). This result recovers the “reduction principle” for mutation probabilities (Liberman & Feldman 1986; Liberman et al. 2016; Altenberg et al. 2017), as well as related results showing indirect selection for lower mutation probabilities in large populations at the mutation-selection equilibrium (e.g., Leigh 1973; Gillespie 1981a; Kondrashov 1995). In a population with variable mutation probabilities and where mutations are bad on average, trait values responsible for larger rates become associated with larger load (here, larger trait variation around the optimum) and are selected against. Interestingly, this effect is of order three in trait deviations, while the change due to indirect directional selection is of order two. This means that, if covariance is large enough and the landscape is steep enough, higher mutation probabilities can be selected for even in a concave fitness landscape, where mutations decrease individual fitness on average.

##### Indirect diversifying selection favours higher mutation probabilities

Our model also shows that a symmetrical mechanism operates when trait *j* is under diversifying selection, such that 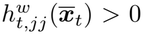, favouring increased mutation probabilities instead (see fig. 3D for the case where 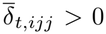). This will typically occur at evolutionary branching points and this effect corresponds to the general expectation that traits increasing phenotypic variance in offspring should be favoured when individual fitness is convex in the trait of its bearer (Bruijning et al. 2020), and may help explain previous results showing the evolution of increased mutation probabilities under some forms of negative frequency-dependent selection (Holsinger et al. 1986).

#### 3.5.2 Modifier traits with direct fitness effects

In addition to indirect selection, modifier traits can also be subject to direct selection pressures whenever they affect viability themselves (see tab. 2). Such effects may arise, for example, because DNA repair, replication fidelity or germline maintenance come at a metabolic cost for the organism. They can also arise if higher-level traits affecting mutation probabilities (e.g., mutagen avoidance) also impact other life-history functions (e.g., foraging efficiency). In any case, the direct effects on fitness can generate direct selective effects on the average of the modifier trait (via eq. 11) that either agree or oppose those of indirect selection described above. When these effects oppose, and depending on the magnitude of each effect involved, the average modifier trait may evolve and remain at an intermediate value where both direct and indirect selective mechanisms balance out. For instance, stable intermediate mutation probabilities are predicted if viability traits are under homogenising selection (favouring reduced mutation probabilities) but reducing mutation probabilities comes at a cost. This recovers previous results describing such equilibria (e.g., Kondrashov 1995; Dawson 1998).

## 4 Example two-trait model

We now study the simple example of a two-trait model, with one viability trait and one modifier trait. In this example, the complete system of trait moments reduces to its core components (appendix C.1), allowing us to display its inner workings.

### 4.1 Assumptions

The first trait considered, referred to using a subscript ‘1’, is a viability trait that determines fecundity in the absence of density-dependent competition and when the trait alone determines reproduction. Specifically, we assume that this baseline fecundity of a focal individual with viability trait *x*_1_ decreases exponentially with the squared distance of *x*_1_ to an optimal value *z*_*t*_,

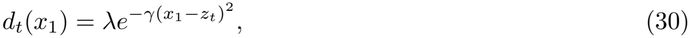

where *γ ≥* 0 and *λ* are two parameters controlling the importance of matching the trait to *z*_*t*_, and the baseline fecundity under a perfect match, respectively. In eq. (30), the subscript *t* indicates that *z*_*t*_, and thus *d*_*t*_(*x*_1_), can change from one generation to the next, leading to a changing fitness landscape.

The second trait, denoted by a subscript ‘2’, is a modifier trait that controls the mutation probability at both loci. More specifically, we assume that the mutation probability at both loci is

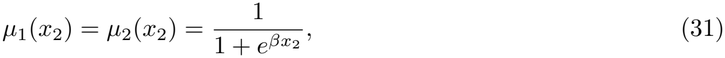

where the parameter *β ≥* 0 controls the sensitivity of the mutation probability to *x*_2_. Since eq. (31) decreases monotonically with *x*_2_, the modifier trait here can be thought of as an effort towards transmission fidelity (e.g., some energy investment in producing an accurate DNA polymerase).

In terms of the above, we assume that the fitness of a focal individual with traits ***x*** = (*x*_1_, *x*_2_) is

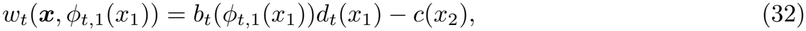

where

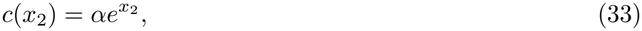

is a cost of effort towards transmission fidelity where *α ≥* 0 is a parameter. When *α* = 0, *x*_2_ has no direct effect on individual fitness. Finally

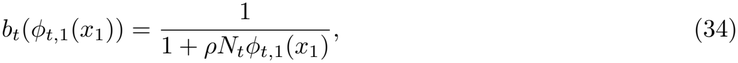

describes density-dependent competition, with the parameter *ρ ≥* 0 modulating its importance. Here *N*_*t*_*ϕ*_*t*,1_(*x*_1_) is the ‘effective population density’ perceived by a focal individual with viability trait *x*_1_, modulated by the regulating factor

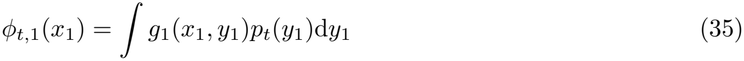

with

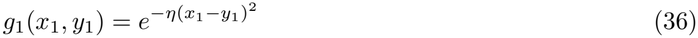

being a competition kernel giving the contribution of an individual with viability trait *y*_1_ to the total competitive pressure acting on the focal, with the parameter *η ≥* 0 controlling how narrow the kernel is (i.e., how similar individuals must be to compete). Larger values of *η* make competition more local (dissimilar individuals matter less for the focal); with *η* = 0, *ϕ*_*t*,1_(*x*_1_) = 1 and every individual perceives the same effective population density *N*_*t*_.

### 4.2 Selection coefficients

Injecting eq. (32) into eq. (12) and using the fact that here 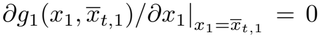 (due to the symmetry of the competition kernel; see eq. 36) we have

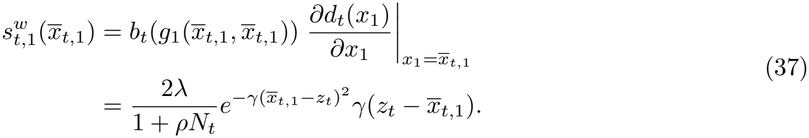

Hence, 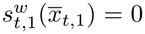 obtains when 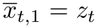. Note that in the example model, individual fitness and its derivatives depend on the vector of average traits 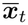 only through one of its two entries (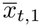for 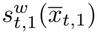 and 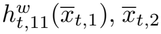 for 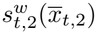 and 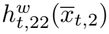, while mutation probabilities and their derivatives depend only on 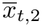. We make this explicit throughout by writing each of these quantities as a function of that single average trait (e.g., 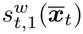) becomes 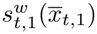, and 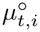 becomes 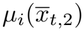).

Injecting eq. (32) into eq. (14) and making use of the facts that 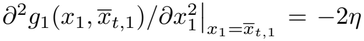 and that 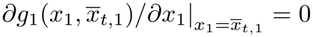, we also have

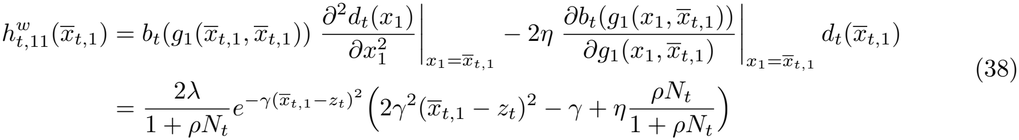

showing that 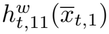 depends on the curvature of the efficiency function and on the effect of the regulating factor. The latter is always negative (i.e., 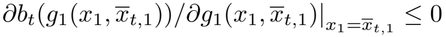) and thus tends to make selection diversifying on the first trait. From eq. (38), we see that, when 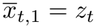, selection is diversifying (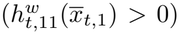) if and only if *γ < ηρN*_*t*_*/*(1 + *ρN*_*t*_), meaning that competition must be stronger than the necessity to match *z*_*t*_.

Besides selection on the first trait only, injecting eq. (32) into eqs. (12) and (14) gives 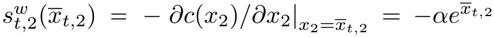 and 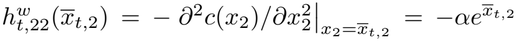 (the two coincide because the cost function is an exponential), meaning that selection acts on the second trait directly only when *α≠* 0. It also gives 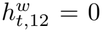, showing that our choice of fitness function implies the absence of correlational selection.

### 4.3 System of moments

Injecting eqs. (31) and (32) into eqs. (10)–(29) yields a reduced system of moments. To further simplify the presentation, we describe this system under the additional assumptions that the modifier trait has no direct fitness effects (i.e., no cost, *α* = 0) such that 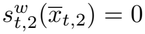 and 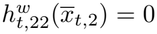, and that it has only linear effects on the mutation probability, such that 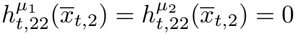. The full system of moments used in numerical analyses below is available in appendix C.1. Note that in this appendix we also give the small-variance approximation of eq. (5), used for tracking the dynamics of total population density.

#### Means

The average of the viability trait changes according to

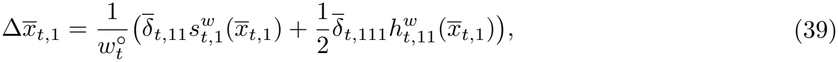

where

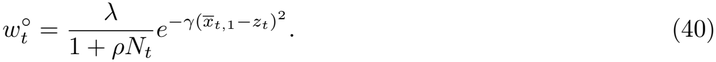

Eq. (39) shows that trait 1 first evolves in response to direct directional selection. When directional selection vanishes and the distribution of trait 1 is skewed, quadratic selection can also act, but without significant consequences since any induced change quickly reintroduces first-order selection. This is consistent with the idea that second-order selection on average traits can be ignored when studying systems of viability traits only.

The average of the modifier trait changes according to

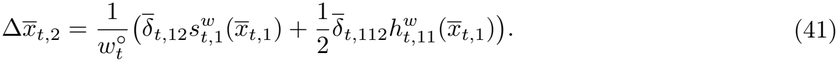

Eq. (41) is functionally equivalent to eq. (39), but its two components of selection are indirect. Trait 2 first evolves in response to indirect directional selection, if it is associated with trait 1 such that 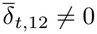. When directional selection vanishes and 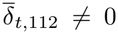, indirect quadratic selection can bring additional change in the average of trait 2. The latter change can be significant in this case, since it does not restore any first-order effect as above.

#### Variances and covariances

Variance in the viability trait changes according to

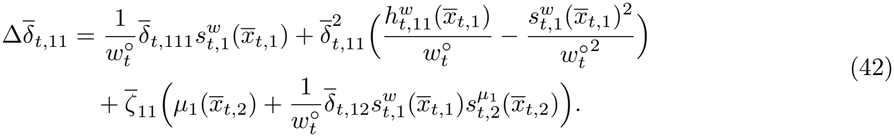

Thus, the variance in trait 1 changes because of selection (first line) and due to mutational input (second line). The leading term under small variance is the (second-order) neutral mutational input 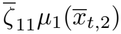, continuously introducing new variation. This term can be complemented by a second (fourth-order) mutation term active when 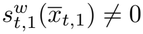 and the traits covary, such that variation in fitness is associated with variation in mutation probabilities, and fitter individuals mutate more (or less) often. In turn, when variation in trait 1 is large enough, selection acts on 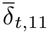 as well. Directional selection acts first via a third-order term, but only if the distribution of trait 1 is skewed. It acts more consistently towards a reduction in 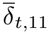 via a fourth-order negative term, proportional to 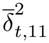. Quadratic selection acts only via a fourth-order term whose effect changes with the sign of 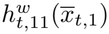: homogenising selection 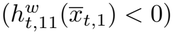 and diversifying selection 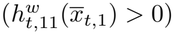 decrease or increase variance, respectively.

Variance in the modifier trait changes according to

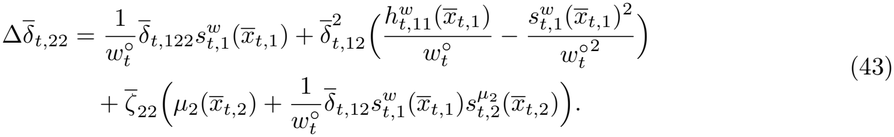

The mutation-related components of eq. (43) are equivalent to those of eq. (42). The selection-related components, however, differ in that they are indirect and require associations between the two traits. Directional selection on trait 1 affects 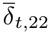 only when 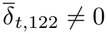 (i.e., trait 1 is associated with variation in trait 2) or when 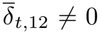, while quadratic selection on trait 1 affects 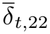 only when 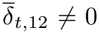. But even if selection on 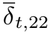 is indirect, its effects are qualitatively the same as in the direct case: directional selection promotes a reduction in 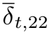 unless individuals favoured by directional selection are more diverse (i.e., when 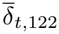 and 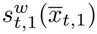 have the same sign); while homogenising and diversifying selection promote a reduction and increase in 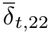, respectively.

The covariance between the two traits changes according to

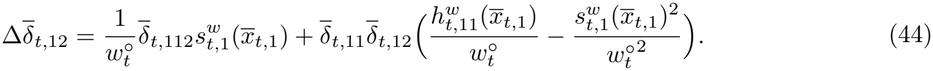

Eq. (44) has no mutation component, since mutation cannot directly contribute to trait covariance (see eq. 19), and covariance thus responds to selection only. In particular, if trait 1 is under directional selection, covariance changes proportionally to 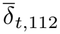. This is the only source of new covariance, since the two other components are zero when 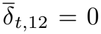, and since 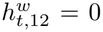. Thus, eq. (44) shows that for covariance between the traits to evolve, enabling the action of indirect directional selection on the average of trait 2 (see eq. 41), it is required that the two-way coskewness 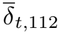 is non-zero.

#### Skewnesses and coskewnesses

Skewness in the viability trait changes according to

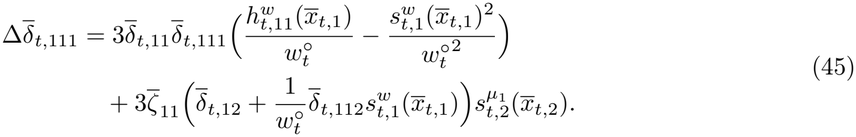

Eq. (45) shows that skewness in trait 1 changes because of selection (first line) and due to mutational input (second line). The leading term under small variance is the (fourth-order) neutral mutational input, which affects skewness in trait 1 whenever 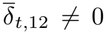, such that trait 1 covaries with mutation probabilities. This term is complemented by a second (fifth-order) selection-related mutation term that acts when 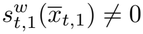 and the favoured individuals have non-average mutation probabilities 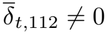. Selection acts on 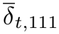 via fifth-order terms only. We can see that while most selection pressures on trait 1 (homogenising and directional) erode 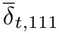, diversifying selection can inflate pre-existing skew.

Skewness in the modifier trait changes according to

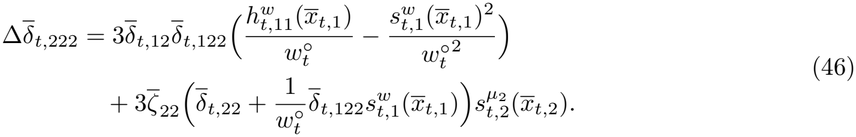

Mutation-related components of change in eq. (46) (second line) are functionally equivalent to those of eq. (45). The main difference is that the neutral contribution of mutation to 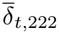 does not require association between the traits, but only that trait 2 varies. Selection-related components (first line) are also similar, but now describe indirect selection. For (secondary) directional or quadratic selection on trait 1 to affect 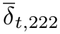, it is required that the traits covary, and that trait 1 is associated with variation in trait 2.

The coskewness 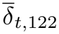, which measures associations between values of trait 1 and variation in trait 2, changes according to

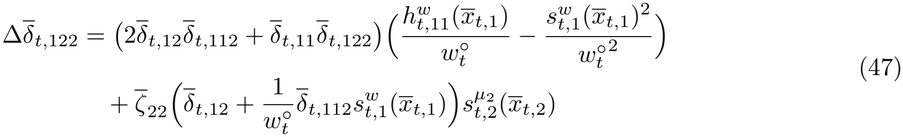

As in eq. (45), here mutation contributes to 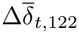 only if 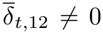 or 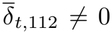. In that case trait 1 covaries with the mutation probability at locus 2, such that some values of trait 1 become associated with more variation in trait 2. Eq. (47) also shows that both directional and quadratic selection can change 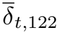 through the factor 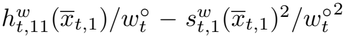. When this factor is negative (when selection is homogenising, or if directional selection is strong enough to outweigh any positive curvature), selection pushes 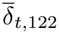 towards zero, unless 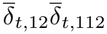 is of opposite sign to 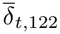 and large enough (i.e., 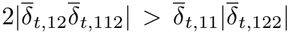), in which case selection amplifies 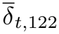. When the factor is positive (diversifying selection strong enough relative to directional selection), the two cases are reversed.

The coskewness 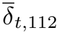, which measures associations between values of trait 2 and variation in trait 1, changes according to

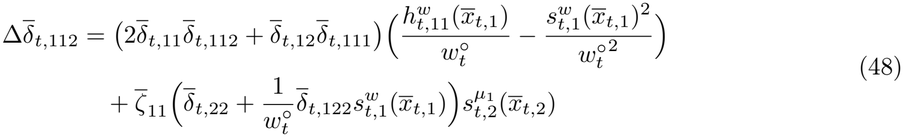

Both selection- and mutation-related components of change in eq. (48) can be understood in the same way as in eq. (47). A fundamental difference, however, resides in the neutral contribution of mutation. Here, this contribution does not require that the traits covary, but only that trait 2 varies and affects the mutation probability at locus 1, such that some values of trait 2 become associated with more variation in trait 1. Because these conditions are always true in our model, mutation constantly builds association between values of trait 2 and variation in trait 1.

### 4.4 Evolutionary scenarios

In this section, we analyse the dynamics induced by the above system under two evolutionary scenarios, one of transient directional selection and one of fluctuating selection. We then compare the resulting deterministic predictions to individual-based simulations (see implementation details in appendix C.2) of finite populations with stochasticity in individual fitness (i.e., reproduction is a Poisson process with intensity equal to the fitness). This allows for a qualitative assessment of the robustness of our results to the presence of genetic drift, which plays an important role in the evolution of mutation probabilities.

#### 4.4.1 Transient directional selection can temporarily increase mutation probabilities

In this scenario we assume that the average viability trait is initially far from a single constant optimal value *z*_*t*_ = *z*_op_. This leads to two distinct phases: transient directional selection on the viability trait as its average evolves towards *z*_op_, followed by homogenising selection once it is reached. These phases let us observe the separate effects of directional and homogenising selection, in interaction with the cost to transmission fidelity.

Initially, the population has size *N*_0_ = 10^4^ and traits follow a bivariate normal distribution, with means 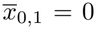 and 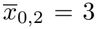, variances 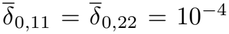, and no associations between the traits 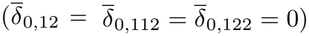. With *β* = 2, the average mutation probability at both loci is initially approximately 2.5 *×* 10^*−*3^, and mutational variances are set to 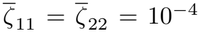. The viability optimum is *z*_op_ = 2, two trait units away from the initial average of the viability trait, and competition is set to be weaker than the necessity to match *z*_op_ (*η* = 10^*−*2^ and *γ* = 10^*−*1^). The density-dependence parameter is set to *ρ* = 10^*−*5^ and the basal growth rate to *λ* = 3. Together, these choices imply transient directional selection on the viability trait, followed by homogenising selection once the population is near the optimum. The population grows even in the first generations (i.e., even though it is maladapted), and reaches approximately 2 *×* 10^5^ individuals near the viability optimum.

In numerical iterations of the system of moments, we observe an immediate decrease in the average of trait 2 (fig. 6A, orange lines), and thus an increase in mutation probabilities (fig. 6C), in response to directional selection directly after initialisation. This is because the two-way coskewness 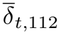 immediately starts decreasing, as parts of the population with lower values of trait 2 mutate more often and accumulate more trait variation (fig. 6B). Negative values of 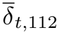 then translate into negative values of 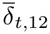 (i.e., to associations between more mutagenic trait values and fitter viability trait values), via the effect of directional selection (fig. 6D). These associations lead to the hitch-hiking of smaller values of trait 2 (fig. 6A, orange line), as trait 1 evolves towards the optimum, increasing the mutation probability. Once the population is close to *z*_op_ and directional selection has stopped, 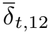 is eroded by direct homogenising selection, and quickly evolves back to zero. The change in the average of trait 2 then becomes dominated by indirect homogenising selection, favouring larger trait values and thus a reduction in the mutation probability. In the absence of cost to fidelity, this results in the mutation probability evolving ever closer to zero (i.e., as per the reduction principle). In the presence of costs, a stable intermediate long-term mutation probability is obtained instead (fig. 6, dashed lines).

**Fig. 6:**
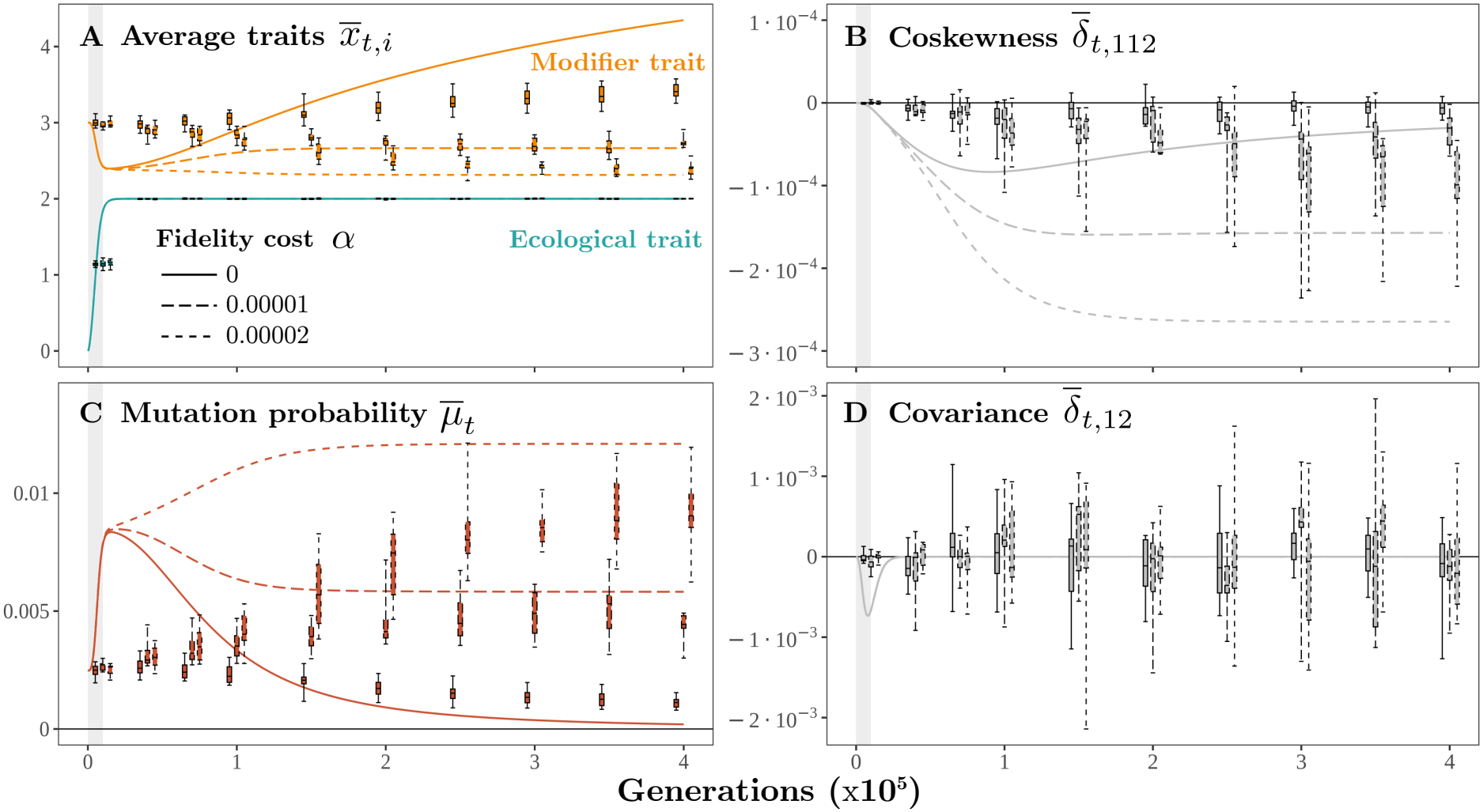
Transient evolutionary dynamics following environmental change. The population starts with a mean viability trait (trait 1) two trait units from the optimum *z*_*t*_ = 2 and evolves for 400 000 generations. After *∼* 10000 generations of directional selection (grey shading), it reaches the optimum, where it experiences constant homogenising selection. The modifier trait (trait 2) controls the mutation rate at both loci under varying costs (line types: fidelity cost *α*). **A**. Mean viability trait (blue) and modifier trait (orange). **B**. Two-way coskew-ness between the modifier trait and variation at the viability trait. **C**. Mean mutation rate. **D**. Genetic covariance between the viability and modifier traits. Lines show deterministic predictions obtained by iterating the system of moments; boxplots give the median, interquartile range and total range across 10 independent individual-based simulations. Other fixed parameters: *β* = 2; *λ* = 3; *γ* = 0.1; 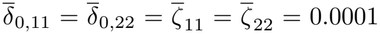; *η* = 0.01; *ρ* = 10^*−*5^, leading to a population of *∼* 2 *×* 10^5^ individuals. **Alt text:** Time series over 400 000 generations. Panel A shows the mean viability and modifier traits, panel B two-way coskewness, panel C mean mutation rate and panel D covariance. Lines give deterministic predictions for several fidelity costs, and boxplots give 10 individual-based simulations. A grey band marks the first 10 000 generations, during which the population approaches the optimum.

When the same scenario is explored in individual-based simulations, we observe qualitatively different dynamics (fig. 6, boxplots). There, the average of trait 1 also evolves to match the new viability optimum, but in that window of time we observe little change in 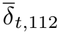 (fig. 6B), and therefore no temporary increase in the mutation probability (fig. 6C). Only in the longer term, after asymmetries in the trait distribution have had the time to build (e.g., in our simulations, after approximately 50 000 generations), does indirect selection start having a significant effect on the evolution of mutation probabilities. In the absence of cost to fidelity, indirect selection then results in mutation probabilities evolving towards smaller values, although they remain larger than the deterministic expectation. In the presence of costs, a stable intermediate long-term mutation probability is obtained (fig. 6, dashed lines). In that case, deterministic predictions do well at predicting the long-term order of magnitude of mutation probabilities observed in the simulations, but fail at predicting the observed values precisely, which are consistently lower than expected.

Some of the observed discrepancies between our deterministic predictions and observed values in individual-based simulations can be explained by the effect of genetic drift. In particular, a slower accumulation of asymmetries in smaller populations is expected since genetic drift is known to reduce genetic (co)variation (Lande 1979; Bürger 1993; Week 2026). In the simulations, the observed absence of temporary increases in mutation probabilities is likely a consequence of this effect: the average viability trait reaches equilibrium before any substantial associations can be formed. This agrees with two observations: (i) the accumulation of coskewness is even weaker in smaller populations (see fig. S2) and (ii) compensating for this effect of drift by increasing the mutational input of variation in trait 2 in the simulations (by setting 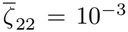, which accelerates the build-up of 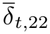 via eq. 19 and thereby strengthens the production of coskewness in eq. 27) qualitatively recovers the trajectories predicted by the deterministic model with 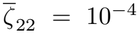 (i.e., short increases in mutation probabilities during adaptation; see fig. S3).

Genetic drift can also help explain discrepancies between observed and predicted long-term mutation probabilities. In the absence of costs to fidelity, indirect selection on mutation rates is known to become too weak to act below a threshold that depends on population size (Lynch 2010; Gervais & Roze 2017). Our observation that simulated mutation probabilities seem to plateau clearly above zero is consistent with this, as is the observation that this plateau is higher under smaller population size (see fig. S2). Interestingly, while in the absence of a cost to fidelity weaker indirect selection leads to larger-than-expected mutation probabilities, in the presence of a cost the observed mutation probabilities are consistently smaller than expected, a pattern which weaker indirect selection alone would not produce. A possible explanation is that there exists an additional selective effect against high mutation probabilities in finite populations (see Raynes et al. 2018). Understanding the interaction between the effects of drift and selection any further is beyond the scope of this work.

Note that we also explored the case where strong competition results in diversifying selection on the viability trait once the population has reached *z*_op_, leading to branching and to the long-term coexistence of multiple morphs. Simulations in this case suggest that such a selection regime can maintain intermediate mutation probabilities even in the absence of costs to fidelity (see fig. S1). This observation however cannot be fully explained by our model, since branching and increasing variance quickly violate the small-variance assumption, and for that reason we do not attempt a detailed description of that case here.

#### 4.4.2 Fluctuating selection increases mutation probabilities on the long term

The simulations described above showed no temporary increase in mutation probabilities during the directional selection phase, partly because that phase was relatively brief. There are, however, cases where directional selection acts during longer periods, for instance when the fitness landscape changes periodically or when species coevolve. To illustrate these cases, we focus on the simple case of an environment regularly switching between two states. We initiate the population in the same way as above, but the trait optimum now changes from 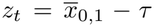 to 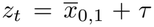 every *T* generations. The population thus experiences alternating episodes of directional (when en route to the current trait optimum) and homogenising selection (when the current optimum is close). Here we arbitrarily set *τ* = 0.05 and *γ* = 0.3 throughout, such that evolving from one optimum to the next is relatively quick, and we can observe many alternating selection phases within a reasonable time frame. These values (together with *η* = 10^*−*2^) also ensure that the fitness landscape between the optima is never convex (i.e., mutations remain deleterious on average), such that none of the observations can be due to indirect diversifying selection. We vary *T* to vary the relative time spent in each regime by the population. To isolate these effects from those of direct selection, we set the cost to fidelity to zero *α* = 0. The exact relationship between *τ, T* and mutation probabilities is a complicated unknown function of most other variables (since they set the pace of evolution and thus the relative time spent in each regime) that we do not attempt to characterise here. Our goal is only to show some of the possible consequences of mixed selection regimes on long-term mutation probabilities.

When this scenario is iterated numerically (fig. 7, lines), we find that alternating selection regimes can lead to a long-term stable intermediate mutation probability. More precisely, they can make the mutation probability oscillate (directional phases push it up and homogenising phases push it down) within an intermediate range that settles in the long term (fig. 7C). At the start of each new directional phase, selection quickly associates mutator trait values with the new most successful variants, leading to large oscillations in covariance (between positive and negative values; fig. 7D). In that case, decreasing *T* leads to more frequent episodes of directional selection, and thus to higher observed long-term mutation probabilities.

**Fig. 7:**
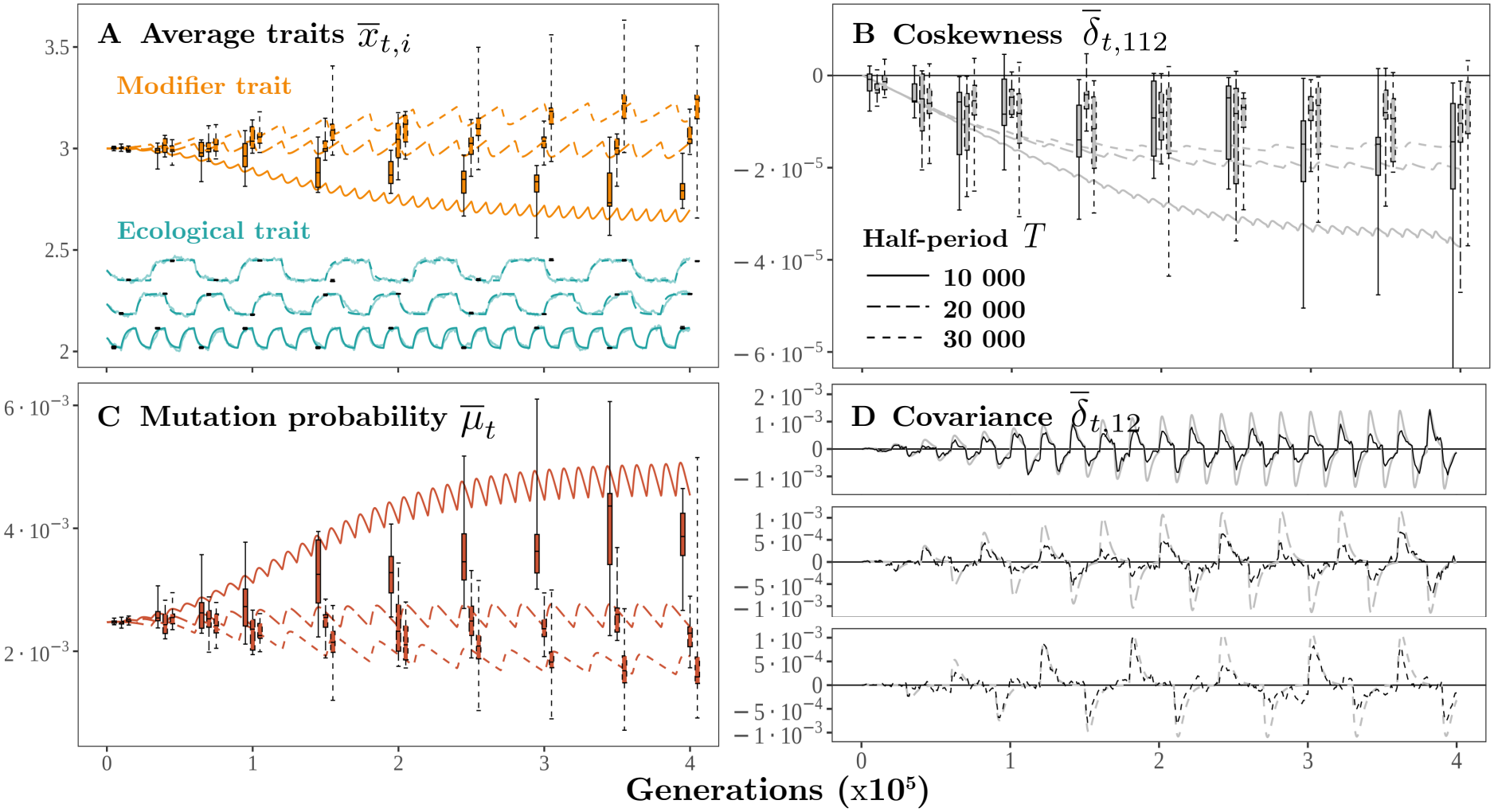
Evolutionary dynamics in a fluctuating environment. The viability optimum alternates between two values 0.1 trait units apart, changing every *T* generations (line types: half-period *T* ). Near the current optimum, selection on the viability trait is homogenising. Mean viability trait (blue) and modifier trait (orange). **B**. Two-way coskewness between the modifier trait and variation at the viability trait. **C**. Mean mutation rate, showing a stable long-term range despite no cost to fidelity (*α* = 0). **D**. Genetic covariance between the viability and modifier traits. Lines show deterministic predictions obtained by iterating the system of moments; boxplots give the median, interquartile range and total range across 10 independent individual-based simulations. Other fixed parameters: *β* = 2; *λ* = 3; *γ* = 0.3; 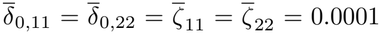; *η* = 0.01; *ρ* = 10^*−*5^, leading to a population of *∼* 2 *×* 10^5^ individuals. **Alt text:** Time series for an optimum that alternates between two nearby values. Panel A shows the mean viability and modifier traits, panel B two-way coskewness, panel C mean mutation rate and panel D covariance. Lines give deterministic predictions for several half-periods, and boxplots give 10 individual-based simulations. Mean mutation rate remains within a stable range over the long term.

When simulating a finite population under the same conditions (fig. 7, boxplots), we recover the same qualitative results, albeit with some delay due to the longer time required for associations between traits to build up. After enough coskewness has accumulated, episodes of directional selection following environmental change do lead to temporary associations between advantageous trait values and mutagenic trait values (fig. 7D). This drives temporary increases in the mutation probability, which compensate for the decreases resulting from the reduction principle near the optimum, leading to a stable range of observed mutation probabilities in the long term. As in the transient case, while numerical iterations predict the right order of magnitude for mutation probabilities, and the right relationship with *T*, they fail at predicting the exact range of long-term mutation probabilities. The simulated long-term ranges are lower than the predicted ranges on average (fig. 7C), although individual simulations can overshoot leading to rates higher than expected (especially in smaller populations; see fig. S4). Unless finite population sizes disfavour high mutation probabilities generally (Raynes et al. 2018), this suggests that the hitch-hiking effect is more vulnerable to small population size than indirect homogenising selection, in agreement with previous descriptions (e.g., Tenaillon et al. 2000). We leave the exploration of the complex interaction between genetic drift on the one hand, and the tug-of-war between pathways of indirect selection on the other, to future work.

## 5 Discussion

We introduced a formalisation of the role of natural selection in the evolution of mutation rates, in which locus-specific mutation probabilities evolve through their dependence on quantitative traits. Unlike more classical modifier models (Liberman & Feldman 1986; Gervais & Roze 2017; Shen et al. 2020), which follow the frequencies of a finite number of alleles, we track a multivariate trait distribution, as in quantitative genetics (Lande 1979), using the continuum-of-alleles model (Kimura 1965; Bürger 1998; Waxman 2003) under a small-variance assumption. Allowing for infinitely many possible alleles comes at a cost: indirect selection on mutation rates then depends on third-order associations, and devising a closed system tracking these associations requires an additional closure assumption (i.e., neglecting fourth- and fifth-order cumulants). In exchange, the resulting description provides a hierarchical account of several selective mechanisms acting on mutation rates, in terms of moments of the trait distribution (i.e., variational properties of the population), derivatives of the fitness landscape (the selection regime), and derivatives of the mutation probabilities (the effect of traits on these rates), each with a biological interpretation. This clarifies the role of ecology (i.e., present and past fitness landscapes) and functional trade-offs in shaping mutation rates over short and long timescales.

With this approach, separate known effects of selection on mutation rates can be recovered and compared. First, the reduction principle (Liberman & Feldman 1986; Altenberg et al. 2017), often framed as selection against linked genetic load, is understood here as the result of indirect homogenising selection at a viability optimum, enabled by associations between modifier trait values and the variation they produce at viability traits (i.e., two-way coskewness). In this framing, variation is equivalent to genetic load because a concave fitness landscape implies that extreme viability trait values are deleterious. Indirect selection against such load favours smaller mutation rates, and may balance opposing direct effects (e.g., cost to fidelity). If extreme trait values are favoured by a convex landscape instead, our results imply that selection favours an increase in mutation rates, and variation can be viewed as genetic *potential* instead. One particular weakness of our approach is that it ceases to produce predictions in that latter case, since strong variation violates our working assumptions. Nonetheless, the promotion of intermediate mutation rates by prolonged regimes of diversifying selection is supported by our individual-based simulations (see fig. S1).

Another, separate case where more frequent mutations can be favoured is when directional trait change increases fitness, such that some specific alleles are under positive selection. In this case, selection for increased mutation rates is often described as mutator alleles hitch-hiking with the successful variants they generate (e.g., Eshel 1973; Taddei et al. 1997b; Johnson 1999b). Here, we frame it as indirect directional selection on mutagenic trait values, via their association (i.e., covariance) with favourable trait values, and show that this association is expected to be generated by the combination of variable mutation rates and directional selection on viability traits. Because this mechanism relies on first-order selection, it can outweigh second-order selection against load when the landscape is steep enough for long enough (i.e., given that the required associations can be built), for instance under fluctuating selection as in our example model. This confirms that selection can maintain intermediate mutation rates even if the landscape remains concave, that is when mutations are still deleterious on average, and even in the absence of direct costs to fidelity or other functional trade-offs. This extends previous findings (Ishii et al. 1989; Travis & Travis 2002; Carja et al. 2014; Rosenbloom & Allen 2014) to a more general ecological scenario, and removes the technical requirement of a separation of timescales between the modifying and the modified locus.

One class of problems that our approach can help tackle is that involving several modifying loci and/or several modified loci. For instance, the case explored by Ishii et al. (1989) of a modifier of two loci under different regimes could be extended to many loci and mixed selection regimes, asking what selection pressures act on genome-wide modifying traits such as DNA-polymerase quality or error-correcting mechanisms. In the future, such analyses could help further determine whether variation in selection regimes across genomic regions could explain some of the observed variation in genome-wide mutation rates (Lynch et al. 2023). Our approach could also help study the selective mechanisms at play in observed cases of local hyper-mutability, at loci experiencing fluctuating selective pressures (Barnett et al. 2025; Fyon & Úbeda 2025). In particular, it could help generalise these mechanisms, asking whether and when we should observe increased mutation rates at other types of loci that undergo frequent directional or diversifying selection (e.g., underlying competitive or immunity traits). Finally, it could participate in explaining differences across species with different histories of selection, asking whether and why specific evolutionary events (e.g., domestication, adaptive radiations) should be expected to affect mutation rates.

Our results rely on a small-variance and a cumulant-neglect assumption. Beyond the case of diversifying selection discussed above, predicted dynamics are thus only reliable insofar as (co)variances and (co)skewnesses (and the variation they imply) remain small, and as long as the neglected cumulants never play an important role. To go further and extend our results to cases involving adaptive polymorphism might be possible with the appropriate methods (Sasaki & Dieckmann 2011; Lion et al. 2023). In addition, even if trait variation does remain small, comparing numerical iterations and individual-based simulations shows that our model cannot be expected to produce precise quantitative predictions in finite populations (see figs. 6 and 7). This is in part because genetic drift erodes genetic variation and associations, thus weakening each of the described selective mechanisms to different extents. In the future, moment-based descriptions of the interplay between regimes of selection and genetic drift in the evolution of mutation rates could be obtained by using mathematical tools developed to study genetic drift in similar models (Rice 2020; Week 2026).

It is important to stress that several common biological processes neglected here have important roles in the evolution of mutation rates. First and foremost, recombination is known to reduce genetic associations across loci and therefore the scope for indirect selection (Bürger 2000; Altenberg et al. 2017; Gervais & Roze 2017). Since our model assumes non-recombining haplotypes, it applies primarily to non-sexual species with small genomes, or to cases where mutation modifiers are tightly linked to the modified locus (e.g., when the modifier is the structure of the genetic locus itself; Barnett et al. 2025; Fyon & Úbeda 2025). Second, stochastic trait expression (e.g., due to environmental variation) also blurs the association between expressed traits and fitness (an effect measured by heritability; Lande 1976), and thus must affect indirect selection, in ways that remain unclear. More generally, any modification to the map between sequence-level information and expressed traits necessarily affects the mutational targets of these traits, i.e., how often and how much they change by mutation. This means that the genetic architecture of viability traits (e.g., regulation pathways, ploidy) is a core determinant of the role and value of genetic mutation, an architecture itself composed of evolvable traits (e.g., regulation parameters). How mutation effects evolve through the evolution of the genotype-phenotype map is still an open question, which must be taken into account when comparing sequence-level mutation rates between organisms with largely different maps. The methodology presented here could be extended to study the coevolution of genotype-phenotype maps and mutation parameters under different selection regimes, improving our understanding of the links between ecology, functional trade-offs, and transmission strategies.

## Data availability

A Mathematica notebook containing the symbolic derivation of the moment equations, their numerical iteration, and the code producing the illustrative panels (Supplementary Mathematica Notebook 1), and the Rust code of the individual-based simulations (Supplementary Archive 1), are available at zenodo.org (DOI: 10.5281/zenodo.24368121).

## Acknowledgements

We used Claude Sonnet 5.5 (Anthropic, 2026) for help writing code and text. The authors reviewed and tested all outputs and take full responsibility for the content of the manuscript and supplementary files.

## 6 Supplementary tables and figures

**Tab. S1:**
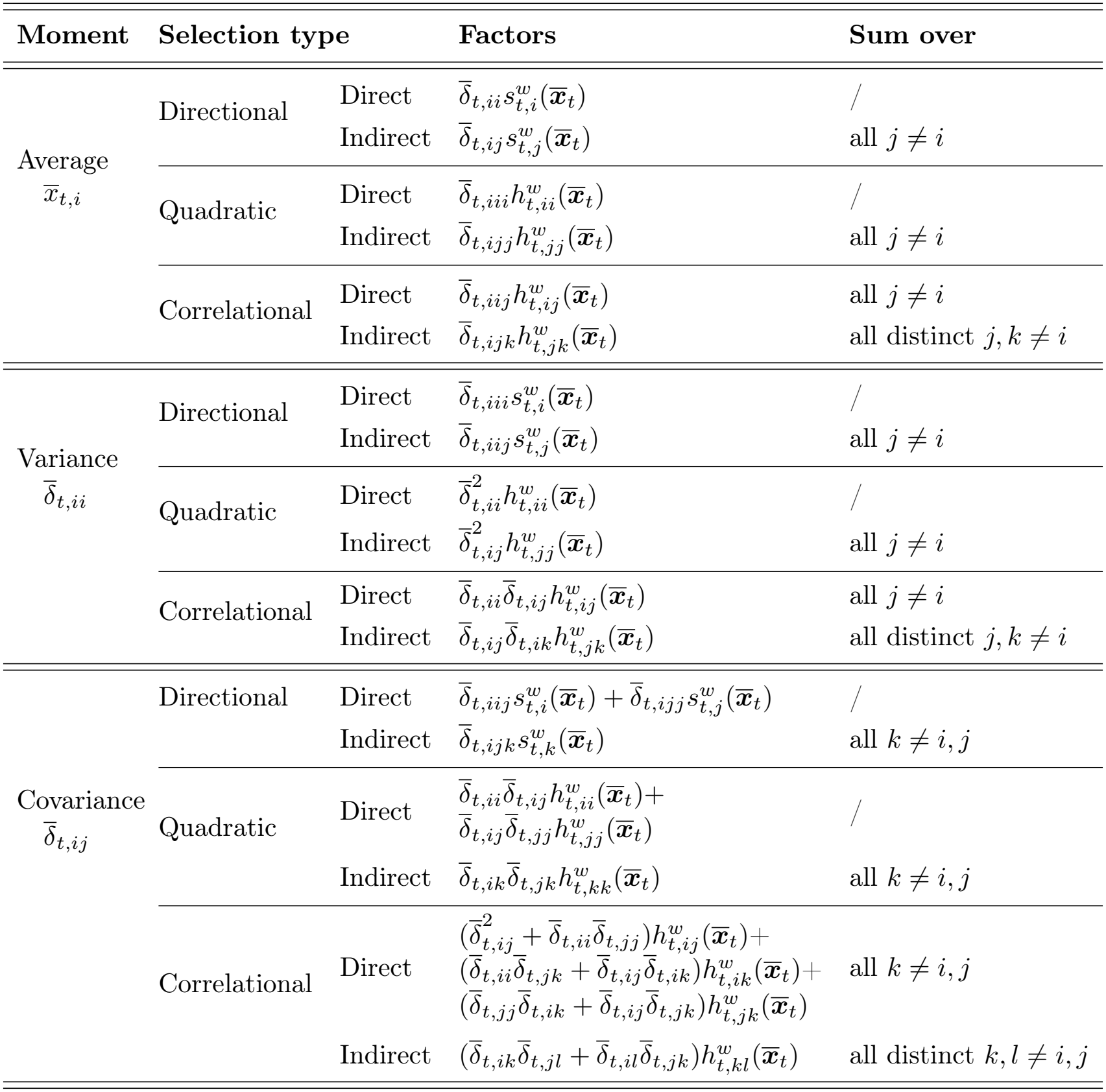
Types of selection acting on trait means and (co)variances. The table gives selection terms associated with the effects of each selection type of each moment. The factor 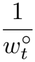, and integer factors due to summing and halving (see eqs. 11, 13, 16 and 17) are omitted from all terms for simplicity.

| Moment | Selection type |  | Factors | Sum over |
|---|---|---|---|---|
| Average<br>$\overline{x}_{t,i}$ | Directional | Direct | $\overline{\delta}_{t,ii}s^{w}_{t,i}(\overline{\boldsymbol{x}}_{t})$ | / |
|  |  | Indirect | $\overline{\delta}_{t,ij}s^{w}_{t,j}(\overline{\boldsymbol{x}}_{t})$ | all $j \neq i$ |
|  | Quadratic | Direct | $\overline{\delta}_{t,iii}h^{w}_{t,ii}(\overline{\boldsymbol{x}}_{t})$ | / |
|  |  | Indirect | $\overline{\delta}_{t,ijj}h^{w}_{t,jj}(\overline{\boldsymbol{x}}_{t})$ | all $j \neq i$ |
|  | Correlational | Direct | $\overline{\delta}_{t,iij}h^{w}_{t,ij}(\overline{\boldsymbol{x}}_{t})$ | all $j \neq i$ |
|  |  | Indirect | $\overline{\delta}_{t,ijk}h^{w}_{t,jk}(\overline{\boldsymbol{x}}_{t})$ | all distinct $j, k \neq i$ |
| Variance<br>$\overline{\delta}_{t,ii}$ | Directional | Direct | $\overline{\delta}_{t,iii}s^{w}_{t,i}(\overline{\boldsymbol{x}}_{t})$ | / |
|  |  | Indirect | $\overline{\delta}_{t,iij}s^{w}_{t,j}(\overline{\boldsymbol{x}}_{t})$ | all $j \neq i$ |
|  | Quadratic | Direct | $\overline{\delta}^{2}_{t,ii}h^{w}_{t,ii}(\overline{\boldsymbol{x}}_{t})$ | / |
|  |  | Indirect | $\overline{\delta}^{2}_{t,ij}h^{w}_{t,jj}(\overline{\boldsymbol{x}}_{t})$ | all $j \neq i$ |
|  | Correlational | Direct | $\overline{\delta}_{t,ii}\overline{\delta}_{t,ij}h^{w}_{t,ij}(\overline{\boldsymbol{x}}_{t})$ | all $j \neq i$ |
|  |  | Indirect | $\overline{\delta}_{t,ij}\overline{\delta}_{t,ik}h^{w}_{t,jk}(\overline{\boldsymbol{x}}_{t})$ | all distinct $j, k \neq i$ |
| Covariance<br>$\overline{\delta}_{t,ij}$ | Directional | Direct | $\overline{\delta}_{t,iij}s^{w}_{t,i}(\overline{\boldsymbol{x}}_{t}) + \overline{\delta}_{t,ijj}s^{w}_{t,j}(\overline{\boldsymbol{x}}_{t})$ | / |
|  |  | Indirect | $\overline{\delta}_{t,ijk}s^{w}_{t,k}(\overline{\boldsymbol{x}}_{t})$ | all $k \neq i, j$ |
|  | Quadratic | Direct | $\overline{\delta}_{t,ii}\overline{\delta}_{t,ij}h^{w}_{t,ii}(\overline{\boldsymbol{x}}_{t})+$<br>$\overline{\delta}_{t,ij}\overline{\delta}_{t,jj}h^{w}_{t,jj}(\overline{\boldsymbol{x}}_{t})$ | / |
|  |  | Indirect | $\overline{\delta}_{t,ik}\overline{\delta}_{t,jk}h^{w}_{t,kk}(\overline{\boldsymbol{x}}_{t})$ | all $k \neq i, j$ |
|  | Correlational | Direct | $(\overline{\delta}^{2}_{t,ij} + \overline{\delta}_{t,ii}\overline{\delta}_{t,jj})h^{w}_{t,ij}(\overline{\boldsymbol{x}}_{t})+$<br>$(\overline{\delta}_{t,ii}\overline{\delta}_{t,jk} + \overline{\delta}_{t,ij}\overline{\delta}_{t,ik})h^{w}_{t,ik}(\overline{\boldsymbol{x}}_{t})+$<br>$(\overline{\delta}_{t,jj}\overline{\delta}_{t,ik} + \overline{\delta}_{t,ij}\overline{\delta}_{t,jk})h^{w}_{t,jk}(\overline{\boldsymbol{x}}_{t})$ | all $k \neq i, j$ |
|  |  | Indirect | $(\overline{\delta}_{t,ik}\overline{\delta}_{t,jl} + \overline{\delta}_{t,il}\overline{\delta}_{t,jk})h^{w}_{t,kl}(\overline{\boldsymbol{x}}_{t})$ | all distinct $k, l \neq i, j$ |

**Tab. S2:**
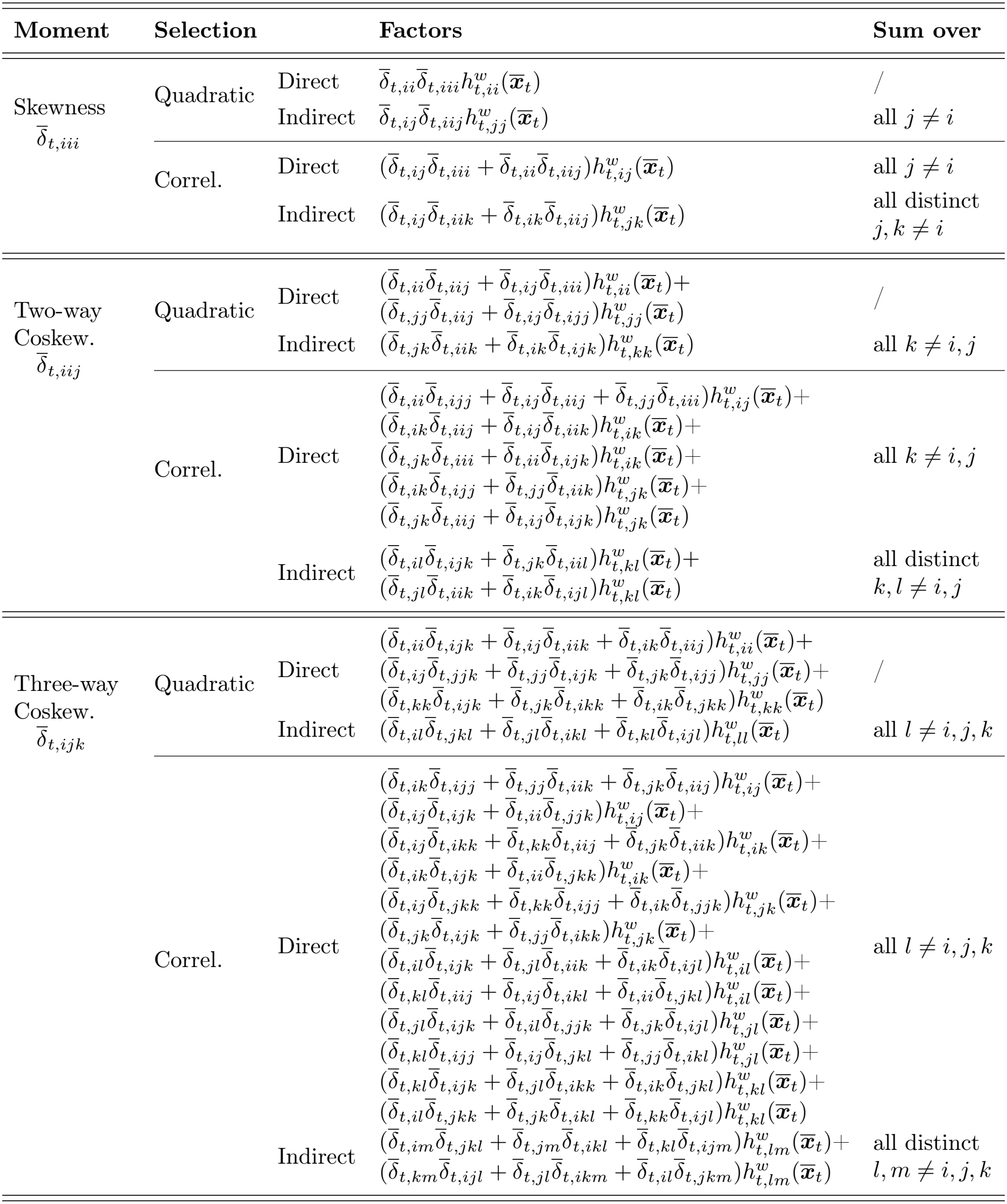
Types of selection acting on trait (co)skewnesses. The table gives selection terms associated with the effects of each selection type of each moment. The factor 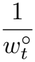, and integer factors due to summing and halving (see eq. 24) are omitted from all terms for simplicity.

| Moment | Selection |  | Factors | Sum over |
|---|---|---|---|---|
| Skewness<br>$\overline{\delta}_{t,iii}$ | Quadratic | Direct | $\overline{\delta}_{t,ii}\overline{\delta}_{t,iii}h^{w}_{t,ii}(\overline{\boldsymbol{x}}_t)$ | / |
|  |  | Indirect | $\overline{\delta}_{t,ij}\overline{\delta}_{t,iij}h^{w}_{t,jj}(\overline{\boldsymbol{x}}_t)$ | all $j \neq i$ |
|  | Correl. | Direct | $(\overline{\delta}_{t,ij}\overline{\delta}_{t,iii} + \overline{\delta}_{t,ii}\overline{\delta}_{t,iij})h^{w}_{t,ij}(\overline{\boldsymbol{x}}_t)$ | all $j \neq i$ |
|  |  | Indirect | $(\overline{\delta}_{t,ij}\overline{\delta}_{t,iik} + \overline{\delta}_{t,ik}\overline{\delta}_{t,iij})h^{w}_{t,jk}(\overline{\boldsymbol{x}}_t)$ | all distinct<br>$j,k \neq i$ |
| Two-way<br>Coskew.<br>$\overline{\delta}_{t,iij}$ | Quadratic | Direct | $(\overline{\delta}_{t,ii}\overline{\delta}_{t,iij} + \overline{\delta}_{t,ij}\overline{\delta}_{t,iii})h^{w}_{t,ii}(\overline{\boldsymbol{x}}_t)+$<br>$(\overline{\delta}_{t,jj}\overline{\delta}_{t,iij} + \overline{\delta}_{t,ij}\overline{\delta}_{t,ijj})h^{w}_{t,jj}(\overline{\boldsymbol{x}}_t)$ | / |
|  |  | Indirect | $(\overline{\delta}_{t,jk}\overline{\delta}_{t,iik} + \overline{\delta}_{t,ik}\overline{\delta}_{t,ijk})h^{w}_{t,kk}(\overline{\boldsymbol{x}}_t)$ | all $k \neq i,j$ |
|  | Correl. | Direct | $(\overline{\delta}_{t,ii}\overline{\delta}_{t,ijj} + \overline{\delta}_{t,ij}\overline{\delta}_{t,iij} + \overline{\delta}_{t,jj}\overline{\delta}_{t,iii})h^{w}_{t,ij}(\overline{\boldsymbol{x}}_t)+$<br>$(\overline{\delta}_{t,ik}\overline{\delta}_{t,iij} + \overline{\delta}_{t,ij}\overline{\delta}_{t,iik})h^{w}_{t,ik}(\overline{\boldsymbol{x}}_t)+$<br>$(\overline{\delta}_{t,jk}\overline{\delta}_{t,iii} + \overline{\delta}_{t,ii}\overline{\delta}_{t,ijk})h^{w}_{t,ik}(\overline{\boldsymbol{x}}_t)+$<br>$(\overline{\delta}_{t,ik}\overline{\delta}_{t,ijj} + \overline{\delta}_{t,jj}\overline{\delta}_{t,iik})h^{w}_{t,jk}(\overline{\boldsymbol{x}}_t)+$<br>$(\overline{\delta}_{t,jk}\overline{\delta}_{t,iij} + \overline{\delta}_{t,ij}\overline{\delta}_{t,ijk})h^{w}_{t,jk}(\overline{\boldsymbol{x}}_t)$ | all $k \neq i,j$ |
|  |  | Indirect | $(\overline{\delta}_{t,il}\overline{\delta}_{t,ijk} + \overline{\delta}_{t,jk}\overline{\delta}_{t,iil})h^{w}_{t,kl}(\overline{\boldsymbol{x}}_t)+$<br>$(\overline{\delta}_{t,jl}\overline{\delta}_{t,iik} + \overline{\delta}_{t,ik}\overline{\delta}_{t,ijl})h^{w}_{t,kl}(\overline{\boldsymbol{x}}_t)$ | all distinct<br>$k,l \neq i,j$ |
| Three-way<br>Coskew.<br>$\overline{\delta}_{t,ijk}$ | Quadratic | Direct | $(\overline{\delta}_{t,ii}\overline{\delta}_{t,ijk} + \overline{\delta}_{t,ij}\overline{\delta}_{t,iik} + \overline{\delta}_{t,ik}\overline{\delta}_{t,iij})h^{w}_{t,ii}(\overline{\boldsymbol{x}}_t)+$<br>$(\overline{\delta}_{t,ij}\overline{\delta}_{t,jjk} + \overline{\delta}_{t,jj}\overline{\delta}_{t,ijk} + \overline{\delta}_{t,jk}\overline{\delta}_{t,ijj})h^{w}_{t,jj}(\overline{\boldsymbol{x}}_t)+$<br>$(\overline{\delta}_{t,kk}\overline{\delta}_{t,ijk} + \overline{\delta}_{t,jk}\overline{\delta}_{t,ikk} + \overline{\delta}_{t,ik}\overline{\delta}_{t,jkk})h^{w}_{t,kk}(\overline{\boldsymbol{x}}_t)$ | / |
|  |  | Indirect | $(\overline{\delta}_{t,il}\overline{\delta}_{t,jkl} + \overline{\delta}_{t,jl}\overline{\delta}_{t,ikl} + \overline{\delta}_{t,kl}\overline{\delta}_{t,ijl})h^{w}_{t,ll}(\overline{\boldsymbol{x}}_t)$ | all $l \neq i,j,k$ |
|  | Correl. | Direct | $(\overline{\delta}_{t,ik}\overline{\delta}_{t,ijj} + \overline{\delta}_{t,jj}\overline{\delta}_{t,iik} + \overline{\delta}_{t,jk}\overline{\delta}_{t,iij})h^{w}_{t,ij}(\overline{\boldsymbol{x}}_t)+$<br>$(\overline{\delta}_{t,ij}\overline{\delta}_{t,ijk} + \overline{\delta}_{t,ii}\overline{\delta}_{t,jjk})h^{w}_{t,ij}(\overline{\boldsymbol{x}}_t)+$<br>$(\overline{\delta}_{t,ij}\overline{\delta}_{t,ikk} + \overline{\delta}_{t,kk}\overline{\delta}_{t,iij} + \overline{\delta}_{t,jk}\overline{\delta}_{t,iik})h^{w}_{t,ik}(\overline{\boldsymbol{x}}_t)+$<br>$(\overline{\delta}_{t,ik}\overline{\delta}_{t,ijk} + \overline{\delta}_{t,ii}\overline{\delta}_{t,jkk})h^{w}_{t,ik}(\overline{\boldsymbol{x}}_t)+$<br>$(\overline{\delta}_{t,ij}\overline{\delta}_{t,jkk} + \overline{\delta}_{t,kk}\overline{\delta}_{t,ijj} + \overline{\delta}_{t,ik}\overline{\delta}_{t,jjk})h^{w}_{t,jk}(\overline{\boldsymbol{x}}_t)+$<br>$(\overline{\delta}_{t,jk}\overline{\delta}_{t,ijk} + \overline{\delta}_{t,jj}\overline{\delta}_{t,ikk})h^{w}_{t,jk}(\overline{\boldsymbol{x}}_t)+$<br>$(\overline{\delta}_{t,il}\overline{\delta}_{t,ijk} + \overline{\delta}_{t,jl}\overline{\delta}_{t,iik} + \overline{\delta}_{t,ik}\overline{\delta}_{t,ijl})h^{w}_{t,il}(\overline{\boldsymbol{x}}_t)+$<br>$(\overline{\delta}_{t,kl}\overline{\delta}_{t,iij} + \overline{\delta}_{t,ij}\overline{\delta}_{t,ikl} + \overline{\delta}_{t,ii}\overline{\delta}_{t,jkl})h^{w}_{t,il}(\overline{\boldsymbol{x}}_t)+$<br>$(\overline{\delta}_{t,jl}\overline{\delta}_{t,ijk} + \overline{\delta}_{t,il}\overline{\delta}_{t,jjk} + \overline{\delta}_{t,jk}\overline{\delta}_{t,ijl})h^{w}_{t,jl}(\overline{\boldsymbol{x}}_t)+$<br>$(\overline{\delta}_{t,kl}\overline{\delta}_{t,ijj} + \overline{\delta}_{t,ij}\overline{\delta}_{t,jkl} + \overline{\delta}_{t,jj}\overline{\delta}_{t,ikl})h^{w}_{t,jl}(\overline{\boldsymbol{x}}_t)+$<br>$(\overline{\delta}_{t,kl}\overline{\delta}_{t,ijk} + \overline{\delta}_{t,jl}\overline{\delta}_{t,ikk} + \overline{\delta}_{t,ik}\overline{\delta}_{t,jkl})h^{w}_{t,kl}(\overline{\boldsymbol{x}}_t)+$<br>$(\overline{\delta}_{t,il}\overline{\delta}_{t,jkk} + \overline{\delta}_{t,jk}\overline{\delta}_{t,ikl} + \overline{\delta}_{t,kk}\overline{\delta}_{t,ijl})h^{w}_{t,kl}(\overline{\boldsymbol{x}}_t)$ | all $l \neq i,j,k$ |
|  |  | Indirect | $(\overline{\delta}_{t,im}\overline{\delta}_{t,jkl} + \overline{\delta}_{t,jm}\overline{\delta}_{t,ikl} + \overline{\delta}_{t,kl}\overline{\delta}_{t,ijm})h^{w}_{t,lm}(\overline{\boldsymbol{x}}_t)+$<br>$(\overline{\delta}_{t,km}\overline{\delta}_{t,ijl} + \overline{\delta}_{t,jl}\overline{\delta}_{t,ikm} + \overline{\delta}_{t,il}\overline{\delta}_{t,jkm})h^{w}_{t,lm}(\overline{\boldsymbol{x}}_t)$ | all distinct<br>$l,m \neq i,j,k$ |

**Fig. S1:**
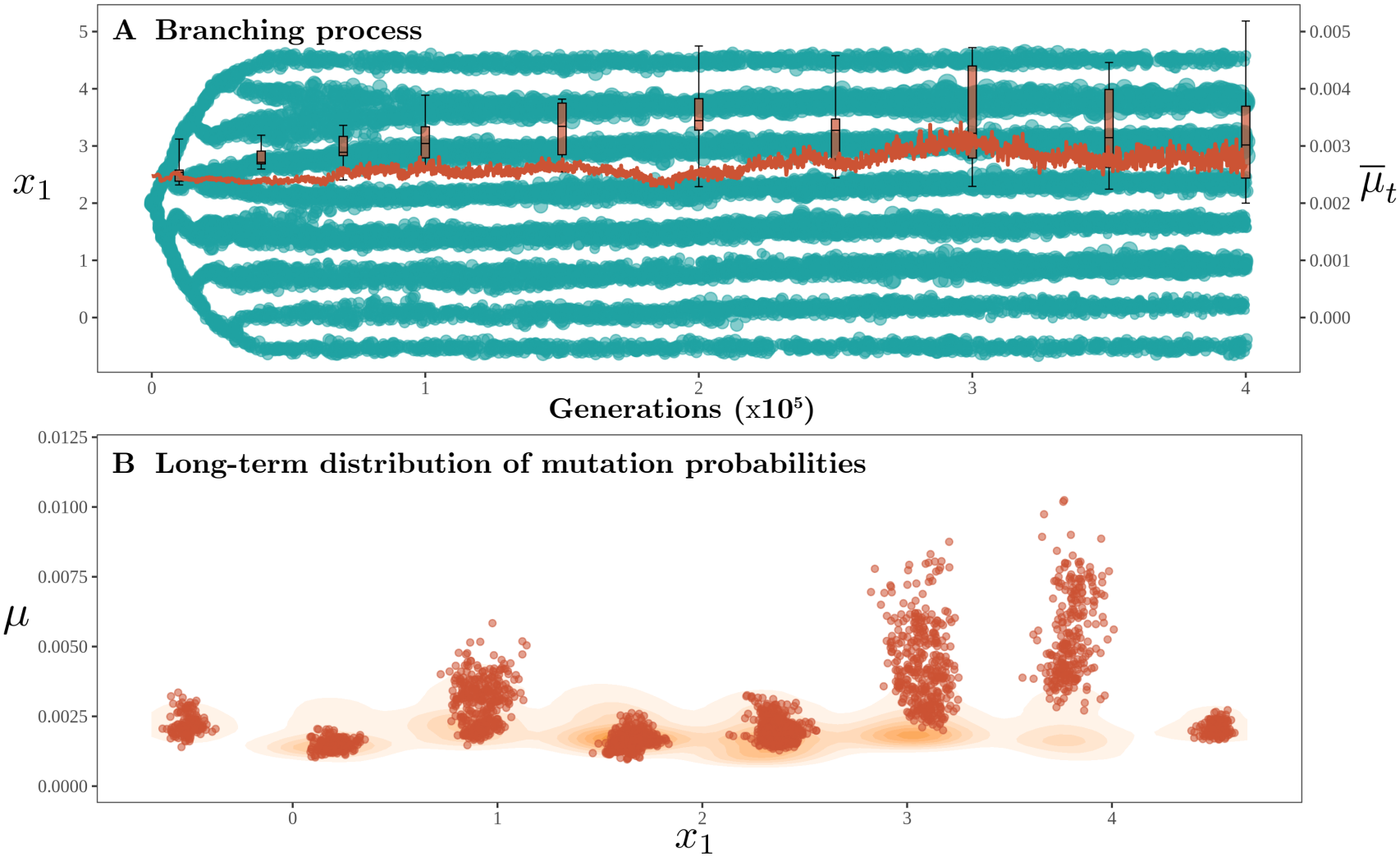
Evolutionary dynamics under strong diversifying selection. The population begins with 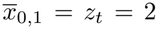 and 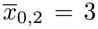, and evolves for 400 000 generations. Competition is strong (*η* = 2) such that selection is diversifying, and leads to successive branching events. Mutation probabilities are maintained positive, close to their initial values. **A**. Values of the viability trait (blue points; left axis) and the average mutation probability (red line; right axis) over time in one individual simulation chosen at random. Boxplots give the median, interquartile range, and total range of average mutation probabilities across 10 independent simulations. **B**. The distribution of average trait values and mutation probabilities in individuals sampled in later generations (350 000–400 000), in one individual simulation chosen at random (points) and across 10 independent simulations (background colours represent density). Other fixed parameters: *β* = 2; *λ* = 3; *γ* = 0.1; 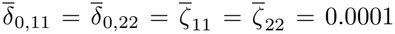; *α* = 0; *ρ* = 10^*−*5^, leading to a population of *∼* 2 *×* 10^5^ individuals.

**Fig. S2:**
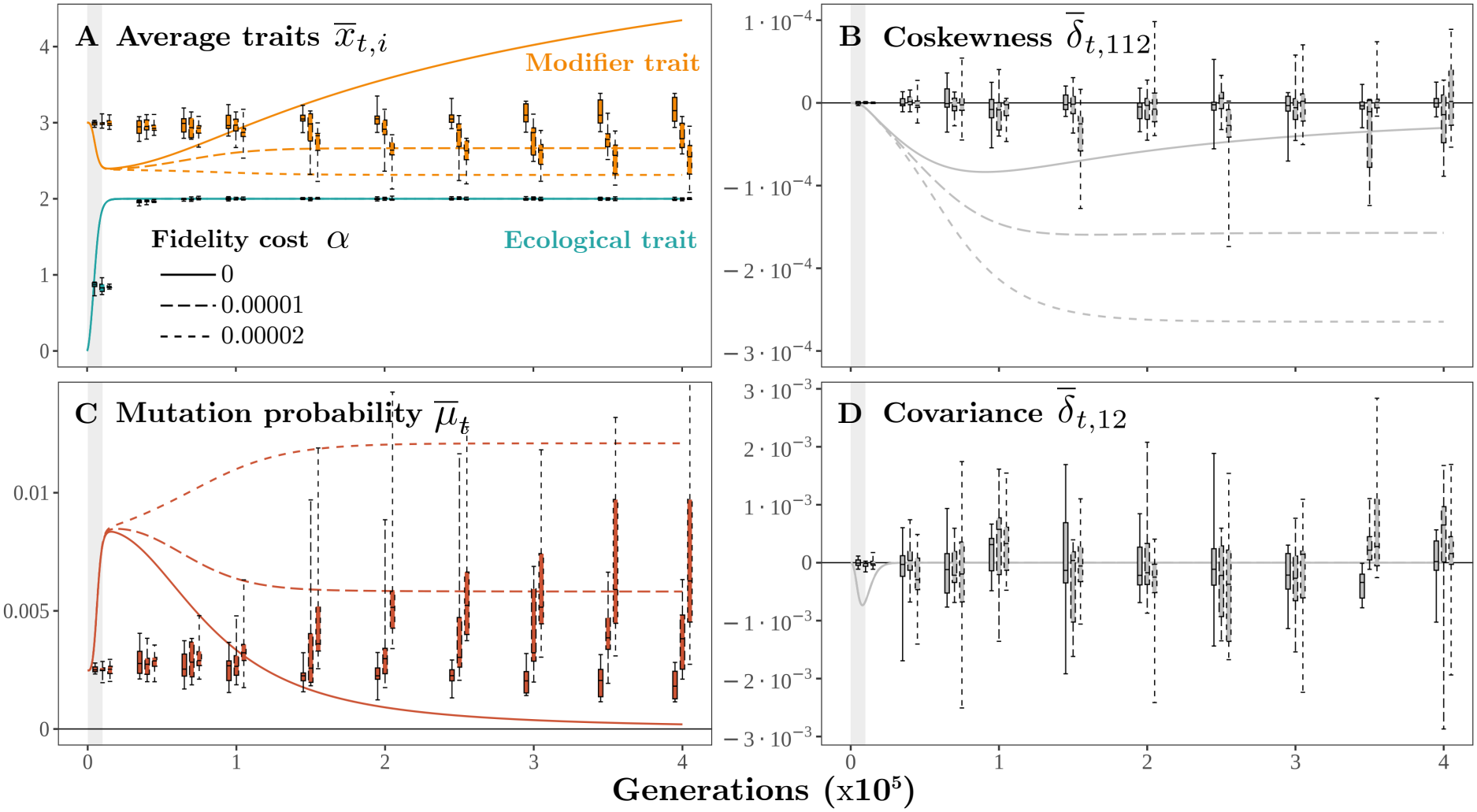
Transient evolutionary dynamics following environmental change, in a smaller population. The population begins with an average viability trait (trait 1) placed two trait units away from the viability optimum *z*_*t*_ = 2, and evolves for 400 000 generations. After *∼* 10000 generations of directional selection (marked by a grey shading), the population reaches the optimum, where it experiences constant homogenising selection. The modifier trait (trait 2) controls the mutation rate at both loci under varying costs (line types indicate the fidelity cost *α*). **A**. Mean value of the viability trait (blue) and the modifier trait (orange) over time. **B**. Two-way coskewness measuring the association between the modifier trait and variation at the viability trait. **C**. Mean mutation rate. **D**. Genetic covariance between the viability and modifier traits. In all panels, lines show deterministic predictions obtained by iterating the system of moments. Boxplots give the median, interquartile range, and total range of each statistic across 10 independent individual-based simulations. Other fixed parameters: *β* = 2; *λ* = 3; *γ* = 0.1; 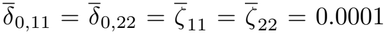; *η* = 0.01; *ρ* = 10^*−*4^, leading to a population of *∼* 2 *×* 10^4^ individuals.

**Fig. S3:**
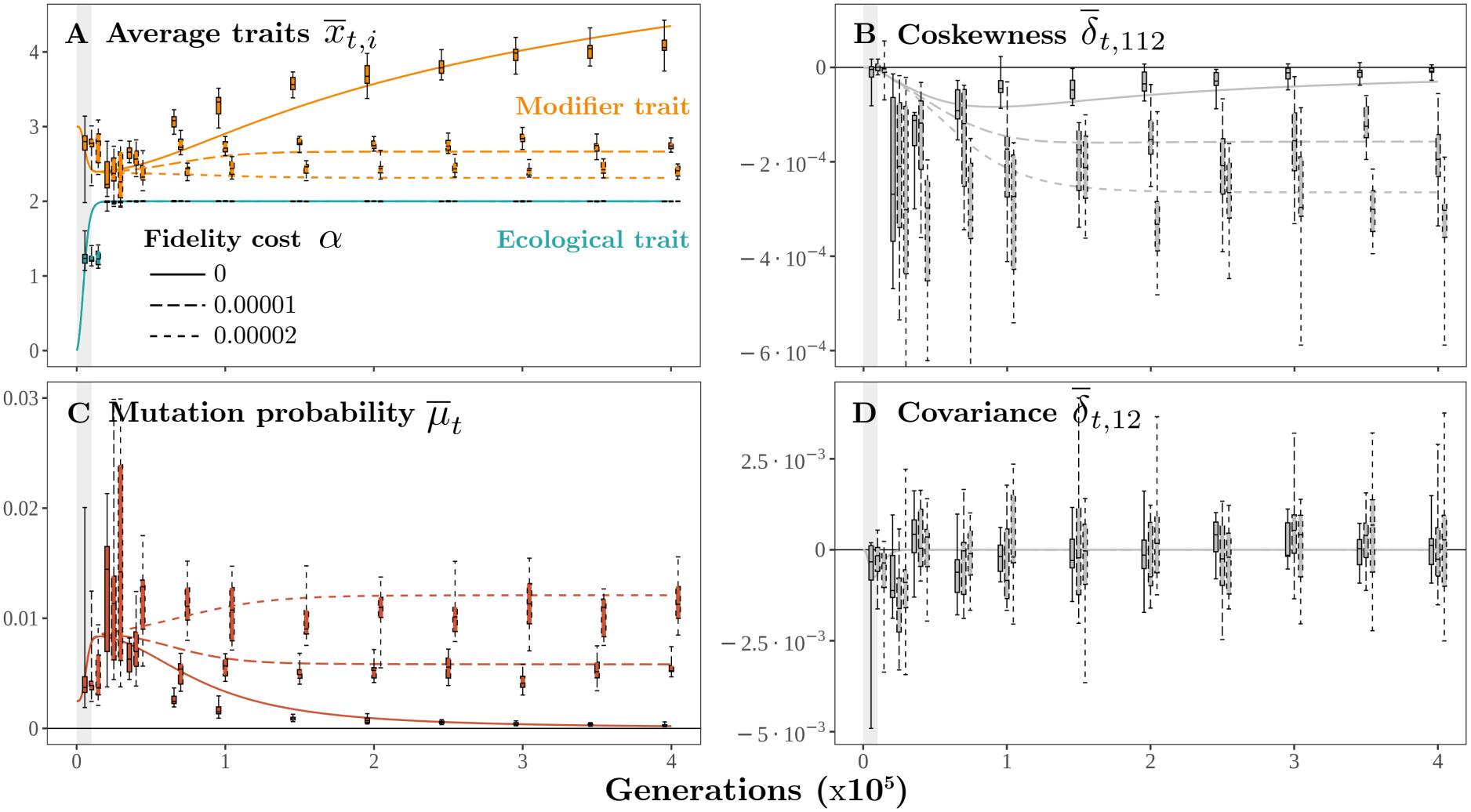
Transient evolutionary dynamics following environmental change, with large mutational variance on trait 2. The population begins with an average viability trait (trait 1) placed two trait units away from the viability optimum *z*_*t*_ = 2, and evolves for 400 000 generations. After *∼* 10000 generations of directional selection (marked by a grey shading), the population reaches the optimum, where it experiences constant homogenising selection. The modifier trait (trait 2) controls the mutation rate at both loci under varying costs (line types indicate the fidelity cost *α*). **A**. Mean value of the viability trait (blue) and the modifier trait (orange) over time. **B**. Two-way coskewness measuring the association between the modifier trait and variation at the viability trait. **C**. Mean mutation rate. **D**. Genetic covariance between the viability and modifier traits. In all panels, lines show deterministic predictions obtained by iterating the system of moments with 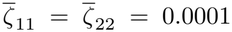, as in the main text. Boxplots, however, give the median, interquartile range, and total range of each statistic across 10 independent individual-based simulations obtained with 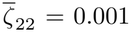. This change makes the evolution of trait 2 faster such that it “catches up” to deterministic expectations, and shows large increases in the mutation rate during adaptation. Other fixed parameters: *β* = 2; *λ* = 3; *γ* = 0.1; 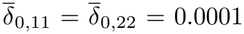; *η* = 0.01; *ρ* = 10^*−*4^, leading to a population of *∼* 2 *×* 10^4^ individuals.

**Fig. S4:**
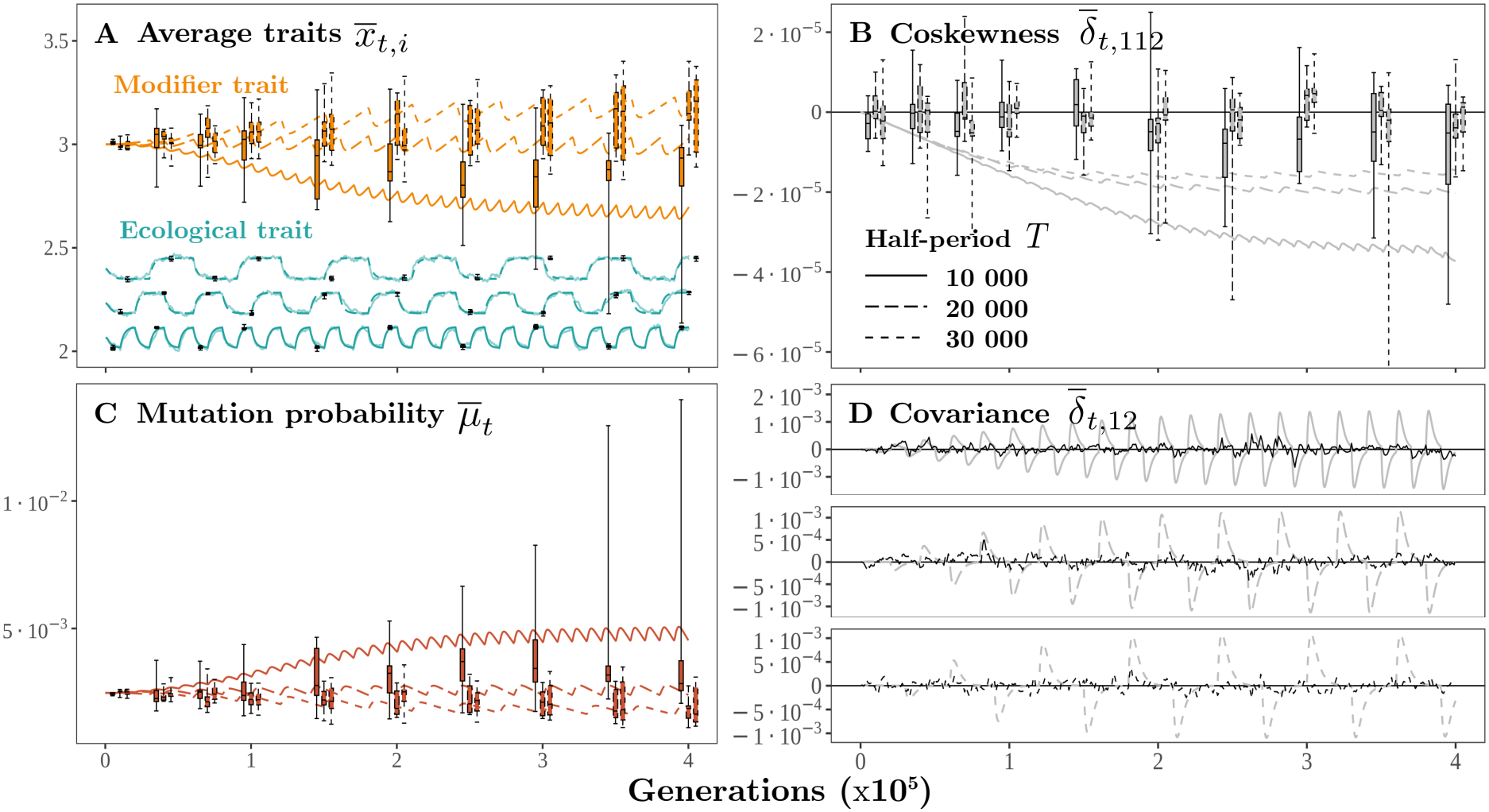
Evolutionary dynamics in a fluctuating environment, in a smaller population. The viability optimum alternates between two nearby values separated by 0.1 trait units, changing every *T* generations (line type indicates the value of the half-period *T* ). Near the current optimum, selection on the viability trait is homogenising. **A**. Mean value of the viability trait (blue) and the modifier trait (orange) over time. **B**. Two-way coskewness measuring the association between the modifier trait and variation at the viability trait. **C**. Mean mutation rate, showing a stable long-term range (despite no cost to fidelity, as here *α* = 0). **D**. Genetic covariance between the viability and modifier traits. In all panels, lines show deterministic predictions obtained by iterating the system of moments. Boxplots give the median, interquartile range, and total range of each statistic across 10 independent individual-based simulations. Other fixed parameters: *β* = 2; *λ* = 3; *γ* = 0.3; 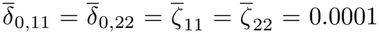; *η* = 0.01; *ρ* = 10^*−*4^, leading to a population of *∼* 2 *×* 10^4^ individuals.

