## Appendices for "How gradual evolution, fitness landscapes, and trait associations shape mutation rates in large populations"

**Appendix A - Trait dynamics in terms of moment-generating functions** **56**

### Appendix A - Trait dynamics in terms of moment-generating functions

In this appendix, we derive exact expressions for the change in average traits, trait (co)variances and trait (co)skewnesses. These expressions form the basis of the approximations presented in the main text, and derived in the next appendices.

#### A.1 Multivariate moment-generating functions

We start by deriving a general moment-generating function (MGF) representation for the trait distribution, which is useful to compute the dynamics of its centred moments.

##### A.1.1 Centred MGF

By definition, the MGF of the distribution of traits at generation $t$, when centred around the current average trait, is

$$\mathcal{G}_t(\boldsymbol{\theta}) = \int_{\mathbb{R}^{n_x}} e^{\boldsymbol{\theta}\cdot(\boldsymbol{x}-\overline{\boldsymbol{x}}_t)} p_t(\boldsymbol{x}) \mathrm{d}\boldsymbol{x}, \tag{A-1}$$

where $\boldsymbol{\theta}\cdot\boldsymbol{x} = \sum_{i=1}^{n_x} \theta_i x_i$. This function entails that

$$\mathcal{G}_t(\mathbf{0}) = 1, \quad \left.\frac{\partial \mathcal{G}_t(\boldsymbol{\theta})}{\partial \theta_i}\right|_{\boldsymbol{\theta}=\mathbf{0}} = 0, \quad \left.\frac{\partial^2 \mathcal{G}_t(\boldsymbol{\theta})}{\partial \theta_i \partial \theta_j}\right|_{\boldsymbol{\theta}=\mathbf{0}} = \overline{\delta}_{t,ij}, \quad \left.\frac{\partial^3 \mathcal{G}_t(\boldsymbol{\theta})}{\partial \theta_i \partial \theta_j \partial \theta_k}\right|_{\boldsymbol{\theta}=\mathbf{0}} = \overline{\delta}_{t,ijk}. \tag{A-2}$$

With $\Delta\overline{\boldsymbol{x}}_t = \overline{\boldsymbol{x}}_{t+1} - \overline{\boldsymbol{x}}_t$ the MGF at generation $t+1$ can thus be written as

$$\begin{aligned}\mathcal{G}_{t+1}(\boldsymbol{\theta}) &= \int_{\mathbb{R}^{n_x}} e^{\boldsymbol{\theta}\cdot(\boldsymbol{x}-\overline{\boldsymbol{x}}_{t+1})} p_{t+1}(\boldsymbol{x}) \mathrm{d}\boldsymbol{x} \\ &= e^{-\boldsymbol{\theta}\cdot\Delta\overline{\boldsymbol{x}}_t} \mathcal{H}_{t+1}(\boldsymbol{\theta}),\end{aligned} \tag{A-3}$$

where

$$\mathcal{H}_{t+1}(\boldsymbol{\theta}) = \int_{\mathbb{R}^{n_x}} e^{\boldsymbol{\theta}\cdot(\boldsymbol{x}-\overline{\boldsymbol{x}}_t)} p_{t+1}(\boldsymbol{x}) \mathrm{d}\boldsymbol{x} \tag{A-4}$$

is the MGF of the distribution of traits at generation $t+1$, but centred around the parental average trait. The first-order derivative of $\mathcal{H}_{t+1}(\boldsymbol{\theta})$ gives the new average trait relative to the parental average, that is

$$\left.\frac{\partial \mathcal{H}_{t+1}(\boldsymbol{\theta})}{\partial \theta_i}\right|_{\boldsymbol{\theta}=\mathbf{0}} = \overline{x}_{t+1,i} - \overline{x}_{t,i} = \Delta\overline{x}_{t,i}. \tag{A-5}$$

Therefore, eqs. (A-3) and (A-4) are useful to track the means, (co)variances, and (co)skewnesses of the trait distribution.

##### A.1.2 Explicit MGF

We next express eq. (A-4) in terms of quantities in the parental generation so as to be able to describe changes over time. To that end we substitute eq. (6) into eq. (A-4) and use $\boldsymbol{\zeta} = \boldsymbol{y} - \boldsymbol{x}$ to denote the vector of parent-offspring differences, giving

$$\begin{aligned}
\mathcal{H}_{t+1}(\boldsymbol{\theta}) &= \frac{1}{\overline{w}_t} \int_{\mathbb{R}^{n_x}} \int_{\mathbb{R}^{n_x}} e^{\boldsymbol{\theta}\cdot(\boldsymbol{y}-\overline{\boldsymbol{x}}_t)} \pi(\boldsymbol{y}-\boldsymbol{x}|\boldsymbol{x}) w_t(\boldsymbol{x}, \boldsymbol{\phi}_t(\boldsymbol{x})) p_t(\boldsymbol{x}) \mathrm{d}\boldsymbol{y} \mathrm{d}\boldsymbol{x}, \\
&= \frac{1}{\overline{w}_t} \int_{\mathbb{R}^{n_x}} e^{\boldsymbol{\theta}\cdot(\boldsymbol{x}-\overline{\boldsymbol{x}}_t)} \left[ \int_{\mathbb{R}^{n_x}} e^{\boldsymbol{\theta}\cdot(\boldsymbol{y}-\boldsymbol{x})} \pi(\boldsymbol{y}-\boldsymbol{x}|\boldsymbol{x}) \mathrm{d}\boldsymbol{y} \right] w_t(\boldsymbol{x}, \boldsymbol{\phi}_t(\boldsymbol{x})) p_t(\boldsymbol{x}) \mathrm{d}\boldsymbol{x} \\
&= \frac{1}{\overline{w}_t} \int_{\mathbb{R}^{n_x}} e^{\boldsymbol{\theta}\cdot(\boldsymbol{x}-\overline{\boldsymbol{x}}_t)} \left[ \int_{\mathbb{R}^{n_x}} e^{\boldsymbol{\theta}\cdot\boldsymbol{\zeta}} \pi(\boldsymbol{\zeta}|\boldsymbol{x}) \mathrm{d}\boldsymbol{\zeta} \right] w_t(\boldsymbol{x}, \boldsymbol{\phi}_t(\boldsymbol{x})) p_t(\boldsymbol{x}) \mathrm{d}\boldsymbol{x} \\
&= \frac{1}{\overline{w}_t} \int_{\mathbb{R}^{n_x}} e^{\boldsymbol{\theta}\cdot(\boldsymbol{x}-\overline{\boldsymbol{x}}_t)} \mathcal{U}(\boldsymbol{\theta} \mid \boldsymbol{x}) w_t(\boldsymbol{x}, \boldsymbol{\phi}_t(\boldsymbol{x})) p_t(\boldsymbol{x}) \mathrm{d}\boldsymbol{x},
\end{aligned} \tag{A-6}$$

where

$$\mathcal{U}(\boldsymbol{\theta} \mid \boldsymbol{x}) = \int_{\mathbb{R}^{n_x}} e^{\boldsymbol{\theta}\cdot\boldsymbol{\zeta}} \pi(\boldsymbol{\zeta}|\boldsymbol{x}) \, \mathrm{d}\boldsymbol{\zeta} \tag{A-7}$$

is the conditional MGF of the distribution of parent-offspring differences. Substituting eq. (3) therein gives

$$\begin{aligned}
\mathcal{U}(\boldsymbol{\theta} \mid \boldsymbol{x}) &= \int_{\mathbb{R}^{n_x}} e^{\boldsymbol{\theta}\cdot\boldsymbol{\zeta}} \prod_{i=1}^{n_x} \left[ (1-\mu_i(\boldsymbol{x}))\delta(\zeta_i) + \mu_i(\boldsymbol{x}) q_i(\zeta_i) \right] \, \mathrm{d}\boldsymbol{\zeta} \\
&= \int_{\mathbb{R}^{n_x}} \prod_{i=1}^{n_x} \left\{ e^{\theta_i \zeta_i} \left[ (1-\mu_i(\boldsymbol{x}))\delta(\zeta_i) + \mu_i(\boldsymbol{x}) q_i(\zeta_i) \right] \right\} \, \mathrm{d}\zeta_1 \cdots \mathrm{d}\zeta_{n_x}.
\end{aligned} \tag{A-8}$$

The integrand in eq. (A-8) is a product across traits and the $n_x$-dimensional integral can thus be written as a product of one-dimensional integrals,

$$\begin{aligned}
\mathcal{U}(\boldsymbol{\theta} \mid \boldsymbol{x}) &= \prod_{i=1}^{n_x} \int_{\mathbb{R}} e^{\theta_i \zeta_i} \left[ (1-\mu_i(\boldsymbol{x}))\delta(\zeta_i) + \mu_i(\boldsymbol{x}) q_i(\zeta_i) \right] \, \mathrm{d}\zeta_i \\
&= \prod_{i=1}^{n_x} \left[ (1-\mu_i(\boldsymbol{x})) \int_{\mathbb{R}} e^{\theta_i \zeta_i} \delta(\zeta_i) \, \mathrm{d}\zeta_i + \mu_i(\boldsymbol{x}) \int_{\mathbb{R}} e^{\theta_i \zeta_i} q_i(\zeta_i) \, \mathrm{d}\zeta_i \right].
\end{aligned} \tag{A-9}$$

For trait $i$, now define the MGF of mutational effects at locus $i$,

$$\mathcal{M}_i(\theta_i) = \int_{\mathbb{R}} e^{\theta_i \zeta_i} q_i(\zeta_i) \, \mathrm{d}\zeta_i. \tag{A-10}$$

Since $\int_{\mathbb{R}} e^{\theta_i \zeta_i} \delta(\zeta_i) \, \mathrm{d}\zeta_i = 1$, eq. (A-9) becomes

$$\begin{aligned}
\mathcal{U}(\boldsymbol{\theta} \mid \boldsymbol{x}) &= \prod_{i=1}^{n_x} \left[ (1-\mu_i(\boldsymbol{x})) + \mu_i(\boldsymbol{x}) \mathcal{M}_i(\theta_i) \right] \\
&= \prod_{i=1}^{n_x} \left[ 1 + \mu_i(\boldsymbol{x}) (\mathcal{M}_i(\theta_i) - 1) \right].
\end{aligned} \tag{A-11}$$

Substituting into eq. (A-6) and using $\boldsymbol{x} = \overline{\boldsymbol{x}}_t + \boldsymbol{\delta}_t$ yields

$$\begin{aligned}\mathcal{H}_{t+1}(\boldsymbol{\theta}) &= \frac{1}{\overline{w}_t}\mathbb{E}_t\left[e^{\boldsymbol{\theta}\cdot\boldsymbol{\delta}_t}\,\mathcal{U}(\boldsymbol{\theta} \mid \overline{\boldsymbol{x}}_t + \boldsymbol{\delta}_t)w_t(\overline{\boldsymbol{x}}_t + \boldsymbol{\delta}_t, \boldsymbol{\phi}_t(\overline{\boldsymbol{x}}_t + \boldsymbol{\delta}_t))\right] \\ &= \frac{1}{\overline{w}_t}\mathbb{E}_t\left[e^{\boldsymbol{\theta}\cdot\boldsymbol{\delta}_t}\prod_{i=1}^{n_x}\left[1 + \mu_i(\overline{\boldsymbol{x}}_t + \boldsymbol{\delta}_t)(\mathcal{M}_i(\theta_i) - 1)\right] w_t(\overline{\boldsymbol{x}}_t + \boldsymbol{\delta}_t, \boldsymbol{\phi}_t(\overline{\boldsymbol{x}}_t + \boldsymbol{\delta}_t))\right],\end{aligned} \tag{A-12}$$

where

$$\mathbb{E}_t[a_t] = \overline{a}_t = \int_{\mathbb{R}^{n_x}} a_t(\boldsymbol{x})p_t(\boldsymbol{x})\mathrm{d}\boldsymbol{x} \tag{A-13}$$

denotes the expectation of a function $a_t : \mathbb{R}^{n_x} \to \mathbb{R}$ (which can depend on time) over the distribution of parental trait values.

Let

$$\mathcal{Q}_{t+1}(\boldsymbol{\theta}) = \frac{1}{\overline{w}_t}\mathbb{E}_t\left[e^{\boldsymbol{\theta}\cdot\boldsymbol{\delta}_t}w_t(\overline{\boldsymbol{x}}_t + \boldsymbol{\delta}_t, \boldsymbol{\phi}_t(\overline{\boldsymbol{x}}_t + \boldsymbol{\delta}_t))\right] \tag{A-14}$$

stand for the MGF of the trait distribution after selection only. In addition, for any non-empty subset $S \subseteq \{1, \ldots, n_x\}$ of loci, define the MGF of the trait after (normalised) mutation (including its interaction with selection) by

$$\mathcal{V}_{t+1,S}(\boldsymbol{\theta}) = \frac{1}{\overline{w}_t}\mathbb{E}_t\left[e^{\boldsymbol{\theta}\cdot\boldsymbol{\delta}_t}w_t(\overline{\boldsymbol{x}}_t + \boldsymbol{\delta}_t, \boldsymbol{\phi}_t(\overline{\boldsymbol{x}}_t + \boldsymbol{\delta}_t))\mu_S(\boldsymbol{x})\right], \tag{A-15}$$

where

$$\mu_S(\boldsymbol{x}) = \prod_{i\in S}\mu_i(\overline{\boldsymbol{x}}_t + \boldsymbol{\delta}_t) \tag{A-16}$$

is the joint mutation rate of the subset $S$, i.e., the probability that the loci listed in $S$ mutate jointly in a single reproduction event. With this, eq. (A-12) becomes

$$\mathcal{H}_{t+1}(\boldsymbol{\theta}) = \mathcal{Q}_{t+1}(\boldsymbol{\theta}) + \sum_{\substack{S\subseteq\{1,\ldots,n_x\}\\ S\neq\emptyset}}\left(\prod_{i\in S}[\mathcal{M}_i(\theta_i) - 1]\right)\mathcal{V}_{t+1,S}(\boldsymbol{\theta}), \tag{A-17}$$

where the product between parentheses accounts for the distribution of mutation effects, and $\emptyset$ denotes the empty set. Eq. (A-17) separates the effects of selection and mutation: the first term is the MGF after selection only, and each term of the sum accounts for the effect of simultaneous mutations at the loci in $S$, i.e., including events where several mutations co-occur (when the size of $S$ is more than one). In the next subsection we show that, up to third-order moments, only single-locus mutation events matter.

##### A.1.3 MGF under symmetric mutations

Under symmetric mutations (i.e., $q_i(\zeta_i) = q_i(-\zeta_i)$), eq. (A-10) entails that

$$\mathcal{M}_i(0) = 1, \quad \left.\frac{\partial\mathcal{M}_i(\theta_i)}{\partial\theta_i}\right|_{\theta_i=0} = 0, \quad \left.\frac{\partial^2\mathcal{M}_i(\theta_i)}{\partial\theta_i^2}\right|_{\theta_i=0} = \overline{\zeta}_{ii}, \quad \left.\frac{\partial^3\mathcal{M}_i(\theta_i)}{\partial\theta_i^3}\right|_{\theta_i=0} = 0. \tag{A-18}$$

Eq. (A-18) implies that each mutational factor in eq. (A-17) (i.e., of the form $(\mathcal{M}_i(\theta_i) - 1)$) must receive two derivatives to survive. Since we will distribute at most three derivatives, this means that most terms in the sum in eq. (A-17) will not survive. We now rearrange eq. (A-17) to isolate the terms that will cancel under isotropy. For this, let $S_k$ denote a subset of size $k$ (with $1 \leq k \leq n_x$). This lets us separate the terms in $\mathcal{H}_{t+1}(\boldsymbol{\theta})$ corresponding to zero or one mutation event from those corresponding to two or more as

$$\mathcal{H}_{t+1}(\boldsymbol{\theta}) = \tilde{\mathcal{H}}_{t+1}(\boldsymbol{\theta}) + \sum_{k=2}^{n_x} \sum_{S_k \subseteq \{1,\ldots,n_x\}} \left( \prod_{i \in S_k} [\mathcal{M}_i(\theta_i) - 1] \right) \mathcal{V}_{t+1,S_k}(\boldsymbol{\theta}), \tag{A-19}$$

with

$$\tilde{\mathcal{H}}_{t+1}(\boldsymbol{\theta}) = \mathcal{Q}_{t+1}(\boldsymbol{\theta}) + \sum_{i=1}^{n_x} [\mathcal{M}_i(\theta_i) - 1] \mathcal{V}_{t+1,i}(\boldsymbol{\theta}), \tag{A-20}$$

where we used the short-hand $\mathcal{V}_{t+1,i}(\boldsymbol{\theta}) = \mathcal{V}_{t+1,\{i\}}(\boldsymbol{\theta})$.

Every summand in the last term of eq. (A-19) has two or more mutational factors (i.e., $(\mathcal{M}_i(\theta_i) - 1)(\mathcal{M}_j(\theta_j) - 1)\ldots$). For this sum to survive differentiation, each summand must therefore receive at least four derivatives. Up to third-order derivatives, this is impossible and thus,

$$\left.\frac{\partial \mathcal{H}_{t+1}(\boldsymbol{\theta})}{\partial \theta_i}\right|_{\boldsymbol{\theta}=\mathbf{0}} = \left.\frac{\partial \tilde{\mathcal{H}}_{t+1}(\boldsymbol{\theta})}{\partial \theta_i}\right|_{\boldsymbol{\theta}=\mathbf{0}}, \tag{A-21}$$

$$\left.\frac{\partial^2 \mathcal{H}_{t+1}(\boldsymbol{\theta})}{\partial \theta_i \partial \theta_j}\right|_{\boldsymbol{\theta}=\mathbf{0}} = \left.\frac{\partial^2 \tilde{\mathcal{H}}_{t+1}(\boldsymbol{\theta})}{\partial \theta_i \partial \theta_j}\right|_{\boldsymbol{\theta}=\mathbf{0}}, \tag{A-22}$$

and

$$\left.\frac{\partial^3 \mathcal{H}_{t+1}(\boldsymbol{\theta})}{\partial \theta_i \partial \theta_j \partial \theta_k}\right|_{\boldsymbol{\theta}=\mathbf{0}} = \left.\frac{\partial^3 \tilde{\mathcal{H}}_{t+1}(\boldsymbol{\theta})}{\partial \theta_i \partial \theta_j \partial \theta_k}\right|_{\boldsymbol{\theta}=\mathbf{0}}. \tag{A-23}$$

Hence, under symmetric mutations, computing the change in trait moments up to third order only requires locus-specific mutational generating functions.

#### A.2 Total change in trait moments

We now use the moment-generating functions described above to express the change in the moments of interest.

##### A.2.1 Average traits

According to eqs. (A-5), (A-20) and (A-21), we have

$$\begin{aligned}\Delta\overline{x}_{t,i} &= \overline{x}_{t+1,i} - \overline{x}_{t,i} \\ &= \left.\frac{\partial\mathcal{H}_{t+1}(\boldsymbol{\theta})}{\partial\theta_i}\right|_{\boldsymbol{\theta}=\mathbf{0}} \\ &= \left.\frac{\partial\tilde{\mathcal{H}}_{t+1}(\boldsymbol{\theta})}{\partial\theta_i}\right|_{\boldsymbol{\theta}=\mathbf{0}} \\ &= \left.\frac{\partial\mathcal{Q}_{t+1}(\boldsymbol{\theta})}{\partial\theta_i}\right|_{\boldsymbol{\theta}=\mathbf{0}}.\end{aligned} \tag{A-24}$$

The last equality in eq. (A-24) obtains due to our assumption of symmetric mutations (eq. A-18), which imply no effect of mutation on average traits. Differentiating eq. (A-14) gives

$$\left.\frac{\partial\mathcal{Q}_{t+1}(\boldsymbol{\theta})}{\partial\theta_i}\right|_{\boldsymbol{\theta}=\mathbf{0}} = \frac{1}{\overline{w}_t}\mathbb{E}_t[w_t\delta_{t,i}] = \frac{1}{\overline{w}_t}\mathrm{Cov}_t\big[w_t, \delta_{t,i}\big], \tag{A-25}$$

where

$$\begin{aligned}\mathrm{Cov}_t\big[a_t, b_t\big] &= \mathbb{E}_t\big[(a_t(\boldsymbol{x}) - \overline{a}_t)(b_t(\boldsymbol{x}) - \overline{b}_t)\big] \\ &= \mathbb{E}_t[a_t b_t] - \overline{a}_t\overline{b}_t\end{aligned} \tag{A-26}$$

denotes the covariance between two functions $a_t$ and $b_t$ under the distribution of trait values at generation $t$. Therefore

$$\Delta\overline{x}_{t,i} = \frac{1}{\overline{w}_t}\mathrm{Cov}_t\big[w_t, \delta_{t,i}\big]. \tag{A-27}$$

##### A.2.2 Trait (co)variances

According to eqs. (A-2), (A-3), (A-22) and (A-24) we have

$$\begin{aligned}\Delta\overline{\delta}_{t,ij} &= \overline{\delta}_{t+1,ij} - \overline{\delta}_{t,ij} \\ &= \left.\frac{\partial^2\mathcal{G}_{t+1}(\boldsymbol{\theta})}{\partial\theta_i\partial\theta_j}\right|_{\boldsymbol{\theta}=\mathbf{0}} - \overline{\delta}_{t,ij} \\ &= \left.\frac{\partial^2\tilde{\mathcal{H}}_{t+1}(\boldsymbol{\theta})}{\partial\theta_i\partial\theta_j}\right|_{\boldsymbol{\theta}=\mathbf{0}} + \Delta\overline{x}_{t,i}\Delta\overline{x}_{t,j} - \Delta\overline{x}_{t,i}\left.\frac{\partial\tilde{\mathcal{H}}_{t+1}(\boldsymbol{\theta})}{\partial\theta_j}\right|_{\boldsymbol{\theta}=\mathbf{0}} - \left.\frac{\partial\tilde{\mathcal{H}}_{t+1}(\boldsymbol{\theta})}{\partial\theta_i}\right|_{\boldsymbol{\theta}=\mathbf{0}}\Delta\overline{x}_{t,j} - \overline{\delta}_{t,ij} \\ &= \left.\frac{\partial^2\tilde{\mathcal{H}}_{t+1}(\boldsymbol{\theta})}{\partial\theta_i\partial\theta_j}\right|_{\boldsymbol{\theta}=\mathbf{0}} - \Delta\overline{x}_{t,i}\Delta\overline{x}_{t,j} - \overline{\delta}_{t,ij}.\end{aligned} \tag{A-28}$$

Since only second-order derivatives of $\mathcal{M}_k(\theta_k)$ can survive, differentiating eq. (A-20) twice gives

$$\left.\frac{\partial^2\tilde{\mathcal{H}}_{t+1}(\boldsymbol{\theta})}{\partial\theta_i\partial\theta_j}\right|_{\boldsymbol{\theta}=\mathbf{0}} = \left.\frac{\partial^2\mathcal{Q}_{t+1}(\boldsymbol{\theta})}{\partial\theta_i\partial\theta_j}\right|_{\boldsymbol{\theta}=\mathbf{0}} + \Delta_{m+s}\overline{\delta}_{t,ij}, \tag{A-29}$$

where

$$\Delta_{m+s}\overline{\delta}_{t,ij} = \sum_{k=1}^{n_x} \frac{\partial^2 \mathcal{M}_k(\theta_k)}{\partial\theta_i \partial\theta_j}\bigg|_{\boldsymbol{\theta}=\mathbf{0}} \mathcal{V}_{t+1,k}(\mathbf{0}) \tag{A-30}$$

is the change due to mutation and its interaction with selection. Differentiating eq. (A-14) twice, the first term of eq. (A-29) is

$$\frac{\partial^2 \mathcal{Q}_{t+1}(\boldsymbol{\theta})}{\partial\theta_i \partial\theta_j}\bigg|_{\boldsymbol{\theta}=\mathbf{0}} = \Delta_s\overline{\delta}_{t,ij} + \overline{\delta}_{t,ij}, \tag{A-31}$$

where

$$\Delta_s\overline{\delta}_{t,ij} = \frac{1}{\overline{w}_t}\mathrm{Cov}_t\big[w_t, \delta_{t,i}\delta_{t,j}\big]. \tag{A-32}$$

According to eq. (A-15), we also have

$$\mathcal{V}_{t+1,i}(\mathbf{0}) = \frac{1}{\overline{w}_t}\mathbb{E}_t[w_t\mu_i] = \frac{1}{\overline{w}_t}\mathrm{Cov}_t\big[w_t, \mu_i\big] + \overline{\mu}_{t,i}. \tag{A-33}$$

Thus, since $\partial^2\mathcal{M}_i(\theta_i)/\partial\theta_i\partial\theta_i\big|_{\boldsymbol{\theta}=\mathbf{0}} = \overline{\zeta}_{ii}$ (see eq. A-18) and $\partial^2\mathcal{M}_k(\theta_k)/\partial\theta_i\partial\theta_j\big|_{\boldsymbol{\theta}=\mathbf{0}} = 0$ whenever $i \neq j$, $i \neq k$ or $j \neq k$, we can partition the change involving mutation as

$$\Delta_{m+s}\overline{\delta}_{t,ij} = \Delta_m\overline{\delta}_{t,ij} + \Delta_{m\times s}\overline{\delta}_{t,ij} \tag{A-34}$$

where

$$\Delta_m\overline{\delta}_{t,ij} = \begin{cases} \overline{\zeta}_{ii}\overline{\mu}_{t,i} & \text{if } i = j, \\ 0 & \text{if } i \neq j \end{cases} \tag{A-35}$$

and

$$\Delta_{m\times s}\overline{\delta}_{t,ij} = \begin{cases} \overline{\zeta}_{ii}\dfrac{1}{\overline{w}_t}\mathrm{Cov}_t\big[w_t, \mu_i\big] & \text{if } i = j, \\ 0 & \text{if } i \neq j. \end{cases} \tag{A-36}$$

Finally, using eq. (A-27), define

$$\Delta_{s\times s}\overline{\delta}_{t,ij} = -\Delta\overline{x}_{t,i}\Delta\overline{x}_{t,j} = -\frac{1}{\overline{w}_t^2}\mathrm{Cov}_t\big[w_t, \delta_{t,i}\big]\mathrm{Cov}_t\big[w_t, \delta_{t,j}\big]. \tag{A-37}$$

Gathering terms and simplifying, we find that eq. (A-28) reduces to

$$\Delta\overline{\delta}_{t,ij} = \Delta_s\overline{\delta}_{t,ij} + \Delta_{s\times s}\overline{\delta}_{t,ij} + \Delta_m\overline{\delta}_{t,ij} + \Delta_{m\times s}\overline{\delta}_{t,ij}. \tag{A-38}$$

##### A.2.3 Trait (co)skewnesses

According to eqs. (A-2), (A-3), and (A-23), under symmetric mutations we have

$$\begin{aligned}\Delta\overline{\delta}_{t,ijk} &= \overline{\delta}_{t+1,ijk} - \overline{\delta}_{t,ijk}\\ &= \left.\frac{\partial^3\mathcal{G}_{t+1}(\boldsymbol{\theta})}{\partial\theta_i\partial\theta_j\partial\theta_k}\right|_{\boldsymbol{\theta}=\mathbf{0}} - \overline{\delta}_{t,ijk}\\ &= \left.\frac{\partial^3\tilde{\mathcal{H}}_{t+1}(\boldsymbol{\theta})}{\partial\theta_i\partial\theta_j\partial\theta_k}\right|_{\boldsymbol{\theta}=\mathbf{0}} - \Delta\overline{x}_{t,i}\Delta\overline{x}_{t,j}\Delta\overline{x}_{t,k} - \overline{\delta}_{t,ijk}\\ &\quad - \Delta\overline{x}_{t,i}\left.\frac{\partial^2\tilde{\mathcal{H}}_{t+1}(\boldsymbol{\theta})}{\partial\theta_j\partial\theta_k}\right|_{\boldsymbol{\theta}=\mathbf{0}} - \Delta\overline{x}_{t,j}\left.\frac{\partial^2\tilde{\mathcal{H}}_{t+1}(\boldsymbol{\theta})}{\partial\theta_i\partial\theta_k}\right|_{\boldsymbol{\theta}=\mathbf{0}} - \Delta\overline{x}_{t,k}\left.\frac{\partial^2\tilde{\mathcal{H}}_{t+1}(\boldsymbol{\theta})}{\partial\theta_i\partial\theta_j}\right|_{\boldsymbol{\theta}=\mathbf{0}}\\ &\quad + \Delta\overline{x}_{t,i}\Delta\overline{x}_{t,j}\left.\frac{\partial\tilde{\mathcal{H}}_{t+1}(\boldsymbol{\theta})}{\partial\theta_k}\right|_{\boldsymbol{\theta}=\mathbf{0}} + \Delta\overline{x}_{t,i}\Delta\overline{x}_{t,k}\left.\frac{\partial\tilde{\mathcal{H}}_{t+1}(\boldsymbol{\theta})}{\partial\theta_j}\right|_{\boldsymbol{\theta}=\mathbf{0}}\\ &\quad + \Delta\overline{x}_{t,j}\Delta\overline{x}_{t,k}\left.\frac{\partial\tilde{\mathcal{H}}_{t+1}(\boldsymbol{\theta})}{\partial\theta_i}\right|_{\boldsymbol{\theta}=\mathbf{0}}.\end{aligned}\tag{A-39}$$

Substituting eqs. (A-24) and (A-29) into eq. (A-39) gives

$$\begin{aligned}\Delta\overline{\delta}_{t,ijk} = &\left.\frac{\partial^3\tilde{\mathcal{H}}_{t+1}(\boldsymbol{\theta})}{\partial\theta_i\partial\theta_j\partial\theta_k}\right|_{\boldsymbol{\theta}=\mathbf{0}} + 2\Delta\overline{x}_{t,i}\Delta\overline{x}_{t,j}\Delta\overline{x}_{t,k} - \Delta\overline{x}_{t,i}(\Delta_s\overline{\delta}_{t,jk} + \Delta_{m+s}\overline{\delta}_{t,jk} + \overline{\delta}_{t,jk})\\ &- \Delta\overline{x}_{t,j}(\Delta_s\overline{\delta}_{t,ik} + \Delta_{m+s}\overline{\delta}_{t,ik} + \overline{\delta}_{t,ik}) - \Delta\overline{x}_{t,k}(\Delta_s\overline{\delta}_{t,ij} + \Delta_{m+s}\overline{\delta}_{t,ij} + \overline{\delta}_{t,ij}) - \overline{\delta}_{t,ijk}.\end{aligned}\tag{A-40}$$

Since only second-order derivatives of $\mathcal{M}_k(\theta_k)$ can survive, differentiating eq. (A-20) three times gives

$$\left.\frac{\partial^3\tilde{\mathcal{H}}_{t+1}(\boldsymbol{\theta})}{\partial\theta_i\partial\theta_j\partial\theta_k}\right|_{\boldsymbol{\theta}=\mathbf{0}} = \left.\frac{\partial^3\mathcal{Q}_{t+1}(\boldsymbol{\theta})}{\partial\theta_i\partial\theta_j\partial\theta_k}\right|_{\boldsymbol{\theta}=\mathbf{0}} + \tilde{\Delta}_{m+s,i}\overline{\delta}_{t,ijk} + \tilde{\Delta}_{m+s,j}\overline{\delta}_{t,ijk} + \tilde{\Delta}_{m+s,k}\overline{\delta}_{t,ijk},\tag{A-41}$$

where

$$\tilde{\Delta}_{m+s,i}\overline{\delta}_{t,ijk} = \sum_{l=1}^{n_x}\left.\frac{\partial^2[\mathcal{M}_l(\theta_l)-1]}{\partial\theta_j\partial\theta_k}\right|_{\boldsymbol{\theta}=\mathbf{0}}\left.\frac{\partial\mathcal{V}_{t+1,l}(\boldsymbol{\theta})}{\partial\theta_i}\right|_{\boldsymbol{\theta}=\mathbf{0}}.\tag{A-42}$$

Differentiating eq. (A-14) three times, the first term of eq. (A-41) is

$$\left.\frac{\partial^3\mathcal{Q}_{t+1}(\boldsymbol{\theta})}{\partial\theta_i\partial\theta_j\partial\theta_k}\right|_{\boldsymbol{\theta}=\mathbf{0}} = \frac{1}{\overline{w}_t}\mathrm{Cov}_t\big[w_t, \delta_{t,i}\delta_{t,j}\delta_{t,k}\big] + \overline{\delta}_{t,ijk}.\tag{A-43}$$

In turn, to facilitate simplifications below, we use eqs. (A-30) and (A-33) to re-write eq. (A-42) as

$$\begin{aligned}\tilde{\Delta}_{m+s,i}\overline{\delta}_{t,ijk} &= \sum_{l=1}^{n_x}\left.\frac{\partial^2[\mathcal{M}_l(\theta_l)-1]}{\partial\theta_j\partial\theta_k}\right|_{\boldsymbol{\theta}=\mathbf{0}}\left(\left.\frac{\partial\mathcal{V}_{t+1,l}(\boldsymbol{\theta})}{\partial\theta_i}\right|_{\boldsymbol{\theta}=\mathbf{0}} + \underbrace{\Delta\overline{x}_{t,i}\mathcal{V}_{t+1,l}(\mathbf{0}) - \Delta\overline{x}_{t,i}\mathcal{V}_{t+1,l}(\mathbf{0})}_{=0}\right)\\ &= \Delta\overline{x}_{t,i}\Delta_{m+s}\overline{\delta}_{t,jk} + \Delta_{m+s,i}\overline{\delta}_{t,ijk} + \Delta_{mds,i}\overline{\delta}_{t,ijk}\end{aligned}\tag{A-44}$$

where, using eq. (A-27),

$$\begin{aligned}\Delta_{m+s,i}\overline{\delta}_{t,ijk} &= -\Delta\overline{x}_{t,i}\sum_{l=1}^{n_x}\left.\frac{\partial^2[\mathcal{M}_l(\theta_l)-1]}{\partial\theta_j\partial\theta_k}\right|_{\boldsymbol{\theta}=\mathbf{0}}\overline{\mu}_{t,l}+\sum_{l=1}^{n_x}\left.\frac{\partial^2[\mathcal{M}_l(\theta_l)-1]}{\partial\theta_j\partial\theta_k}\right|_{\boldsymbol{\theta}=\mathbf{0}}\left.\frac{\partial\mathcal{V}_{t+1,l}(\boldsymbol{\theta})}{\partial\theta_i}\right|_{\boldsymbol{\theta}=\mathbf{0}}\\ &=\sum_{l=1}^{n_x}\left.\frac{\partial^2[\mathcal{M}_l(\theta_l)-1]}{\partial\theta_j\partial\theta_k}\right|_{\boldsymbol{\theta}=\mathbf{0}}\left(-\frac{1}{\overline{w}_t}\mathrm{Cov}_t\big[w_t,\delta_{t,i}\big]\overline{\mu}_{t,l}+\left.\frac{\partial\mathcal{V}_{t+1,l}(\boldsymbol{\theta})}{\partial\theta_i}\right|_{\boldsymbol{\theta}=\mathbf{0}}\right),\end{aligned}\tag{A-45}$$

and where

$$\Delta_{mds,i}\overline{\delta}_{t,ijk} = -\Delta\overline{x}_{t,i}\frac{1}{\overline{w}_t}\sum_{l=1}^{n_x}\left.\frac{\partial^2[\mathcal{M}_l(\theta_l)-1]}{\partial\theta_j\partial\theta_k}\right|_{\boldsymbol{\theta}=\mathbf{0}}\mathrm{Cov}_t\big[w_t,\mu_l\big].\tag{A-46}$$

Note that here, the subscript "mds" refers to the fact that this term contains the change in (co)skewness due to mutation (the "m" part) coupled to change in the average (i.e., due to selection; the "ds" part).

Using eq. (A-26), we have

$$\begin{aligned}\left.\frac{\partial\mathcal{V}_{t+1,j}(\boldsymbol{\theta})}{\partial\theta_i}\right|_{\boldsymbol{\theta}=\mathbf{0}} &= \frac{1}{\overline{w}_t}\mathbb{E}_t[w_t\mu_j\delta_{t,i}]\\ &=\frac{1}{\overline{w}_t}\mathrm{Cos}_t\big[w_t,\mu_j,\delta_{t,i}\big]+\frac{1}{\overline{w}_t}\mathbb{E}_t[w_t\delta_{t,i}]\,\overline{\mu}_{t,j}+\mathbb{E}_t[\mu_j\delta_{t,i}]\\ &=\frac{1}{\overline{w}_t}\mathrm{Cos}_t\big[w_t,\mu_j,\delta_{t,i}\big]+\frac{1}{\overline{w}_t}\mathrm{Cov}_t\big[w_t,\delta_{t,i}\big]\overline{\mu}_{t,j}+\mathrm{Cov}_t\big[\mu_j,\delta_{t,i}\big],\end{aligned}\tag{A-47}$$

where

$$\begin{aligned}\mathrm{Cos}_t\big[a_t,b_t,c_t\big] &= \mathbb{E}_t\big[(a_t(\boldsymbol{x})-\overline{a}_t)(b_t(\boldsymbol{x})-\overline{b}_t)(c_t(\boldsymbol{x})-\overline{c}_t)\big]\\ &=\mathbb{E}_t[a_tb_tc_t]-\mathbb{E}_t[a_tb_t]\,\overline{c_t}-\mathbb{E}_t[a_tc_t]\,\overline{b_t}-\mathbb{E}_t[b_tc_t]\,\overline{a_t}+2\overline{a}_t\overline{b}_t\overline{c}_t\end{aligned}\tag{A-48}$$

denotes the coskewness between three functions $a_t$, $b_t$ and $c_t$ under the distribution of trait values at generation $t$. Thus, since $\partial^2\mathcal{M}_i(\theta_i)/\partial\theta_i\partial\theta_i|_{\boldsymbol{\theta}=\mathbf{0}}=\overline{\zeta}_{ii}$ (see eq. A-18) and $\partial^2\mathcal{M}_k(\theta_k)/\partial\theta_i\partial\theta_j|_{\boldsymbol{\theta}=\mathbf{0}}=0$ whenever $i\neq j$, $i\neq k$ or $j\neq k$, substituting (A-47) into eq. (A-45) gives

$$\Delta_{m+s,i}\overline{\delta}_{t,ijk}=\Delta_{m,i}\overline{\delta}_{t,ijk}+\Delta_{m\times s,i}\overline{\delta}_{t,ijk}\tag{A-49}$$

where

$$\Delta_{m,i}\overline{\delta}_{t,ijk}=\begin{cases}\overline{\zeta}_{jj}\mathrm{Cov}_t\big[\mu_j,\delta_{t,i}\big] & \text{if } j=k,\\ 0 & \text{if } j\neq k,\end{cases}\tag{A-50}$$

and

$$\Delta_{m\times s,i}\overline{\delta}_{t,ijk}=\begin{cases}\overline{\zeta}_{jj}\dfrac{1}{\overline{w}_t}\mathrm{Cos}_t\big[w_t,\mu_j,\delta_{t,i}\big] & \text{if } j=k,\\ 0 & \text{if } j\neq k.\end{cases}\tag{A-51}$$

For the same reasons, eq. (A-46) becomes

$$\Delta_{mds,i}\overline{\delta}_{t,ijk}=\begin{cases}-\Delta\overline{x}_{t,i}\overline{\zeta}_{jj}\dfrac{1}{\overline{w}_t}\mathrm{Cov}_t\big[w_t,\mu_j\big] & \text{if } j=k,\\ 0 & \text{if } j\neq k.\end{cases}\tag{A-52}$$

Gathering terms and simplifying, eq. (A-40) becomes

$$\Delta\overline{\delta}_{t,ijk} = \Delta_s\overline{\delta}_{t,ijk} + \Delta_{s\times s}\overline{\delta}_{t,ijk} + \Delta_{s\times s\times s}\overline{\delta}_{t,ijk} + \Delta_m\overline{\delta}_{t,ijk} + \Delta_{m\times s}\overline{\delta}_{t,ijk} + \Delta_{mds}\overline{\delta}_{t,ijk} \tag{A-53}$$

where

$$\Delta_s\overline{\delta}_{t,ijk} = \frac{1}{\overline{w}_t}\mathrm{Cov}_t\big[w_t, \delta_{t,i}\delta_{t,j}\delta_{t,k}\big] - \Delta\overline{x}_{t,i}\overline{\delta}_{t,jk} - \Delta\overline{x}_{t,j}\overline{\delta}_{t,ik} - \Delta\overline{x}_{t,k}\overline{\delta}_{t,ij}, \tag{A-54}$$

$$\Delta_{s\times s}\overline{\delta}_{t,ijk} = -\Delta\overline{x}_{t,i}\Delta_s\overline{\delta}_{t,jk} - \Delta\overline{x}_{t,j}\Delta_s\overline{\delta}_{t,ik} - \Delta\overline{x}_{t,k}\Delta_s\overline{\delta}_{t,ij}, \tag{A-55}$$

$$\Delta_{s\times s\times s}\overline{\delta}_{t,ijk} = 2\Delta\overline{x}_{t,i}\Delta\overline{x}_{t,j}\Delta\overline{x}_{t,k}, \tag{A-56}$$

and where

$$\begin{aligned}\Delta_m\overline{\delta}_{t,ijk} &= \Delta_{m,i}\overline{\delta}_{t,ijk} + \Delta_{m,j}\overline{\delta}_{t,ijk} + \Delta_{m,k}\overline{\delta}_{t,ijk},\\ \Delta_{m\times s}\overline{\delta}_{t,ijk} &= \Delta_{m\times s,i}\overline{\delta}_{t,ijk} + \Delta_{m\times s,j}\overline{\delta}_{t,ijk} + \Delta_{m\times s,k}\overline{\delta}_{t,ijk},\\ \Delta_{mds}\overline{\delta}_{t,ijk} &= \Delta_{mds,i}\overline{\delta}_{t,ijk} + \Delta_{mds,j}\overline{\delta}_{t,ijk} + \Delta_{mds,k}\overline{\delta}_{t,ijk}.\end{aligned} \tag{A-57}$$

Note that the terms $\Delta_{s\times s\times s}\overline{\delta}_{t,ijk}$ and $\Delta_{mds}\overline{\delta}_{t,ijk}$ contain only high-order terms, and will always be negligible to leading orders (this is shown in the next sections).

### Appendix B - Approximation and closure

In this appendix, we describe the approach to close the system of moments of order up to three in terms of these moments themselves.

#### B.1 Taylor expansions

We start by writing Taylor expansions for all the functions of interest appearing in appendix A, with the aim of separating terms of different orders in trait deviations. Throughout, we write $O(\epsilon^k)$ for remainders of $k^{\text{th}}$-order in the largest trait deviation.

##### B.1.1 Individual fitness

We start by expanding the $k^{\text{th}}$ regulating kernel $g_{t,k}(\boldsymbol{x},\boldsymbol{y})$ around $g_{t,k}(\boldsymbol{x},\overline{\boldsymbol{x}}_t)$

$$\begin{aligned} g_{t,k}(\boldsymbol{x},\boldsymbol{y}) &= g_{t,k}(\boldsymbol{x},\overline{\boldsymbol{x}}_t) + \sum_{i=1}^{n_x} \delta'_{t,i} \left.\frac{\partial g_{t,k}(\boldsymbol{x},\boldsymbol{y})}{\partial y_i}\right|_{\boldsymbol{y}=\overline{\boldsymbol{x}}_t} \\ &+ \frac{1}{2}\sum_{i=1}^{n_x}\sum_{j=1}^{n_x} \delta'_{t,i}\delta'_{t,j} \left.\frac{\partial^2 g_{t,k}(\boldsymbol{x},\boldsymbol{y})}{\partial y_i \partial y_j}\right|_{\boldsymbol{y}=\overline{\boldsymbol{x}}_t} + O(\epsilon^3), \end{aligned} \tag{B-58}$$

where $\delta'_{t,i} = y_i - \overline{x}_{t,i}$. Injecting this into eq. (1) gives

$$\phi_{t,k}(\boldsymbol{x}) = \overline{\phi}_{t,k}(\boldsymbol{x}) + \frac{1}{2}\sum_{i=1}^{n_x}\sum_{j=1}^{n_x} \overline{\delta}_{t,ij} \left.\frac{\partial^2 g_{t,k}(\boldsymbol{x},\boldsymbol{y})}{\partial y_i \partial y_j}\right|_{\boldsymbol{y}=\overline{\boldsymbol{x}}_t} + O(\epsilon^3), \tag{B-59}$$

where $\overline{\phi}_{t,k}(\boldsymbol{x}) = g_{t,k}(\boldsymbol{x},\overline{\boldsymbol{x}}_t)$. With this, individual fitness can be expanded around the vector $\overline{\boldsymbol{\phi}}_t(\boldsymbol{x}) = \boldsymbol{g}_t(\boldsymbol{x},\overline{\boldsymbol{x}}_t) = (g_{t,1}(\boldsymbol{x},\overline{\boldsymbol{x}}_t),\dots,g_{t,n_\phi}(\boldsymbol{x},\overline{\boldsymbol{x}}_t))$,

$$w_t(\boldsymbol{x},\boldsymbol{\phi}_t(\boldsymbol{x})) = w_t(\boldsymbol{x},\overline{\boldsymbol{\phi}}_t(\boldsymbol{x})) + \omega_t(\boldsymbol{x}) + O(\epsilon^3), \tag{B-60}$$

where

$$\begin{aligned} \omega_t(\boldsymbol{x}) &= \sum_{i=1}^{n_\phi} (\phi_{t,i}(\boldsymbol{x}) - \overline{\phi}_{t,i}(\boldsymbol{x})) \left.\frac{\partial w_t(\boldsymbol{x},\boldsymbol{\phi}_t(\boldsymbol{x}))}{\partial \phi_{t,i}(\boldsymbol{x})}\right|_{\boldsymbol{\phi}_t(\boldsymbol{x})=\overline{\boldsymbol{\phi}}_t(\boldsymbol{x})} \\ &= \frac{1}{2}\sum_{i=1}^{n_\phi}\sum_{j=1}^{n_x}\sum_{k=1}^{n_x} \overline{\delta}_{t,jk} \left.\frac{\partial^2 g_{t,i}(\boldsymbol{x},\boldsymbol{y})}{\partial y_j \partial y_k}\right|_{\boldsymbol{y}=\overline{\boldsymbol{x}}_t} \left.\frac{\partial w_t(\boldsymbol{x},\boldsymbol{\phi}_t(\boldsymbol{x}))}{\partial \phi_{t,i}(\boldsymbol{x})}\right|_{\boldsymbol{\phi}_t(\boldsymbol{x})=\overline{\boldsymbol{\phi}}_t(\boldsymbol{x})} \end{aligned} \tag{B-61}$$

is a regulating term of second order in trait deviations. Using the above we can expand individual fitness around average traits,

$$w_t(\boldsymbol{x},\boldsymbol{\phi}_t(\boldsymbol{x})) = w_t^\circ + \sum_{i=1}^{n_x} \delta_{t,i} s^w_{t,i}(\overline{\boldsymbol{x}}_t) + \frac{1}{2}\sum_{i=1}^{n_x}\sum_{j=1}^{n_x} \delta_{t,i}\delta_{t,j} h^w_{t,ij}(\overline{\boldsymbol{x}}_t) + \omega_t(\overline{\boldsymbol{x}}_t) + O(\epsilon^3), \tag{B-62}$$

where $w_t^\circ = w_t(\overline{\boldsymbol{x}}_t, \boldsymbol{g}_t(\overline{\boldsymbol{x}}_t, \overline{\boldsymbol{x}}_t))$ and where $s^w_{t,i}(\overline{\boldsymbol{x}}_t)$ and $h^w_{t,ij}(\overline{\boldsymbol{x}}_t)$ are given by eqs. (12) and (14), respectively.

According to eq. (B-62), the average fitness is

$$\overline{w}_t = w_t^\circ + \frac{1}{2}\sum_{i=1}^{n_x}\sum_{j=1}^{n_x}\overline{\delta}_{t,ij}h^w_{t,ij}(\overline{\boldsymbol{x}}_t) + \omega_t(\overline{\boldsymbol{x}}_t) + O(\epsilon^3). \tag{B-63}$$

Using eq. (B-63), we can write the inverse of average fitness $1/\overline{w}_t$ as,

$$\frac{1}{\overline{w}_t} = \frac{1}{w_t^\circ}\frac{1}{\left(1+H\right)}, \tag{B-64}$$

where

$$H = \frac{1}{w_t^\circ}\left(\frac{1}{2}\sum_{i=1}^{n_x}\sum_{j=1}^{n_x}\overline{\delta}_{t,ij}h^w_{t,ij}(\overline{\boldsymbol{x}}_t) + \omega_t(\overline{\boldsymbol{x}}_t)\right) + O(\epsilon^3). \tag{B-65}$$

Applying the ordinary reciprocal (geometric) series, i.e., $(1+H)^{-1} = 1 - H + H^2 + O(H^3)$, gives

$$\frac{1}{\overline{w}_t} = \frac{1}{w_t^\circ} - \frac{1}{{w_t^\circ}^2}\left(\frac{1}{2}\sum_{i=1}^{n_x}\sum_{j=1}^{n_x}\overline{\delta}_{t,ij}h^w_{t,ij}(\overline{\boldsymbol{x}}_t) + \omega_t(\overline{\boldsymbol{x}}_t)\right) + O(\epsilon^3). \tag{B-66}$$

##### B.1.2 Mutation probabilities

The mutation probability at locus $k$ is

$$\mu_k(\boldsymbol{x}) = \mu_{t,k}^\circ + \sum_{i=1}^{n_x}\delta_{t,i}s^{\mu_k}_{t,i}(\overline{\boldsymbol{x}}_t) + \frac{1}{2}\sum_{i=1}^{n_x}\sum_{j=1}^{n_x}\delta_{t,i}\delta_{t,j}h^{\mu_k}_{t,ij}(\overline{\boldsymbol{x}}_t) + O(\epsilon^3), \tag{B-67}$$

where $\mu_{t,k}^\circ = \mu_k(\overline{\boldsymbol{x}}_t)$. Accordingly, the average mutation probability at locus $k$ is

$$\overline{\mu}_{t,k} = \mu_{t,k}^\circ + \frac{1}{2}\sum_{i=1}^{n_x}\sum_{j=1}^{n_x}\overline{\delta}_{t,ij}h^{\mu_k}_{t,ij}(\overline{\boldsymbol{x}}_t) + O(\epsilon^3). \tag{B-68}$$

##### B.1.3 (Co)variances and (Co)skewnesses

Using the expansions derived above, we now obtain expansions for the (co)variances and (co)skewnesses between trait, fitness and mutation involved in trait change. We give each expression up to an order that will be sufficient when closing the system of moments in the next section. In doing so, we will need the fourth- and fifth-order trait moments,

$$\overline{\delta}_{t,ijkl} = \int \delta_{t,i}\delta_{t,j}\delta_{t,k}\delta_{t,l}p_t(\boldsymbol{x})\mathrm{d}\boldsymbol{x}, \qquad \overline{\delta}_{t,ijklm} = \int \delta_{t,i}\delta_{t,j}\delta_{t,k}\delta_{t,l}\delta_{t,m}p_t(\boldsymbol{x})\mathrm{d}\boldsymbol{x}. \tag{B-69}$$

Using eqs. (A-26), (B-62) and (B-63), we have

$$\begin{aligned}\mathrm{Cov}_t\big[w_t, \delta_{t,i}\big] &= \mathbb{E}_t[w_t \delta_{t,i}] \\ &= \sum_{j=1}^{n_x} \overline{\delta}_{t,ij} s^w_{t,j}(\overline{\boldsymbol{x}}_t) + \frac{1}{2}\sum_{j=1}^{n_x}\sum_{k=1}^{n_x} \overline{\delta}_{t,ijk} h^w_{t,jk}(\overline{\boldsymbol{x}}_t) + O(\epsilon^4).\end{aligned} \tag{B-70}$$

Similarly, we have

$$\begin{aligned}\mathrm{Cov}_t\big[w_t, \delta_{t,i}\delta_{t,j}\big] &= \mathbb{E}_t[w_t \delta_{t,i}\delta_{t,j}] - \overline{w}_t\overline{\delta}_{t,ij} \\ &= \sum_{k=1}^{n_x} \overline{\delta}_{t,ijk} s^w_{t,k}(\overline{\boldsymbol{x}}_t) + \frac{1}{2}\sum_{k=1}^{n_x}\sum_{l=1}^{n_x} \left(\overline{\delta}_{t,ijkl} - \overline{\delta}_{t,ij}\overline{\delta}_{t,kl}\right) h^w_{t,kl}(\overline{\boldsymbol{x}}_t) + O(\epsilon^5),\end{aligned} \tag{B-71}$$

and

$$\begin{aligned}\mathrm{Cov}_t\big[w_t, \delta_{t,i}\delta_{t,j}\delta_{t,k}\big] &= \mathbb{E}_t[w_t \delta_{t,i}\delta_{t,j}\delta_{t,k}] - \overline{w}_t\overline{\delta}_{t,ijk} \\ &= \sum_{l=1}^{n_x} \overline{\delta}_{t,ijkl} s^w_{t,l}(\overline{\boldsymbol{x}}_t) + \frac{1}{2}\sum_{l=1}^{n_x}\sum_{m=1}^{n_x} \left(\overline{\delta}_{t,ijklm} - \overline{\delta}_{t,ijk}\overline{\delta}_{t,lm}\right) h^w_{t,lm}(\overline{\boldsymbol{x}}_t) + O(\epsilon^6).\end{aligned} \tag{B-72}$$

Using eqs. (B-62) and (B-67), we also have

$$\mathrm{Cov}_t\big[w_t, \mu_i\big] = \sum_{j=1}^{n_x}\sum_{k=1}^{n_x} \overline{\delta}_{t,jk} s^w_{t,j}(\overline{\boldsymbol{x}}_t) s^{\mu_i}_{t,k}(\overline{\boldsymbol{x}}_t) + O(\epsilon^3), \tag{B-73}$$

$$\mathrm{Cos}_t\big[w_t, \mu_j, \delta_{t,i}\big] = \sum_{k=1}^{n_x}\sum_{l=1}^{n_x} \overline{\delta}_{t,ikl} s^w_{t,k}(\overline{\boldsymbol{x}}_t) s^{\mu_j}_{t,l}(\overline{\boldsymbol{x}}_t) + O(\epsilon^4), \tag{B-74}$$

and

$$\mathrm{Cov}_t\big[\mu_j, \delta_{t,i}\big] = \sum_{k=1}^{n_x} \overline{\delta}_{t,ik} s^{\mu_j}_{t,k}(\overline{\boldsymbol{x}}_t) + \frac{1}{2}\sum_{k=1}^{n_x}\sum_{l=1}^{n_x} \overline{\delta}_{t,ikl} h^{\mu_j}_{t,kl}(\overline{\boldsymbol{x}}_t) + O(\epsilon^4). \tag{B-75}$$

#### B.2 Cumulant-neglect closure

We now show how to approximate fourth- and fifth-order trait moments, appearing in the small-variance approximations above, in terms of lower-order moments.

Recall that $\mathcal{G}_t(\boldsymbol{\theta}) = \mathbb{E}_t\left[e^{\boldsymbol{\theta}\cdot\boldsymbol{\delta}_t}\right]$ is the centred moment-generating function of the trait distribution (eq. A-1) and define the corresponding cumulant-generating function $\mathcal{K}_t(\boldsymbol{\theta}) = \log \mathcal{G}_t(\boldsymbol{\theta})$, such that $\mathcal{G}_t(\boldsymbol{\theta}) = e^{\mathcal{K}_t(\boldsymbol{\theta})}$. The derivatives of $\mathcal{K}_t(\boldsymbol{\theta})$ give the trait cumulants, another set of statistics describing the same distribution on a logarithmic scale. By definition, they are given by

$$\begin{gathered}\left.\frac{\partial \mathcal{K}_t(\boldsymbol{\theta})}{\partial \theta_i}\right|_{\boldsymbol{\theta}=\mathbf{0}} = \overline{\kappa}_{t,i} \qquad \left.\frac{\partial^2 \mathcal{K}_t(\boldsymbol{\theta})}{\partial \theta_i \partial \theta_j}\right|_{\boldsymbol{\theta}=\mathbf{0}} = \overline{\kappa}_{t,ij} \qquad \left.\frac{\partial^3 \mathcal{K}_t(\boldsymbol{\theta})}{\partial \theta_i \partial \theta_j \partial \theta_k}\right|_{\boldsymbol{\theta}=\mathbf{0}} = \overline{\kappa}_{t,ijk}, \\ \left.\frac{\partial^4 \mathcal{K}_t(\boldsymbol{\theta})}{\partial \theta_i \partial \theta_j \partial \theta_k \partial \theta_l}\right|_{\boldsymbol{\theta}=\mathbf{0}} = \overline{\kappa}_{t,ijkl}, \qquad \left.\frac{\partial^5 \mathcal{K}_t(\boldsymbol{\theta})}{\partial \theta_i \partial \theta_j \partial \theta_k \partial \theta_l \partial \theta_m}\right|_{\boldsymbol{\theta}=\mathbf{0}} = \overline{\kappa}_{t,ijklm}.\end{gathered} \tag{B-76}$$

Expanding the exponential, we have the general relationship

$$\mathcal{G}_t(\boldsymbol{\theta}) = e^{\mathcal{K}_t(\boldsymbol{\theta})} = 1 + \sum_{k=1}^{\infty} \frac{1}{k!}\mathcal{K}_t(\boldsymbol{\theta})^k. \tag{B-77}$$

This relationship can be used to express trait moments in terms of cumulants. Since $\mathcal{K}_t(\mathbf{0}) = 0$, $n^{\text{th}}$-order derivatives of $\mathcal{K}_t(\boldsymbol{\theta})^k$ will vanish for $n < k$. Accordingly, the first-order centred moment is

$$\bar{\delta}_{t,i} = \left.\frac{\partial \mathcal{G}_t(\boldsymbol{\theta})}{\partial \theta_i}\right|_{\boldsymbol{\theta}=\mathbf{0}} = \left.\frac{\partial \mathcal{K}_t(\boldsymbol{\theta})}{\partial \theta_i}\right|_{\boldsymbol{\theta}=\mathbf{0}} = \bar{\kappa}_{t,i} = 0. \tag{B-78}$$

Using the chain rule, the second-order centred moment is

$$\begin{aligned}
\bar{\delta}_{t,ij} &= \left.\frac{\partial^2 \mathcal{G}_t(\boldsymbol{\theta})}{\partial \theta_i \partial \theta_j}\right|_{\boldsymbol{\theta}=\mathbf{0}} \\
&= \left.\frac{\partial \mathcal{K}_t(\boldsymbol{\theta})}{\partial \theta_i}\right|_{\boldsymbol{\theta}=\mathbf{0}} \left.\frac{\partial \mathcal{K}_t(\boldsymbol{\theta})}{\partial \theta_j}\right|_{\boldsymbol{\theta}=\mathbf{0}} + \left.\frac{\partial^2 \mathcal{K}_t(\boldsymbol{\theta})}{\partial \theta_i \partial \theta_j}\right|_{\boldsymbol{\theta}=\mathbf{0}} \\
&= \bar{\kappa}_{t,i}\bar{\kappa}_{t,j} + \bar{\kappa}_{t,ij} \\
&= \bar{\kappa}_{t,ij},
\end{aligned} \tag{B-79}$$

and the third-order centred moment is

$$\begin{aligned}
\bar{\delta}_{t,ijk} &= \left.\frac{\partial^3 \mathcal{G}_t(\boldsymbol{\theta})}{\partial \theta_i \partial \theta_j \partial \theta_k}\right|_{\boldsymbol{\theta}=\mathbf{0}} \\
&= \left.\frac{\partial \mathcal{K}_t(\boldsymbol{\theta})}{\partial \theta_i}\right|_{\boldsymbol{\theta}=\mathbf{0}} \left.\frac{\partial \mathcal{K}_t(\boldsymbol{\theta})}{\partial \theta_j}\right|_{\boldsymbol{\theta}=\mathbf{0}} \left.\frac{\partial \mathcal{K}_t(\boldsymbol{\theta})}{\partial \theta_k}\right|_{\boldsymbol{\theta}=\mathbf{0}} + \left.\frac{\partial^2 \mathcal{K}_t(\boldsymbol{\theta})}{\partial \theta_i \partial \theta_j}\right|_{\boldsymbol{\theta}=\mathbf{0}} \left.\frac{\partial \mathcal{K}_t(\boldsymbol{\theta})}{\partial \theta_k}\right|_{\boldsymbol{\theta}=\mathbf{0}} \\
&\quad + \left.\frac{\partial^2 \mathcal{K}_t(\boldsymbol{\theta})}{\partial \theta_i \partial \theta_k}\right|_{\boldsymbol{\theta}=\mathbf{0}} \left.\frac{\partial \mathcal{K}_t(\boldsymbol{\theta})}{\partial \theta_j}\right|_{\boldsymbol{\theta}=\mathbf{0}} + \left.\frac{\partial^2 \mathcal{K}_t(\boldsymbol{\theta})}{\partial \theta_j \partial \theta_k}\right|_{\boldsymbol{\theta}=\mathbf{0}} \left.\frac{\partial \mathcal{K}_t(\boldsymbol{\theta})}{\partial \theta_i}\right|_{\boldsymbol{\theta}=\mathbf{0}} + \left.\frac{\partial^3 \mathcal{K}_t(\boldsymbol{\theta})}{\partial \theta_i \partial \theta_j \partial \theta_k}\right|_{\boldsymbol{\theta}=\mathbf{0}} \\
&= \bar{\kappa}_{t,i}\bar{\kappa}_{t,j}\bar{\kappa}_{t,k} + \bar{\kappa}_{t,ij}\bar{\kappa}_{t,k} + \bar{\kappa}_{t,ik}\bar{\kappa}_{t,j} + \bar{\kappa}_{t,jk}\bar{\kappa}_{t,i} + \bar{\kappa}_{t,ijk} \\
&= \bar{\kappa}_{t,ijk}.
\end{aligned} \tag{B-80}$$

Eqs. (B-78)–(B-80) show that first-, second- and third-order cumulants are equivalent to same-order moments. Applying the same approach, the fourth-order centred moment is

$$\begin{aligned}
\bar{\delta}_{t,ijkl} &= \left.\frac{\partial^4 \mathcal{G}_t(\boldsymbol{\theta})}{\partial \theta_i \partial \theta_j \partial \theta_k \partial \theta_l}\right|_{\boldsymbol{\theta}=\mathbf{0}} \\
&= \bar{\kappa}_{t,i}\bar{\kappa}_{t,j}\bar{\kappa}_{t,k}\bar{\kappa}_{t,l} + \bar{\kappa}_{t,ij}\bar{\kappa}_{t,k}\bar{\kappa}_{t,l} + \bar{\kappa}_{t,ik}\bar{\kappa}_{t,j}\bar{\kappa}_{t,l} + \bar{\kappa}_{t,il}\bar{\kappa}_{t,j}\bar{\kappa}_{t,k} + \bar{\kappa}_{t,jk}\bar{\kappa}_{t,i}\bar{\kappa}_{t,l} \\
&\quad + \bar{\kappa}_{t,jl}\bar{\kappa}_{t,i}\bar{\kappa}_{t,k} + \bar{\kappa}_{t,kl}\bar{\kappa}_{t,i}\bar{\kappa}_{t,j} + \bar{\kappa}_{t,ij}\bar{\kappa}_{t,kl} + \bar{\kappa}_{t,ik}\bar{\kappa}_{t,jl} + \bar{\kappa}_{t,il}\bar{\kappa}_{t,jk} + \bar{\kappa}_{t,ijk}\bar{\kappa}_{t,l} \\
&\quad + \bar{\kappa}_{t,ijl}\bar{\kappa}_{t,k} + \bar{\kappa}_{t,ikl}\bar{\kappa}_{t,j} + \bar{\kappa}_{t,jkl}\bar{\kappa}_{t,i} + \bar{\kappa}_{t,ijkl} \\
&= \bar{\kappa}_{t,ij}\bar{\kappa}_{t,kl} + \bar{\kappa}_{t,ik}\bar{\kappa}_{t,jl} + \bar{\kappa}_{t,il}\bar{\kappa}_{t,jk} + \bar{\kappa}_{t,ijkl}.
\end{aligned} \tag{B-81}$$

Finally, directly discarding terms with first-order cumulants as factors for brevity, the fifth-order centred moment is

$$\begin{aligned}\overline{\delta}_{t,ijklm} = \ & \left.\frac{\partial^5 \mathcal{G}_t(\boldsymbol{\theta})}{\partial\theta_i\partial\theta_j\partial\theta_k\partial\theta_l\partial\theta_m}\right|_{\boldsymbol{\theta}=\mathbf{0}} \\ & = \overline{\kappa}_{t,ijk}\overline{\kappa}_{t,lm} + \overline{\kappa}_{t,ijl}\overline{\kappa}_{t,km} + \overline{\kappa}_{t,ijm}\overline{\kappa}_{t,kl} + \overline{\kappa}_{t,ikl}\overline{\kappa}_{t,jm} + \overline{\kappa}_{t,ikm}\overline{\kappa}_{t,jl} + \overline{\kappa}_{t,ilm}\overline{\kappa}_{t,jk} \\ & \quad + \overline{\kappa}_{t,jkl}\overline{\kappa}_{t,im} + \overline{\kappa}_{t,jkm}\overline{\kappa}_{t,il} + \overline{\kappa}_{t,jlm}\overline{\kappa}_{t,ik} + \overline{\kappa}_{t,klm}\overline{\kappa}_{t,ij} + \overline{\kappa}_{t,ijklm}.\end{aligned} \tag{B-82}$$

Substituting eqs. (B-79) and (B-80) into eqs. (B-81) and (B-82) gives

$$\overline{\delta}_{t,ijkl} = \overline{\delta}_{t,ij}\overline{\delta}_{t,kl} + \overline{\delta}_{t,ik}\overline{\delta}_{t,jl} + \overline{\delta}_{t,il}\overline{\delta}_{t,jk} + \overline{\kappa}_{t,ijkl}, \tag{B-83}$$

and

$$\begin{aligned}\overline{\delta}_{t,ijklm} = \ & \overline{\delta}_{t,ijk}\overline{\delta}_{t,lm} + \overline{\delta}_{t,ijl}\overline{\delta}_{t,km} + \overline{\delta}_{t,ijm}\overline{\delta}_{t,kl} + \overline{\delta}_{t,ikl}\overline{\delta}_{t,jm} + \overline{\delta}_{t,ikm}\overline{\delta}_{t,jl} \\ & + \overline{\delta}_{t,ilm}\overline{\delta}_{t,jk} + \overline{\delta}_{t,jkl}\overline{\delta}_{t,im} + \overline{\delta}_{t,jkm}\overline{\delta}_{t,il} + \overline{\delta}_{t,jlm}\overline{\delta}_{t,ik} + \overline{\delta}_{t,klm}\overline{\delta}_{t,ij} + \overline{\kappa}_{t,ijklm}.\end{aligned} \tag{B-84}$$

We can see from eq. (B-84) that, under the small-variance assumption only, $\overline{\delta}_{t,ijkl}$ and $\overline{\delta}_{t,ijklm}$ contain terms of order four and five ($\overline{\kappa}_{t,ijkl}$ and $\overline{\kappa}_{t,ijklm}$, respectively), orders that we want to include in downstream analyses.

The cumulant-neglect closure method consists in making additional assumptions about the shape of the distribution. In particular, we assume that

$$\overline{\kappa}_{t,ijkl} = O(\epsilon^6), \qquad \overline{\kappa}_{t,ijklm} = O(\epsilon^6). \tag{B-85}$$

Under this assumption and for all purposes here, eqs. (B-83) and (B-84) become

$$\overline{\delta}_{t,ijkl} = \overline{\delta}_{t,ij}\overline{\delta}_{t,kl} + \overline{\delta}_{t,ik}\overline{\delta}_{t,jl} + \overline{\delta}_{t,il}\overline{\delta}_{t,jk}, \tag{B-86}$$

and

$$\begin{aligned}\overline{\delta}_{t,ijklm} = \ & \overline{\delta}_{t,ijk}\overline{\delta}_{t,lm} + \overline{\delta}_{t,ijl}\overline{\delta}_{t,km} + \overline{\delta}_{t,ijm}\overline{\delta}_{t,kl} + \overline{\delta}_{t,ikl}\overline{\delta}_{t,jm} + \overline{\delta}_{t,ikm}\overline{\delta}_{t,jl} \\ & + \overline{\delta}_{t,ilm}\overline{\delta}_{t,jk} + \overline{\delta}_{t,jkl}\overline{\delta}_{t,im} + \overline{\delta}_{t,jkm}\overline{\delta}_{t,il} + \overline{\delta}_{t,jlm}\overline{\delta}_{t,ik} + \overline{\delta}_{t,klm}\overline{\delta}_{t,ij},\end{aligned} \tag{B-87}$$

respectively.

#### B.3 Closed components of change

In this subsection, we combine the results obtained in previous subsections to express components of change in moments as fully closed expressions. For each component of change in average trait, (co)variances and (co)skewnesses, we consider only terms up to the order of second-order selection terms (that is order three, four, and five, respectively). Under our assumptions, higher-order terms do not play a significant role as long as first- and second-order selection coefficients are never zero at the same time. In sorting orders, recall that we assume that parent-offspring differences are of the same size as deviations from the means, i.e., that $\zeta_i = O(\epsilon)$, and that mutation probabilities may be as large as one, $\mu_i(\boldsymbol{x}) = O(1)$.

###### B.3.1 Average traits

Using eqs. (A-27), (B-66) and (B-70), we have

$$\Delta \overline{x}_{t,i} = \underbrace{\frac{1}{w_t^\circ} \sum_{j=1}^{n_x} \overline{\delta}_{t,ij} s_{t,j}^w(\overline{\boldsymbol{x}}_t)}_{\Delta_s^1 \overline{x}_{t,i}} + \underbrace{\frac{1}{2w_t^\circ} \sum_{j=1}^{n_x} \sum_{k=1}^{n_x} \overline{\delta}_{t,ijk} h_{t,jk}^w(\overline{\boldsymbol{x}}_t)}_{\Delta_s^2 \overline{x}_{t,i}} + O(\epsilon^4). \tag{B-88}$$

This recovers eq. (10) of the main text.

###### B.3.2 Trait (co)variances

Using eqs. (A-32), (B-66), (B-71) and (B-86), we have

$$\Delta_s \overline{\delta}_{t,ij} = \underbrace{\frac{1}{w_t^\circ} \sum_{k=1}^{n_x} \overline{\delta}_{t,ijk} s_{t,k}^w(\overline{\boldsymbol{x}}_t)}_{\Delta_s^1 \overline{\delta}_{t,ij}} + \underbrace{\frac{1}{2w_t^\circ} \sum_{k=1}^{n_x} \sum_{l=1}^{n_x} \left( \overline{\delta}_{t,ik} \overline{\delta}_{t,jl} + \overline{\delta}_{t,il} \overline{\delta}_{t,jk} \right) h_{t,kl}^w(\overline{\boldsymbol{x}}_t)}_{\Delta_s^2 \overline{\delta}_{t,ij}} + O(\epsilon^5), \tag{B-89}$$

where one term of the closure approximation of $\overline{\delta}_{t,ijkl}$ exactly cancels the product of variances in eq. (B-71). This recovers eqs. (16) and (17) of the main text.

Using eqs. (A-37) and (B-88), retaining only the leading term of each factor since the product of any two next-order terms exceeds the target order, we have

$$\Delta_{s \times s} \overline{\delta}_{t,ij} = \underbrace{- \frac{1}{w_t^{\circ 2}} \sum_{k=1}^{n_x} \sum_{l=1}^{n_x} \overline{\delta}_{t,ik} \overline{\delta}_{t,jl} s_{t,k}^w(\overline{\boldsymbol{x}}_t) s_{t,l}^w(\overline{\boldsymbol{x}}_t)}_{\Delta_{s \times s}^1 \overline{\delta}_{t,ij}} + O(\epsilon^5). \tag{B-90}$$

This recovers eq. (18) of the main text.

Using eq. (A-35) and eq. (B-68), we have

$$\Delta_m \overline{\delta}_{t,ij} = \begin{cases} \overline{\zeta}_{ii} \left( \mu_{t,i}^\circ + \dfrac{1}{2} \displaystyle\sum_{k=1}^{n_x} \sum_{l=1}^{n_x} \overline{\delta}_{t,kl} h_{t,kl}^{\mu_i}(\overline{\boldsymbol{x}}_t) \right) + O(\epsilon^5) & \text{if } i = j, \\ 0 & \text{if } i \neq j. \end{cases} \tag{B-91}$$

Under our leading-orders truncation, this recovers eq. (19) of the main text.

Using eqs. (A-36), (B-66) and (B-73), keeping only the leading term of $\mathrm{Cov}_t\big[w_t, \mu_i\big]$ since $\overline{\zeta}_{ii}$ is already of order two, we have

$$\Delta_{m\times s}\overline{\delta}_{t,ij} = \begin{cases} \dfrac{\overline{\zeta}_{ii}}{w_t^\circ}\displaystyle\sum_{k=1}^{n_x}\sum_{l=1}^{n_x}\overline{\delta}_{t,kl}s_{t,k}^w(\overline{\boldsymbol{x}}_t)s_{t,l}^{\mu_i}(\overline{\boldsymbol{x}}_t) + O(\epsilon^5) & \text{if } i = j, \\ 0 & \text{if } i \neq j. \end{cases} \tag{B-92}$$

To leading orders, this recovers eq. (21) of the main text.

##### B.3.3 Trait (co)skewnesses

Injecting eqs. (B-66), (B-72) and (B-88) into eq. (A-54), and replacing the fourth- and fifth-order moments by their cumulant-neglect approximations (eqs. B-86 and B-87) gives

$$\begin{aligned}
\Delta_s\overline{\delta}_{t,ijk} = {} & \frac{1}{w_t^\circ}\sum_{l=1}^{n_x}\big(\overline{\delta}_{t,ij}\overline{\delta}_{t,kl} + \overline{\delta}_{t,ik}\overline{\delta}_{t,jl} + \overline{\delta}_{t,il}\overline{\delta}_{t,jk}\big)s_{t,l}^w(\overline{\boldsymbol{x}}_t) \\
& + \frac{1}{2w_t^\circ}\sum_{l=1}^{n_x}\sum_{m=1}^{n_x}\big(\overline{\delta}_{t,ijk}\overline{\delta}_{t,lm} + \overline{\delta}_{t,ijl}\overline{\delta}_{t,km} + \overline{\delta}_{t,ijm}\overline{\delta}_{t,kl} + \overline{\delta}_{t,ikl}\overline{\delta}_{t,jm} \\
& \quad + \overline{\delta}_{t,ikm}\overline{\delta}_{t,jl} + \overline{\delta}_{t,ilm}\overline{\delta}_{t,jk} + \overline{\delta}_{t,jkl}\overline{\delta}_{t,im} + \overline{\delta}_{t,jkm}\overline{\delta}_{t,il} \\
& \quad + \overline{\delta}_{t,jlm}\overline{\delta}_{t,ik} + \overline{\delta}_{t,klm}\overline{\delta}_{t,ij} - \overline{\delta}_{t,ijk}\overline{\delta}_{t,lm}\big)h_{t,lm}^w(\overline{\boldsymbol{x}}_t) \\
& - \frac{1}{w_t^\circ}\Big(\sum_{l=1}^{n_x}\overline{\delta}_{t,il}s_{t,l}^w(\overline{\boldsymbol{x}}_t) + \frac{1}{2}\sum_{l=1}^{n_x}\sum_{m=1}^{n_x}\overline{\delta}_{t,ilm}h_{t,lm}^w(\overline{\boldsymbol{x}}_t)\Big)\overline{\delta}_{t,jk} \\
& - \frac{1}{w_t^\circ}\Big(\sum_{l=1}^{n_x}\overline{\delta}_{t,jl}s_{t,l}^w(\overline{\boldsymbol{x}}_t) + \frac{1}{2}\sum_{l=1}^{n_x}\sum_{m=1}^{n_x}\overline{\delta}_{t,jlm}h_{t,lm}^w(\overline{\boldsymbol{x}}_t)\Big)\overline{\delta}_{t,ik} \\
& - \frac{1}{w_t^\circ}\Big(\sum_{l=1}^{n_x}\overline{\delta}_{t,kl}s_{t,l}^w(\overline{\boldsymbol{x}}_t) + \frac{1}{2}\sum_{l=1}^{n_x}\sum_{m=1}^{n_x}\overline{\delta}_{t,klm}h_{t,lm}^w(\overline{\boldsymbol{x}}_t)\Big)\overline{\delta}_{t,ij} + O(\epsilon^6).
\end{aligned} \tag{B-93}$$

Since the first-order selection terms from eq. (B-88) cancel out those from eq. (B-72), and the second-order selection terms from eq. (B-88) cancel out some of those from eq. (B-72), eq. (B-93) simplifies to

$$\Delta_s\overline{\delta}_{t,ijk} = \Delta_s^2\overline{\delta}_{t,ijk} + O(\epsilon^6), \tag{B-94}$$

with

$$\begin{aligned}
\Delta_s^2\overline{\delta}_{t,ijk} = {} & \frac{1}{2w_t^\circ}\sum_{l=1}^{n_x}\sum_{m=1}^{n_x}\big(\overline{\delta}_{t,il}\overline{\delta}_{t,jkm} + \overline{\delta}_{t,im}\overline{\delta}_{t,jkl} + \overline{\delta}_{t,jl}\overline{\delta}_{t,ikm} \\
& + \overline{\delta}_{t,jm}\overline{\delta}_{t,ikl} + \overline{\delta}_{t,kl}\overline{\delta}_{t,ijm} + \overline{\delta}_{t,km}\overline{\delta}_{t,ijl}\big)h_{t,lm}^w(\overline{\boldsymbol{x}}_t).
\end{aligned} \tag{B-95}$$

This recovers eq. (24) of the main text.

Next, using eqs. (A-55), (B-88) and (B-89), we have

$$\Delta_{s\times s}\overline{\delta}_{t,ijk} = \underbrace{-\frac{1}{{w_t^{\circ}}^2}\sum_{l=1}^{n_x}\sum_{m=1}^{n_x}\left(\overline{\delta}_{t,il}\overline{\delta}_{t,jkm}+\overline{\delta}_{t,jl}\overline{\delta}_{t,ikm}+\overline{\delta}_{t,kl}\overline{\delta}_{t,ijm}\right)s_{t,l}^{w}(\overline{\boldsymbol{x}}_t)s_{t,m}^{w}(\overline{\boldsymbol{x}}_t)}_{\Delta^1_{s\times s}\overline{\delta}_{t,ijk}}+O(\epsilon^6), \tag{B-96}$$

which recovers eq. (25) of the main text.

Using eqs. (A-56) and (B-88), we have

$$\Delta_{s\times s\times s}\overline{\delta}_{t,ijk} = O(\epsilon^6). \tag{B-97}$$

We thus ignore this higher-order term.

Using eqs. (A-50) and (B-75), we have

$$\Delta_{m,i}\overline{\delta}_{t,ijk} = \begin{cases} \overline{\zeta}_{jj}\Big(\sum_{l=1}^{n_x}\overline{\delta}_{t,il}s_{t,l}^{\mu_j}(\overline{\boldsymbol{x}}_t)+\frac{1}{2}\sum_{l=1}^{n_x}\sum_{m=1}^{n_x}\overline{\delta}_{t,ilm}h_{t,lm}^{\mu_j}(\overline{\boldsymbol{x}}_t)\Big)+O(\epsilon^6) & \text{if } j=k,\\ 0 & \text{if } j\neq k. \end{cases} \tag{B-98}$$

To leading orders, this recovers eq. (27) of the main text.

Using eqs. (A-51) and (B-74), we have

$$\Delta_{m\times s,i}\overline{\delta}_{t,ijk} = \begin{cases} \overline{\zeta}_{jj}\frac{1}{w_t^{\circ}}\sum_{l=1}^{n_x}\sum_{m=1}^{n_x}\overline{\delta}_{t,ilm}s_{t,l}^{w}(\overline{\boldsymbol{x}}_t)s_{t,m}^{\mu_j}(\overline{\boldsymbol{x}}_t)+O(\epsilon^6) & \text{if } j=k,\\ 0 & \text{if } j\neq k. \end{cases} \tag{B-99}$$

To leading orders, this recovers eq. (29) of the main text.

Finally, using eqs. (A-52) and (B-73), we have

$$\Delta_{mds,i}\overline{\delta}_{t,ijk} = O(\epsilon^6) \tag{B-100}$$

We thus ignore this higher-order term.

### Appendix C - Example model

This appendix contains results and additional descriptions related to our two-trait example model.

#### C.1 System of moments

Here we give the complete closed system of moments obtained under the assumptions of our example model. We use the fact that $h^w_{t,12} = 0$ according to our definition of fitness, but do not assume that $\alpha = 0$ as in the main text.

##### Means

To leading orders, the equations for the change in average traits are

$$\begin{aligned}
\Delta \overline{x}_{t,1} &= \frac{1}{w^\circ_t}\big(\overline{\delta}_{t,11} s^w_{t,1}(\overline{x}_{t,1}) + \overline{\delta}_{t,12} s^w_{t,2}(\overline{x}_{t,2}) + \frac{1}{2}\overline{\delta}_{t,111} h^w_{t,11}(\overline{x}_{t,1}) + \frac{1}{2}\overline{\delta}_{t,122} h^w_{t,22}(\overline{x}_{t,2})\big),\\
\Delta \overline{x}_{t,2} &= \frac{1}{w^\circ_t}\big(\overline{\delta}_{t,22} s^w_{t,2}(\overline{x}_{t,2}) + \overline{\delta}_{t,12} s^w_{t,1}(\overline{x}_{t,1}) + \frac{1}{2}\overline{\delta}_{t,222} h^w_{t,22}(\overline{x}_{t,2}) + \frac{1}{2}\overline{\delta}_{t,112} h^w_{t,11}(\overline{x}_{t,1})\big),
\end{aligned} \tag{C-101}$$

where

$$w^\circ_t = \frac{\lambda}{1+\rho N_t} e^{-\gamma(\overline{x}_{t,1}-z_t)^2} - c(\overline{x}_{t,2}), \tag{C-102}$$

##### Variances and covariances

To leading orders, the equations for the change in trait (co)variances are

$$\begin{aligned}
\Delta \overline{\delta}_{t,11} &= \frac{1}{w^\circ_t}\big(\overline{\delta}_{t,111} s^w_{t,1}(\overline{x}_{t,1}) + \overline{\delta}_{t,112} s^w_{t,2}(\overline{x}_{t,2})\big) + \overline{\delta}^2_{t,11}\Big(\frac{h^w_{t,11}(\overline{x}_{t,1})}{w^\circ_t} - \frac{s^w_{t,1}(\overline{x}_{t,1})^2}{w^{\circ 2}_t}\Big)\\
&\quad + \overline{\delta}^2_{t,12}\Big(\frac{h^w_{t,22}(\overline{x}_{t,2})}{w^\circ_t} - \frac{s^w_{t,2}(\overline{x}_{t,2})^2}{w^{\circ 2}_t}\Big) - \frac{2}{w^{\circ 2}_t}\overline{\delta}_{t,11}\overline{\delta}_{t,12} s^w_{t,1}(\overline{x}_{t,1}) s^w_{t,2}(\overline{x}_{t,2})\\
&\quad + \overline{\zeta}_{11}\Big(\mu_1(\overline{x}_{t,2}) + \frac{1}{2}\overline{\delta}_{t,22} h^{\mu_1}_{t,22}(\overline{x}_{t,2})\Big)\\
&\quad + \frac{1}{w^\circ_t}\overline{\zeta}_{11}\big(\overline{\delta}_{t,12} s^w_{t,1}(\overline{x}_{t,1}) + \overline{\delta}_{t,22} s^w_{t,2}(\overline{x}_{t,2})\big) s^{\mu_1}_{t,2}(\overline{x}_{t,2}),\\
\Delta \overline{\delta}_{t,22} &= \frac{1}{w^\circ_t}\big(\overline{\delta}_{t,222} s^w_{t,2}(\overline{x}_{t,2}) + \overline{\delta}_{t,122} s^w_{t,1}(\overline{x}_{t,1})\big) + \overline{\delta}^2_{t,22}\Big(\frac{h^w_{t,22}(\overline{x}_{t,2})}{w^\circ_t} - \frac{s^w_{t,2}(\overline{x}_{t,2})^2}{w^{\circ 2}_t}\Big)\\
&\quad + \overline{\delta}^2_{t,12}\Big(\frac{h^w_{t,11}(\overline{x}_{t,1})}{w^\circ_t} - \frac{s^w_{t,1}(\overline{x}_{t,1})^2}{w^{\circ 2}_t}\Big) - \frac{2}{w^{\circ 2}_t}\overline{\delta}_{t,22}\overline{\delta}_{t,12} s^w_{t,1}(\overline{x}_{t,1}) s^w_{t,2}(\overline{x}_{t,2})\\
&\quad + \overline{\zeta}_{22}\Big(\mu_2(\overline{x}_{t,2}) + \frac{1}{2}\overline{\delta}_{t,22} h^{\mu_2}_{t,22}(\overline{x}_{t,2})\Big)\\
&\quad + \frac{1}{w^\circ_t}\overline{\zeta}_{22}\big(\overline{\delta}_{t,22} s^w_{t,2}(\overline{x}_{t,2}) + \overline{\delta}_{t,12} s^w_{t,1}(\overline{x}_{t,1})\big) s^{\mu_2}_{t,2}(\overline{x}_{t,2}),\\
\Delta \overline{\delta}_{t,12} &= \frac{1}{w^\circ_t}\big(\overline{\delta}_{t,122} s^w_{t,2}(\overline{x}_{t,2}) + \overline{\delta}_{t,112} s^w_{t,1}(\overline{x}_{t,1})\big) + \overline{\delta}_{t,11}\overline{\delta}_{t,12}\Big(\frac{h^w_{t,11}(\overline{x}_{t,1})}{w^\circ_t} - \frac{s^w_{t,1}(\overline{x}_{t,1})^2}{w^{\circ 2}_t}\Big)\\
&\quad + \overline{\delta}_{t,12}\overline{\delta}_{t,22}\Big(\frac{h^w_{t,22}(\overline{x}_{t,2})}{w^\circ_t} - \frac{s^w_{t,2}(\overline{x}_{t,2})^2}{w^{\circ 2}_t}\Big)\\
&\quad - \frac{1}{w^{\circ 2}_t}\big(\overline{\delta}_{t,11}\overline{\delta}_{t,22} + \overline{\delta}^2_{t,12}\big) s^w_{t,1}(\overline{x}_{t,1}) s^w_{t,2}(\overline{x}_{t,2}).
\end{aligned} \tag{C-103}$$

**Skewnesses and coskewnesses**

To leading orders, the equations for the change in (co)skewnesses are

$$
\begin{aligned}
\Delta\overline{\delta}_{t,111} &= 3\overline{\delta}_{t,11}\overline{\delta}_{t,111}\Big(\frac{h^{w}_{t,11}(\overline{x}_{t,1})}{w^{\circ}_{t}} - \frac{s^{w}_{t,1}(\overline{x}_{t,1})^2}{w^{\circ 2}_{t}}\Big) + 3\overline{\delta}_{t,12}\overline{\delta}_{t,112}\Big(\frac{h^{w}_{t,22}(\overline{x}_{t,2})}{w^{\circ}_{t}} - \frac{s^{w}_{t,2}(\overline{x}_{t,2})^2}{w^{\circ 2}_{t}}\Big) \\
&\quad - \frac{3}{w^{\circ 2}_{t}}\big(\overline{\delta}_{t,11}\overline{\delta}_{t,112} + \overline{\delta}_{t,12}\overline{\delta}_{t,111}\big)s^{w}_{t,1}(\overline{x}_{t,1})s^{w}_{t,2}(\overline{x}_{t,2}) \\
&\quad + 3\overline{\zeta}_{11}\Big(\overline{\delta}_{t,12} + \frac{1}{w^{\circ}_{t}}\big(\overline{\delta}_{t,112}s^{w}_{t,1}(\overline{x}_{t,1}) + \overline{\delta}_{t,122}s^{w}_{t,2}(\overline{x}_{t,2})\big)\Big)s^{\mu_1}_{t,2}(\overline{x}_{t,2}) \\
&\quad + \frac{3}{2}\overline{\zeta}_{11}\overline{\delta}_{t,122}h^{\mu_1}_{t,22}(\overline{x}_{t,2}), \\
\Delta\overline{\delta}_{t,222} &= 3\overline{\delta}_{t,12}\overline{\delta}_{t,122}\Big(\frac{h^{w}_{t,11}(\overline{x}_{t,1})}{w^{\circ}_{t}} - \frac{s^{w}_{t,1}(\overline{x}_{t,1})^2}{w^{\circ 2}_{t}}\Big) + 3\overline{\delta}_{t,22}\overline{\delta}_{t,222}\Big(\frac{h^{w}_{t,22}(\overline{x}_{t,2})}{w^{\circ}_{t}} - \frac{s^{w}_{t,2}(\overline{x}_{t,2})^2}{w^{\circ 2}_{t}}\Big) \\
&\quad - \frac{3}{w^{\circ 2}_{t}}\big(\overline{\delta}_{t,12}\overline{\delta}_{t,222} + \overline{\delta}_{t,22}\overline{\delta}_{t,122}\big)s^{w}_{t,1}(\overline{x}_{t,1})s^{w}_{t,2}(\overline{x}_{t,2}) \\
&\quad + 3\overline{\zeta}_{22}\Big(\overline{\delta}_{t,22} + \frac{1}{w^{\circ}_{t}}\big(\overline{\delta}_{t,122}s^{w}_{t,1}(\overline{x}_{t,1}) + \overline{\delta}_{t,222}s^{w}_{t,2}(\overline{x}_{t,2})\big)\Big)s^{\mu_2}_{t,2}(\overline{x}_{t,2}) \\
&\quad + \frac{3}{2}\overline{\zeta}_{22}\overline{\delta}_{t,222}h^{\mu_2}_{t,22}(\overline{x}_{t,2}), \\
\Delta\overline{\delta}_{t,112} &= \big(2\overline{\delta}_{t,11}\overline{\delta}_{t,112} + \overline{\delta}_{t,12}\overline{\delta}_{t,111}\big)\Big(\frac{h^{w}_{t,11}(\overline{x}_{t,1})}{w^{\circ}_{t}} - \frac{s^{w}_{t,1}(\overline{x}_{t,1})^2}{w^{\circ 2}_{t}}\Big) \\
&\quad + \big(2\overline{\delta}_{t,12}\overline{\delta}_{t,122} + \overline{\delta}_{t,22}\overline{\delta}_{t,112}\big)\Big(\frac{h^{w}_{t,22}(\overline{x}_{t,2})}{w^{\circ}_{t}} - \frac{s^{w}_{t,2}(\overline{x}_{t,2})^2}{w^{\circ 2}_{t}}\Big) \\
&\quad - \frac{1}{w^{\circ 2}_{t}}\big(3\overline{\delta}_{t,12}\overline{\delta}_{t,112} + \overline{\delta}_{t,22}\overline{\delta}_{t,111} + 2\overline{\delta}_{t,11}\overline{\delta}_{t,122}\big)s^{w}_{t,1}(\overline{x}_{t,1})s^{w}_{t,2}(\overline{x}_{t,2}) \\
&\quad + \overline{\zeta}_{11}\Big(\overline{\delta}_{t,22} + \frac{1}{w^{\circ}_{t}}\big(\overline{\delta}_{t,122}s^{w}_{t,1}(\overline{x}_{t,1}) + \overline{\delta}_{t,222}s^{w}_{t,2}(\overline{x}_{t,2})\big)\Big)s^{\mu_1}_{t,2}(\overline{x}_{t,2}) \\
&\quad + \frac{1}{2}\overline{\zeta}_{11}\overline{\delta}_{t,222}h^{\mu_1}_{t,22}(\overline{x}_{t,2}), \\
\Delta\overline{\delta}_{t,122} &= \big(2\overline{\delta}_{t,12}\overline{\delta}_{t,112} + \overline{\delta}_{t,11}\overline{\delta}_{t,122}\big)\Big(\frac{h^{w}_{t,11}(\overline{x}_{t,1})}{w^{\circ}_{t}} - \frac{s^{w}_{t,1}(\overline{x}_{t,1})^2}{w^{\circ 2}_{t}}\Big) \\
&\quad + \big(2\overline{\delta}_{t,22}\overline{\delta}_{t,122} + \overline{\delta}_{t,12}\overline{\delta}_{t,222}\big)\Big(\frac{h^{w}_{t,22}(\overline{x}_{t,2})}{w^{\circ}_{t}} - \frac{s^{w}_{t,2}(\overline{x}_{t,2})^2}{w^{\circ 2}_{t}}\Big) \\
&\quad - \frac{1}{w^{\circ 2}_{t}}\big(3\overline{\delta}_{t,12}\overline{\delta}_{t,122} + \overline{\delta}_{t,11}\overline{\delta}_{t,222} + 2\overline{\delta}_{t,22}\overline{\delta}_{t,112}\big)s^{w}_{t,1}(\overline{x}_{t,1})s^{w}_{t,2}(\overline{x}_{t,2}) \\
&\quad + \overline{\zeta}_{22}\Big(\overline{\delta}_{t,12} + \frac{1}{w^{\circ}_{t}}\big(\overline{\delta}_{t,112}s^{w}_{t,1}(\overline{x}_{t,1}) + \overline{\delta}_{t,122}s^{w}_{t,2}(\overline{x}_{t,2})\big)\Big)s^{\mu_2}_{t,2}(\overline{x}_{t,2}) \\
&\quad + \frac{1}{2}\overline{\zeta}_{22}\overline{\delta}_{t,122}h^{\mu_2}_{t,22}(\overline{x}_{t,2}).
\end{aligned}
\tag{C-104}
$$

Eqs. (39)–(48) of the main text are obtained from eqs. (C-101)–(C-104) by setting $\alpha = 0$, such that $s^{w}_{t,2}(\overline{x}_{t,2}) = 0$ and $h^{w}_{t,22}(\overline{x}_{t,2}) = 0$, and by assuming that mutation probabilities are linear functions (i.e., $h^{\mu_1}_{t,22}(\overline{x}_{t,2}) = h^{\mu_2}_{t,22}(\overline{x}_{t,2}) = 0$).

**Population density**

The total density follows eq. (5), $N_{t+1} = \overline{w}_t N_t$. Using the second-order approximation of mean fitness (eq. B-63) and with $h^w_{t,12} = 0$, to leading orders we have

$$N_{t+1} = N_t\Big(w^\circ_t + \frac{1}{2}\big(\overline{\delta}_{t,11}h^w_{t,11}(\overline{x}_{t,1}) + \overline{\delta}_{t,22}h^w_{t,22}(\overline{x}_{t,2})\big) + \omega_t(\overline{x}_{t,1}) + O(\epsilon^3)\Big), \tag{C-105}$$

with

$$\omega_t(\overline{x}_{t,1}) = \eta\,\overline{\delta}_{t,11}\,\frac{\lambda\rho N_t}{(1+\rho N_t)^2}e^{-\gamma(\overline{x}_{t,1}-z_t)^2}. \tag{C-106}$$

#### C.2 Individual-based simulations

Here we describe the individual-based simulations of our example two-trait model used in the main text. A Rust program implementing these simulations is are available at zenodo.org (Supplementary archive 1; DOI: 10.5281/zenodo.24368121).

Each individual in the simulation model is characterised by a pair of real numbers, giving the values of the viability and modifier trait for that individual (i.e., its haplotype). We start all simulation runs by initiating a population of ten thousand individuals, whose viability and modifier trait values are drawn independently from normal distributions with the initial means given in the main text (or in the figure caption for fig. S1) and variance $10^{-4}$, such that the initial distribution matches that of the numerical iterations (up to sampling error). The population is then iterated through the following steps:

1. **Environmental value**. The current optimal viability trait value $z_t$ (see eq. 30) is decided. In the transient adaptation case, it is set to the same constant value in every generation (i.e., the new phenotypic optimum). The same holds in the strong-competition case (fig. S1), but the population starts directly at that value. In the fluctuating selection case, $z_t$ is set to one of two values, depending on its previous value and on the number of generations elapsed since the start of the run. If that number is not a multiple of the environmental half-period $T$, $z_t$ is set to the same value as in the previous generation. If it is a multiple of $T$, $z_t$ is set to the other value.

2. **Reproduction**. Individuals each produce a random number of offspring. This number is drawn from a Poisson distribution with rate equal to the fitness of the parent individual (truncated at zero), itself computed from that parent's trait values according to eq. (32). The value of the regulating factor $\phi_{t,1}(x_1)$, for a parent with viability trait $x_1$, is calculated as

$$\phi_{t,1}(x_1) = \frac{1}{|\mathcal{N}|}\sum_{y_1\in\mathcal{N}} g_1(x_1, y_1), \tag{C-107}$$

   where $\mathcal{N}$ is a set of $|\mathcal{N}| = 30$ viability trait values randomly sampled in the population without replacement, drawn once per generation and shared by all individuals. The sample size of 30 follows the usual rule of thumb for obtaining a stable estimate of a mean, and we verified that results are insensitive to this choice.

3. **Mutation**. Each offspring acquires mutations depending on its parent's trait values. At each locus, the mutation probability is calculated according to eq. (31). When a mutation occurs at locus $i \in \{1, 2\}$, the offspring trait value is obtained as the sum of its parent's trait value and of a random mutation effect drawn from a normal distribution with mean zero and variance $\overline{\zeta}_{ii} = 10^{-4}$ (except for runs reported in fig. S3, where $\overline{\zeta}_{22} = 10^{-3}$).

4. **Replacement**. All parents are removed and offspring become the next generation.
